# Slow synaptic plasticity from the hippocampus underlies gradual mapping and fragmentation of novel spaces by grid cells

**DOI:** 10.1101/2025.07.30.667696

**Authors:** Lujia Chen, Ling Liang Dong, Hoon Shin, Farid Shahid, Taylor Joseph Malone, Yan Ma, Sreerag Othayoth Vasu, Nai-Wen Tien, Kyle Cekada, Kevin Jiang Zhang, Lucy Anderson, Sarthak Chandra, Ila Fiete, Veronica A. Alvarez, Yi Gu

**Affiliations:** Spatial Navigation and Memory Unit, National Institute of Neurological Disorders and Stroke, National Institutes of Health, Bethesda, MD 20892, USA; Department of Brain and Cognitive Sciences and McGovern Institute, Massachusetts Institute of Technology, Cambridge, MA 02139, USA; Laboratory on Neurobiology of Compulsive Behaviors, National Institutes of Mental Health, NIH, Bethesda, MD 20892, USA; Department of Ophthalmology and Visual Sciences, Washington University School of Medicine, Saint Louis, Missouri 63110, USA; Department of Psychological & Brain Sciences, University of California, Santa Barbara, Santa Barbara, California 93106, USA; Department of Biological Sciences, University of Maryland, College Park, Maryland 20742, USA

**Author notes:** Co-corresponding authors: Sarthak Chandra; Ila Fiete; Veronica A. Alvarez; Yi Gu. Equal contribution (Ling Liang Dong, Hoon Shin, Farid Shahid).

## Abstract

Animals construct internal “cognitive maps” of the world during navigation in spatial and non-spatial domains, with grid cells in the medial entorhinal cortex (MEC) playing a key role. This requires associating internal position estimates with external cues to reduce spatial uncertainty over time. However, how grid cell representations evolve in novel spaces to support map formation is unclear. To address this question, we longitudinally imaged calcium dynamics of grid cells over 10 days as mice learn operant tasks in novel virtual linear tracks. We observe that spatial tuning of grid cells is present immediately in novel tracks but evolves as a significant fraction of spatial fields shift backward on a run-by-run basis, within and across days. Backward shifts are more prevalent and persistent in successful learners. The fields gradually stabilize across days, anchored by landmarks, suggesting slow plasticity. The backward shifts partially reset daily, reflecting a slower consolidation timescale. While individual fields of a cell shift differentially, co-active fields of co-modular grid cells shift together, indicating their coupled dynamics on the same two-dimensional torus. Spatial learning leads to systematic changes and stabilization of their population phase trajectory, including lateral shift, rotation, and phase resets at landmarks, forming a landmark-fragmented representation for the environment. Next, we build an entorhinal-hippocampal model that provides a mechanistic explanation of the diverse phenomena - grid field shifts, increasing fidelity, and fragmentation of the spatial map - and predicts slow Hebbian plasticity in the hippocampus-to-entorhinal pathway. Supporting this, electrophysiology demonstrates that learning-performance-correlated weakening of local inhibition facilitates potentiation of indirect hippocampal inputs to superficial MEC. Together, our study provides multifaceted evidence of slow hippocampus-to-MEC plasticity, elucidating the formation of stable and fragmented cognitive maps that combine internal and cue-driven positional estimates in rich environments during learning. This mechanism may extend to broader memory processes involving this circuit.

## Introduction

The ability of animals to form an internal “cognitive map” of an environment based on cues such as landmarks and boundaries is critical for navigating complex landscapes (*1*). In mammals, this map is primarily represented by spatially modulated cells in the hippocampus and entorhinal cortex (*2*). Recent studies show that the same cells form cognitive maps of non-spatial domains (*3, 4*). Key components of the map include place cells in the hippocampus (*5*) and grid cells in the medial entorhinal cortex (MEC) (*6*), with place cells active at one or a few locations, and grid cells forming a regular tiling pattern in open arenas. In more complex environments, grid cells fragment the space by forming a number of regular local maps with discontinuities between them (*7*).

We focus on two key questions: How are maps, which involve associating internal positional estimates with external cues, formed? What is the role and dynamics of grid cells in this process? It is known that place cells (*8–10*) and grid cells (*6, 11*) quickly exhibit spatially tuned responses in novel environments, with behavioral time scale synaptic plasticity (BTSP) playing a role in rapid place field emergence (*12*). In grid cells, the rapid appearance of fields in novel environments can be attributed to their internal generation dynamics (*13–15*), which exists even during sleep (*16–18*), and by cue-driven inputs (*11*) and reward locations (*19, 20*). On a very long time-scale (weeks to month), place fields appear and disappear but their combined responses still encode spatial information about the environment (*12, 21–24*).

The timescales of hours to few days, where actual mapping occurs – the bringing of external cue-driven responses into register with internally inferred positional estimates so that they can be used to correct each other (*25–28*) (**Fig. 1a**) – is our focus here. Over this timescale in open fields, the initially less consistent and expanded grid field responses gradually refine and become more regular (*29–31*). On one-dimensional (1D) tracks traversed in one direction, place fields shift backwards with changes in field skewness and width (*24, 32–37*). In other words, the representations in both entorhinal cortex and hippocampus gradually adjust during map acquisition, but it is unclear how they might relate and what plasticity mechanisms drive this refinement.

**Figure 1:**
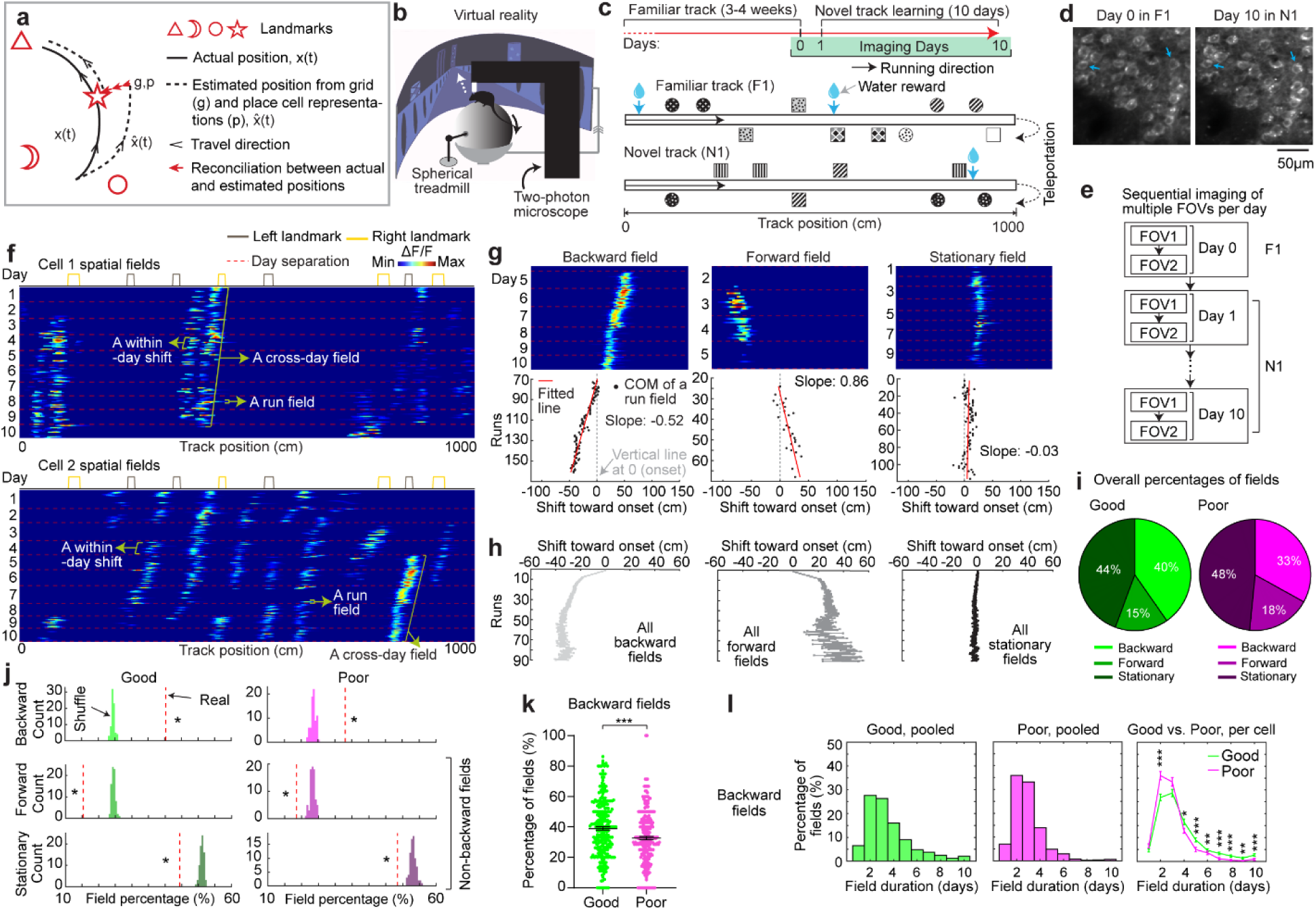
MEC grid cells display activity fields with continuous run-by-run shifts across multiple days **a.** Schematic of forming accurate internal spatial map by grid and place cells for navigation via spatial learning. During spatial learning, internal representation of external environment and estimation of movement are corrected by external landmarks. **b.** Two-photon imaging setup, adapted from Malone et al. 2024 (*38*). **c.** Experiment design. Top: experiment schedule. Mice perform 10 days novel track (N1) learning after training in a familiar track (F1). Bottom: one-dimensional virtual reality (VR) design of the familiar and novel tracks, adapted from Malone et al. 2024 (*38*). **d.** Example of the same field of view (FOV) in F1 and day 10 in N1, adapted from Malone et al. 2024 (*38*). **e.** Imaging workflow. One or multiple FOVs were imaged for one mouse per day in the same order. **f.** Example calcium dynamics of cross-day fields of two grid cells. **g.** Examples of three types of cross-day field (backward, forward, stationary). For each type, top: calcium dynamics of a fields; bottom: centers of mass (COMs) of run fields with their linear regression results. **h.** Averaged trajectories of all the cross-day fields in the three shift types. **i.** Percentage of three cross-day field types, calculated by pooling all cross-day fields of the same shifting type across grid cells and FOVs together. **j.** Comparison of the percentage of three cross-day field types with percentage distribution of run-shuffled fields. **k.** Percentage of backward fields per cell for good and poor performers. **l.** Comparison of cross-day field duration (days) between good and poor performers. Left and Middle: duration distribution of backward fields of good and poor performers. Right: Comparison of the percentage of backward fields with different durations per cell between good and poor performers. *p ≤ 0.05 or the original fraction significantly higher than shuffled fraction in **j**, **p ≤ 0.01, ***p ≤ 0.001, n.s. p > 0.05. Statistical test results are listed in Table S1. Error bars represent mean ± SEM.

Here, we seek a deeper understanding of how map learning unfolds in the brain. To this end, we conducted longitudinal two-photon calcium imaging in the MEC as mice learned novel virtual linear tracks containing local cues over a period of 10 days. The cues served as strong anchoring landmarks that led to the formation of a fragmented grid cell map. We discovered that, as seen earlier for place cells (*24, 32, 34, 37*), grid fields gradually shifted backward on a run-by-run basis with a gradual consolidation process, in which each day’s learning was only partially retained. This finding raises the possibility that grid cell shift might drive or be driven by place cell shift; however, in grid cells, the backward shift is interpretable as revealing a growing map fragmentation in which a consistent map is used within inter-cue intervals but the maps between cues are increasingly different.

We built an entorhinal-hippocampal model that captures diverse phenomena and predicts that backward shifts are shaped by temporally delayed Hebbian plasticity in a projection from cue-driven hippocampal cells to grid cells. Using *ex vivo* slice electrophysiology, we found that plasticity in an indirect synaptic pathway from the hippocampus to grid cells was correlated with spatial learning performance, consistent with the predicted temporally delayed Hebbian plasticity pathway of the model. These combined approaches provide multifaceted evidence for a Hebbian plasticity-based mechanism in synapses from the hippocampus to the MEC that gradually creates a spatially stable fragmented grid map anchored to salient environmental features. This work offers new insights into how the brain integrates internal and external information during spatial learning to construct flexible representations, a core function of spatial cognition.

## Results

### Grid fields continuously shift to earlier positions within and across days

To characterize grid cell activity over extended spatial learning, we analyzed MEC activity in GP5.3 mice (*38*) unidirectionally navigating virtual reality (VR) linear tracks for multiple runs each day, which stably expressed a fluorescent calcium indicator GCaMP6f in excitatory neurons in layer 2 of the MEC (*38–40*) (**Fig. 1b**). The mice first explored a track until familiarization (familiar track, F1), then underwent 10-day spatial learning on a novel track (N1) featuring a reward and eight new landmarks at different locations compared to F1 (**Fig. 1c**). We tracked calcium dynamics in the same group of MEC cells from F1 to day 10 of N1 (**Fig. 1d**). Each day, 1-3 fields of view (FOVs) in each mouse were imaged in the same order (**Fig. 1e**). Of the 15 participating mice, 11 were classified as good performers and 4 as poor performers based on their reward-predictive behaviors on N1, indicating successful and unsuccessful spatial learning, respectively (*38*). Neurons were classified as grid cells based on reliable spatial field patterns (a field is defined by elevated run-averaged calcium responses above chance in adjacent spatial bins (*38*)) on at least two days without conjunctive cue activity, defined as having responses locked to multiple landmarks (*41, 42*) (**Extended Data Figure 1a**). We focused on grid cells active across the 10 days of N1 learning.

We observed that most grid cells exhibited spatial activity in the first run in N1 (**Extended Data Figure 2a**), and the number of significant spatial fields detected on a run-by-run basis (“run fields”) remained largely stable across runs (**Extended Data Figure 2b**). These results are consistent with previous observations that grid fields appear immediately upon environmental change (*6, 11*), supporting the concept of internally generated grid cell dynamics as predicted by attractor models and confirmed by experimental data (*13, 16–18*). Remarkably, grid fields appeared to consistently shift backward to earlier track locations on a run-by-run basis within and across days (**Fig. 1f and Extended Data Figure 1b**), suggesting a gradual adjustment in spatial representation. To characterize this shift, we connected adjacent run fields within and across days (“cross-day fields”) (**Extended Data Figure 3a**). We determined the shift directions of cross-day fields based on the slope of the best-fit line through their centers of mass (COMs). Backward or forward shifting cross-day fields had negative or positive slopes, respectively (**Fig. 1g, h**). The significance of the shift was assessed by comparing the slope with bootstrap slopes of circularly shifted run fields (**Extended Data Figure 3b-d**). This approach revealed cross-day fields that significantly shifted backward (backward fields), forward (forward fields), and those without significant shifts (stationary fields, **Fig.1g, h**). All three shift types were observed in good and poor performers (**Fig. 1i**) and the same cells could display fields in different shift types (**Extended Data Figure 2c**). We then compared the abundance of these field types to random chance by shuffling cross-day fields 100 times within runs and reidentifying their shift features (**Extended Data Figure 3e**). The percentage of backward fields exceeded random chance, while stationary and forward field occurrences were below random (**Fig. 1j**). Therefore, we focused on backward fields and combined forward and stationary fields as non-backward fields.

We examined the characteristics of backward fields in good and poor performers. Good performers had a higher percentage of backward fields per cell than poor performers (**Fig. 1k**). Although backward fields persisted for various durations (the number of days a field was observed) in both groups (**Fig. 1l, left and middle**), good performers had relatively fewer short-duration fields (≤ 3 days) and more long-duration fields (> 3 days) on a per-cell basis (**Fig. 1l, right**). These trends were also seen in non-backward fields but were less pronounced (**Extended Data Figure 4a and b**).

Overall, the significant presence of backward fields reflects dynamic changes in grid cell spatial representation during novel environment learning. The higher abundance and persistence of these fields in good performers suggest that they are associated with successful spatial learning.

### Backward field shifts gradually stabilize during learning

If backward fields contribute to learning, their spatial activity change should saturate over successful learning to produce a stable spatial representation. We tested this hypothesis by examining several features of the backward fields.

We first examined the prevalence of backward fields across learning. In good performers, the number of backward fields per day (these include existing cross-day fields and new fields appeared that day and shifted backward) (Fig. 2a) remained stable during learning (Fig. 2b). The number of new backward fields decreased over days (Fig. 2c), compensated by an increasing fraction of existing backward fields (Fig. 2d). The existing backward fields also increased their fraction among all day fields (backward and non-backward fields) (Fig. 2e and f). In contrast, poor performers had lower fractions of backward fields and no learning-dependent increase (Fig. 2e and f). These results indicate that backward fields exhibit enhanced prevalence and persistence specifically during successful learning. Non-backward fields also improved their persistence, but their prevalence was lower in good performers (**Extended Data** Figure 4c-f). Thus, the high prevalence of persistent backward fields, but not other field types, is associated with successful learning.

**Figure 2:**
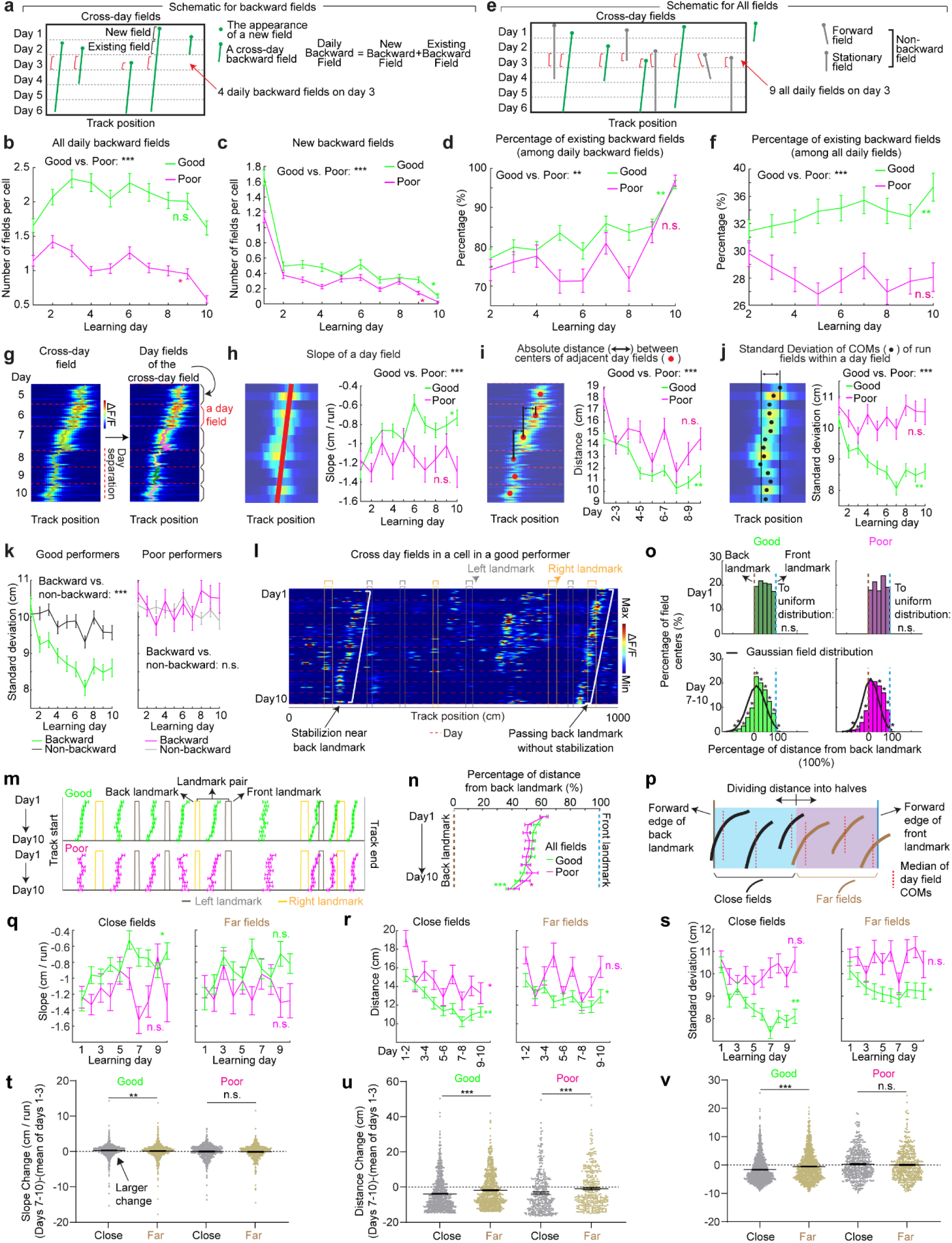
Backward grid fields stabilize and are shaped by landmarks across learning **a.** Schematic for daily, new, and existing backward fields. **b.** Numbers of daily backward fields per cell on each learning day. **c.** Numbers of new backward fields per cell on each learning day. **d.** Percentage of existing backward fields among daily backward fields per cell on each learning day. **e.** Schematic of all day fields, including backward and non-backward fields. **f.** Percentage of existing backward fields among all day fields per cell across learning days. **g.** Schematic of separating cross-day backward fields into day fields. **h.** Slopes of day fields across learning days. **i.** Distance between centers of day fields on adjacent days. **j.** COM standard deviation of run fields within day fields. **k.** Same to **j** but for backward versus non-backward fields in good (left) and poor (right) performers. **l.** A cell containing two backward fields with and without stabilization near landmarks. **m.** Averaged trajectories of backward fields initiated between individual landmark pairs, for good and poor performers. **n.** Averaged trajectories of backward fields initiated between all adjacent landmark pairs for good and poor performers. The distance between each landmark pair was normalized to 100%. **o.** Top: distribution of cross-day fields with their day 1 fields between landmark pairs. Bottom: distribution of day 7-10 fields. The percentages of fields that do or do not cross back landmarks were compared with 20000 simulated percentage values (black), each of which was constructed by the same number of fields that formed a normal distribution around current peak location and covered the same range of track position. **p.** Schematic of defining close and far fields relative to the back landmark **q-s.** Slopes (**q**) and day field center distances (**r**), and COM standard deviation of run fields within days during learning (**s**), separating close (left) and far (right) fields **t-v.** Change in **q-s** between days 1-3 and 7-10 for close and far fields. Change is calculated by subtracting the averaged value of day 1-3 from individual values on days 7-10. Each dot is one day field in day 7-10. *p ≤ 0.05 or distance bins significantly different from simulated gaussian distribution (**o** bottom), **p ≤ 0.01, ***p ≤ 0.001, n.s. p > 0.05. Statistical test results are listed in Table S1. Error bars represent mean ± SEM.

Next, we investigated whether the backward field shift pattern stabilized over learning by dividing each cross-day field by day (“day fields”) (Fig. 2g). In good performers, the per-day negative slope of backward fields gradually approached zero (Fig. 2h). The distance between the COMs of day fields, averaged across run fields within each day, decreased over days (Fig. 2i), and the standard deviation (STD) of run field COMs per day also decreased (Fig. 2j), indicating that backward fields shifted less and reached more consistent locations as learning progressed. In contrast, backward fields in poor performers exhibited persistently negative slopes (Fig. 2h), larger COM distances between days (Fig. 2i) and greater STD of run field COMs per day, without changes over days (Fig. 2j). The stabilization was also observed in the non-backward fields of good but not poor performers (**Extended Data** Figure 4 g-i). Specifically, the STDs of run field COMs for backward and non-backward fields started at similar levels on day 1, however, the STDs of backward fields decreased more robustly across days, indicating more reliable spatial representation by backward fields post-learning, despite their gradual shifts (Fig. 2k**, left**). This trend was absent in poor performers (Fig. 2k**, right**). Together, these findings underscore the significant role of backward field stabilization in spatial learning.

### Landmarks shape backward field shifts during learning

Stable mapping requires that internally derived positional estimates and estimates driven by spatial features reach an equilibrium. Landmarks, which represent key environmental features, have been shown to immediately reshape grid cell activity (*11*). We observed that across multiple learning days, as anticipated by backward shifts, backward fields gradually approached the landmarks that came before them (“back landmarks”) and moved away from those ahead (“front landmarks”). Notably, some backward fields near their back landmarks appeared to stabilize near them, while some passed the landmarks without stabilizing (Fig. 2l). To further explore this phenomenon, we investigated how landmarks influenced backward field shift during learning.

We first analyzed field trajectories in relation to adjacent landmarks (Fig. 2m). Backward fields were grouped between each adjacent pair of landmarks, with landmark locations defined by their front edges. The start and end of the track were also considered landmarks due to their similar visual impact. For both good and poor performers, the average trajectories of most field groups gradually shifted toward their back landmarks without crossing them (Fig. 2m). This pattern persisted when all groups were combined (Fig. 2n). On day 1, day fields were evenly distributed between adjacent landmarks. By days 7 to 10, a significant number of fields accumulated near their back landmarks, and fewer fields crossed the landmarks compared to a Gaussian distribution with the same peak location and track coverage (Fig. 2o, bottom, black curve) representing a symmetrical distribution scenario. These results suggest that back landmarks attract backward fields.

To further determine if landmarks arrest the backward shift of nearby fields, we compared stabilization levels of backward fields at varying distances from back landmarks. We divided the distance between each adjacent landmark pairs into halves, categorizing the fields as either “close” or “far” from the back landmark based on the distance from the field location (median of day field COMs) and the back landmark (Fig. 2p), We then combined all close or far fields across adjacent landmark pairs to analyze their stabilization during learning. In good performers, close and far fields exhibited significant differences, with only close fields showing reliable stabilization. Close field slopes approached zero (Fig. 2q), distances between adjacent day field COMs decreased (Fig. 2r), as did the STDs of run field COMs within days (Fig. 2s). Learning-dependent changes in all these parameters, calculated as parameter values on days 7-10 minus the averaged value on days 1-3, were greater for close fields, indicating greater stabilization (Fig. 2t**-v**). These trends were also observed in poor performers (Fig. 2q**-v**) and in non-backward fields (**Extended Data** Figure 5) but were less consistent. These results indicate that during successful learning, back landmarks preferentially stabilize nearby backward fields, whereas fields proximal to front landmarks are less stabilized.

Overall, during learning, back landmarks attract and stabilize nearby backward fields, leading to an over-representation of landmark information.

### Grid field locations partially reset between adjacent learning days

On each learning day, the mice experienced the novel track only during their relatively brief VR sessions. We sought to examine the offline dynamics of their grid fields. A fast offline memory consolidation mechanism (*43*), or offline predictive plasticity mechanism (*43, 44*), could manifest as a further backward field shift on the first run of a day relative to the last run on the previous day (“progression”) (Fig. 3a**, left**). Alternatively, if stable memory formation involves a slower consolidation process across many days, the latest learning might not fully persist and would be visible as a forward shift (“regression”) of fields relative to their positions on the previous day (Fig. 3a**, middle**). For offline changes to be statistically significant, their magnitudes should exceed typical adjacent-run field shifts within days (Fig. 3a**, right**).

**Figure 3.**
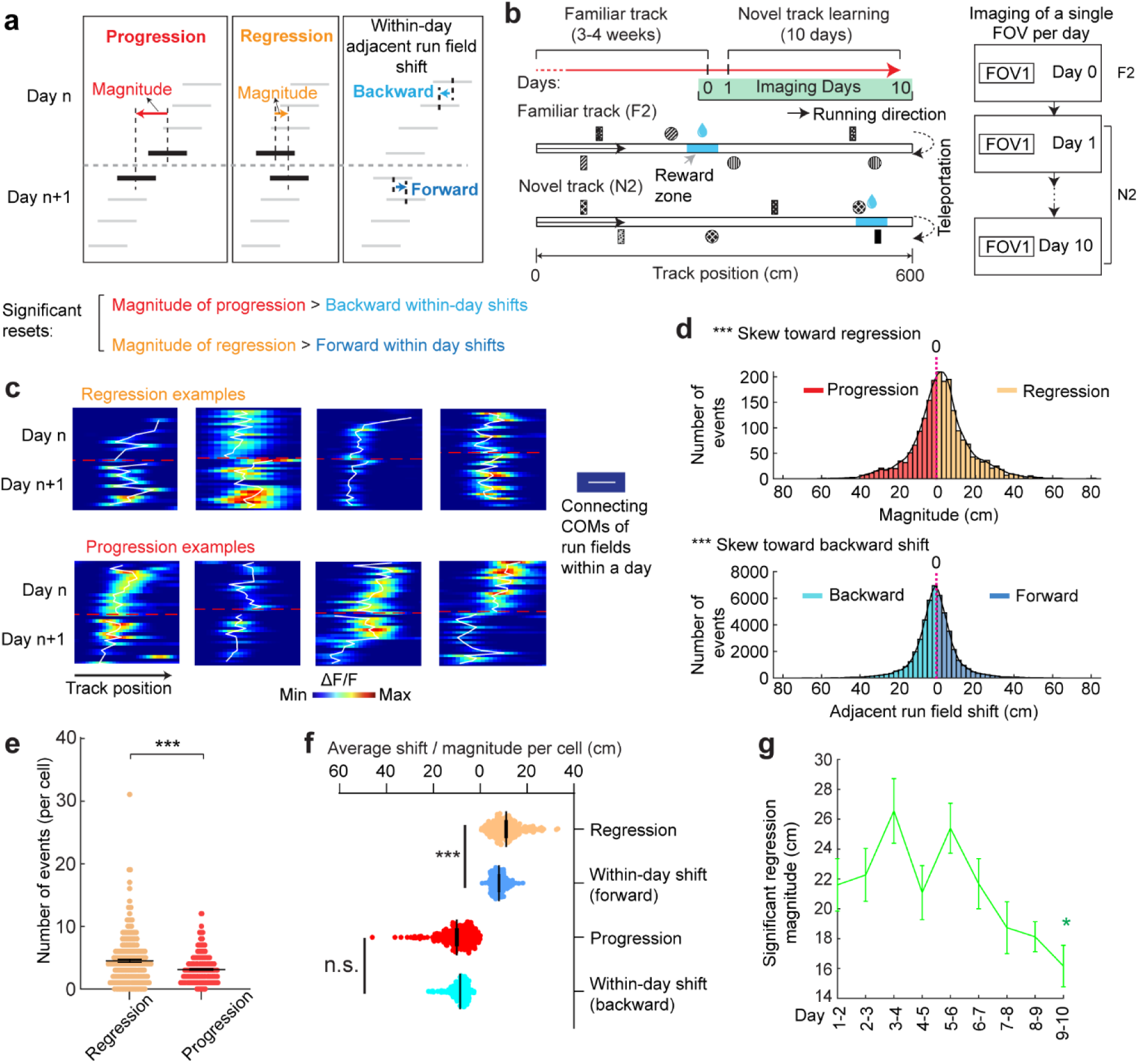
Backward fields partially reset between adjacent days **a.** Schematic of progression and regression events, and the within-day adjacent run field shifts in the same direction (backward and forward, respectively). The magnitude of a progression or regression event is represented as the shift distance between the COMs of the first run field on day n+1 and the last run field on day n. **b.** Schematic of the experiment design for dataset 2. Left: experiment timeline. After 3-4 weeks of training in a familiar track (middle, F2), mice underwent a 10-day novel track learning (bottom, N2). The mice had to correctly stop within a hidden reward zone (blue) to trigger water reward delivery. Right: imaging schedule. Only one FOV was imaged per day in this experiment, thus no additional online learning was missed in between the two imaging sessions. **c.** Example regression and progression events. **d.** Magnitude distribution of regression and progression events, and within-day adjacent run field shifts in forward and backward directions, combining all backward fields. **e.** Numbers of regression and progression events per cell. **f.** Comparing the distances of regression and progression events to the within-day adjacent run field shift in the corresponding direction. **g.** Magnitudes of significant regression events (above 85^th^ percentile of adjacent run field shifts) across days. *p ≤ 0.05, **p ≤ 0.01, ***p ≤ 0.001, n.s. p > 0.05. Statistical test results are listed in Table S1. Error bars represent mean ± SEM.

Testing offline progression versus regression effects and thus drawing conclusions about the existence of consolidative dynamics requires that we compare field locations just before the offline interval with those right after the interval. However, in dataset 1, multiple FOVs were imaged daily, meaning that animals continued to perform online learning on the same day while imaging other cells (Fig. 1e). This protocol could induce apparent but false progression effects because of continued backward shifting of online not offline dynamics (**Extended Data** Figure 6a). Nevertheless, backward fields of good performers showed non-negligible regression magnitudes, suggesting the existence of true regression (**Extended Data** Figure 6b). In contrast, poor performers exhibited no significant regression or progression, consistent with having little online learning to be unlearned during the offline period.

We therefore conducted another experiment by continuously imaging calcium dynamics in the same layer 2 MEC neurons over their entire online experience as mice learned a novel track (N2) over 10 days (“dataset 2”, Fig. 3b right). Mice had to stop in a fixed reward zone to receive a water reward (Fig. 3b). This cohort included eight mice, all evaluated as good performers based on the criteria of dataset 1 (**Extended Data** Figure 7a and b). Similar to dataset 1, we identified grid cells active across the 10 days (**Extended Data** Figure 7c), with a comparable percentage of backward fields to those in good performers from dataset 1 (**Extended Data** Figure 7d). The backward fields, especially those close to back landmarks, exhibited obvious gradual stabilization during learning (**Extended Data** Figure 7e-i). Thus, the grid cell activity in dataset 2 reproduced our observations from the good performers in dataset 1.

Backward fields in dataset 2 exhibited both progression and regression (Fig. 3c), but the magnitude distribution skewed significantly toward regression (Fig. 3d). Regression events per cell outnumbered progression events (Fig. 3e), and regression but not progression magnitudes were significantly larger than the shifts between adjacent run fields in the same direction (Fig. 3f), indicating significant regression, but not progression. As expected, regression magnitudes in dataset 2 were larger than those in dataset 1, while their adjacent run field shift magnitudes within days were comparable (**Extended Data** Figure 6c). Across 10 learning days, the magnitudes of significant regression in dataset 2 (exceeding the 85^th^ percentile of adjacent run field shifts) decreased (Fig. 3g), a similar trend observed in dataset 1 (**Extended Data** Figure 6d), suggesting a gradual memory consolidation that corresponds to less offline memory loss.

The findings of significant daily regression but not progression suggest that learning-induced plasticity during online periods partially degrades during offline periods. This finding indicates the existence of a separate memory consolidation process, which requires online experience and involves a longer time scale than plasticity induction (*45–47*).

### Grid cells exhibit invariant low-dimensional population dynamics and growing map fragmentation over learning

Next, we extended our analysis from the dynamics of individual backward grid fields to the global population dynamics of whole grid modules. We asked whether module activity remains low-dimensional during learning of a new environment when grid fields shift, interact with landmarks, and settle. Though we expect that grid cells maintain a low-dimensional dynamics based on continuous attractor network models (*13–15, 48*) and demonstrations under many conditions (*11, 16–18, 38, 49, 50*), it is unclear if they do so during the formation of the spatial map where we have observed that the fields of even a single neuron shift differentially relative to each other **(Extended Data** Figure 1b **and Extended Data** Figure 2c).

In good performers from dataset 1, we identified “co-modular” grid cells (cells with similar spacings between nearest fields) (*51*) (**Extended Data** Figure 8), and examined whether these cells exhibited preserved attractor dynamics across learning (*52*), so that population states remained localized on the same 2D toroidal manifold. We found that the temporal activity correlations between simultaneously imaged co-modular grid cell pairs (“pairwise correlation”) were highly preserved across runs (significantly higher than shuffles, Fig. 4a), reflecting stable cell-cell activity relationships over learning (*38*).

**Figure 4.**
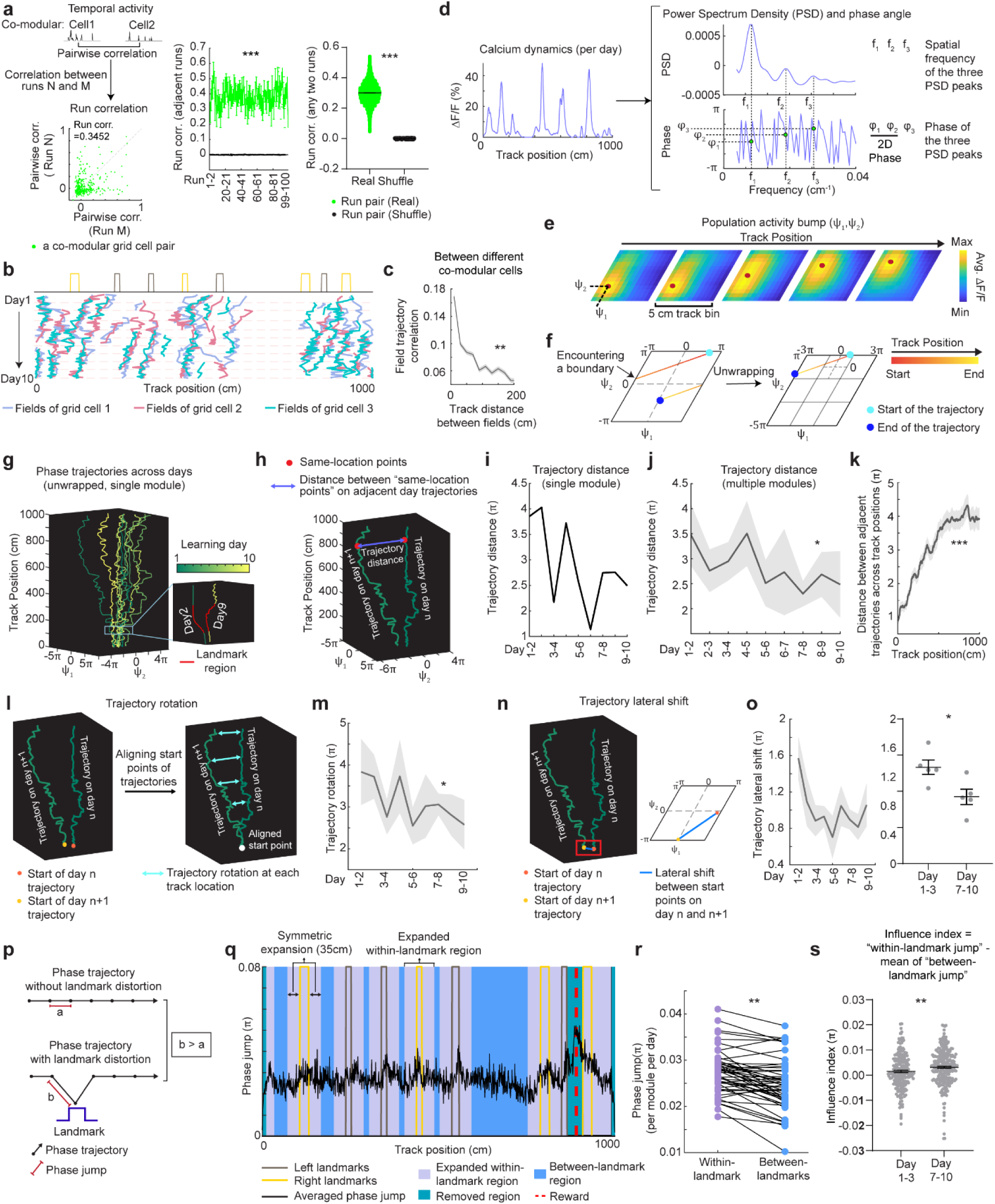
Population dynamics of grid cells over learning remain low-dimensional with collective phase shifts that gradually stabilize **a.** Temporal activity relationships of co-modular grid cells are maintained across runs. Left: the consistency of the relationship was quantified as “run correlation”, which is the correlation between the pairwise correlations of temporal activities of the same co-modular grid cell pairs on different runs (N and M here). Middle: run correlation between adjacent runs for all modules, compared to shuffles. Right: run correlation between all run pairs for all modules, compared to shuffles. **b.** Field trajectories of three co-modular cells, showing similar shift for fields that are spatially adjacent but belong to different cells. **c.** Correlation between cross-day fields from different co-modular grid cells decrease with increased field distance. Correlation is calculated between the field segments of two cross-day fields within the same runs. Correlations were pooled within non-overlapped 20-cm track bins. **d.** Schematic of determining the phase coordinates of grid cells. Left: calcium dynamics of a grid cell, averaged across runs within a day. Right: Power spectrum density (PSD, top) and the corresponding phase (bottom) were calculated from the cell’s calcium dynamics on the left. **e.** Combining phase distribution and calcium dynamics of each cell at individual track locations, an activity bump was revealed and translated within the rhombus. Ψ_1_and Ψ_2_represent bump coordinates within the phase rhombus. **f.** Unwrapping to solve the issue when the bump trajectory encounters boundaries of the rhombus. Left: an example trajectory encountering boundaries of the rhombus. Right: unwrapped trajectory. **g.** Phase trajectories of a group of co-modular grid cells on individual learning days. Right bottom: zoomed in region around the first landmark of the trajectories on two days. **h.** Schematic of distance calculation between two phase trajectories on adjacent days (trajectory distance). The distance was calculated between each pair of points at the same track location on the two trajectories (“same-location points”). **i.** Trajectory distance for the example in **g**. **j.** Trajectory distance between adjacent days, across all modules. **k.** Trajectory distances of same-location points on adjacent day trajectories from the beginning to the end of the track, combining all modules. **l.** Schematic of trajectory rotation estimation based on trajectory distance between adjacent days. The starts of the two trajectories were aligned to the same point to eliminate the lateral shift. **m.** Averaged trajectory rotation across modules between every adjacent learning day. **n.** Schematic of trajectory lateral shift estimation based on the distance between the starting points of adjacent day trajectories. **o.** Trajectory lateral shift across learning. Left: Averaged trajectory lateral shifts across modules between adjacent learning days. Right: Comparing trajectory shifts on days 1-2, 2-3 versus on days 7-8, 8-9, and 9-10. Shifts on day 1-2, 2-3 were averaged within each co-modular cell group, as well as shifts on day 7-8,8-9,9-10. Each dot is a co-modular cell group. **p.** Schematic of phase jumps with and without landmark distortion. **q.** Phase jumps along the track. The jumps were averaged across all grid modules and all 10 days. Phase jumps within the first and last 10 cm of the track, as well as those around the reward (863-909 cm), were removed from analysis. **r.** Comparison of phase jumps between within- and between-landmark regions. Each line represents the jumps of one grid module on one day. **s.** Landmark distortion, quantified by “influence index” in early (days 1-3) and late (days 7-10) learning. Each dot is one run on day 1-3 or day 7-10. *p ≤ 0.05, **p ≤ 0.01, ***p ≤ 0.001, n.s. p > 0.05. Statistical test results are listed in Table S1. Error bars represent mean ± SEM.

Thus, we have two seemingly opposing findings – that the fabric of cell-cell relationships is invariant even though firing fields shift differentially relative to each other in single cells, indicating large spatial tuning distortions. These can only be reconciled if the differential field shifts in a cell are shared population-wide across co-modular cells: that is, the entire population state shifts coherently across cells, but differentially relative to the underling physical location of the animal at different locations. Accordingly, we examined the shifting of spatially overlapping fields from different co-modular cells, finding that they shifted in tandem across cells (Fig. 4b). Consistent with this, spatially nearby fields from different co-modular cells (based on median of run field COMs) exhibited high trajectory correlations, while those further apart showed much lower correlations (Fig. 4c). Thus, well-separated fields of even a single cell could shift differentially relative to external space, the co-active fields of different cells shift together, indicating preserved population coherence and low dimensionality coexisting with collective state changes relative to space that correspond to field shifts.

Finally, we considered the mapping from intrinsic low-dimensional population states to the animal’s spatial position, how this mapping changes over learning, and how these changes relate to observed grid field shifts. Given the evidence of preserved low-dimensional population dynamics, we estimated a population phase from simultaneously imaged co-modular grid cells in good performers using the Yoon-Lewallen technique, which is based on Fourier transforms of the 1D spatial responses of individual cells (*53*). Specifically, Fourier transform produced Power Spectrum Density (PSD) of each cell with three largest peaks in spatial frequencies f1, f2, and f3 (Fig. 4d, top right). f1, f2, and f3 determine the angle of an 1D slice through the underlying two-dimensional (2D) grid that generates the 1D spatial firing fields. For spatial responses of simultaneously imaged co-modular grid cells that are generated from a common underlying 2D triangular grid, f1, f2, and f3 should be the same for the cells, and the sum of f1 and f2 should equal to f3. The 2D phase (φ_1_, φ_2_) of a cell in a phase rhombus was extracted from the Fourier transformation at the locations of f1 and f2 (Fig. 4d, bottom right). Next, we combined the phases of individual co-modular cells with their evolving activity along the track to obtain a moving population activity bump that traced out a population phase trajectory over the track (represented by Ψ_1_, Ψ_2_) (Fig. 4e). To more easily visualize the trajectory with periodic boundary conditions within the phase rhombus (Fig. 4f, left), we unwrapped the population phases across multiple rhombuses, obtaining a largely unidirectional population phase evolution (Fig. 4f, right).

For precise population dynamics estimation, the above analyses were applied to five simultaneously imaged co-modular cell groups with more than 10 cells, whose sum of averaged f1 and f2 across days was near the averaged f3 (difference < 8% of averaged f3) (**Extended Data** Figure 9). We observed that their population phase trajectory changed over learning (Fig. 4g), with the changes decreasing over time. This was quantified via a trajectory distance between adjacent days, defined as the average of the set of point-wise Euclidean distances between the points on the two trajectories at the same track position (Fig. 4h**-j**). Overall, the phase trajectory changed through rotation above the starting point, reflected by the gradually increasing trajectory distances for the beginning toward the end of the track (Fig. 4k). The rotation decreased over learning (Fig. 4l**-m**). The small lateral shifts in the starting phase also decreased over learning, with larger shifts across days 1-3 than days 7-10 (Fig. 4n**-o**). The rotations and lateral shifts of the population phase trajectory as animals run on the same track over learning potentially cause the observed gradual spatial shifts of the grid fields.

Next, we examined how landmarks influence the phase trajectory and shape map fragmentation. If landmarks strongly modify the positional estimate and drive map fragmentation, the population phase should exhibit larger changes (phase jumps) near landmarks than far from them (Fig. 4p). Indeed, phase jumps were bigger around landmarks (Fig. 4q) and near the track end (which contains two landmarks as well as a water reward, known to attract grid fields(*19, 20*)). To isolate influence of landmarks and remove reward and teleportation effects, we excluded regions near the reward and near the start and end, finding that within-landmark jumps were still larger than between-landmark jumps (Fig. 4r). We quantified the evolution of landmark influence over learning by defining an “influence index”: the average within-landmark jump magnitude minus the between-landmark jump per day. Landmark influence on late days (days 7-10) was higher than on early days (days 1-3) (Fig. 4s), suggesting a growing map fragmentation.

These results show that over learning, the internal population state traces a trajectory on an invariant 2D torus. This trajectory gradually rotates and shifts, and it also exhibits discontinuities or phase jumps at landmarks. The rotations and shifts decrease over learning, indicating a stabilization of the map, while the discontinuities at the landmarks are greater on later days than earlier days, meaning that learning resulted in a more fragmented map.

### A mechanistic model of map learning explains diverse phenomena and predicts delayed hippocampal-to-entorhinal plasticity

To model the mechanisms underlying the above grid cell dynamics, we turned to a neural circuit model of the entorhinal-hippocampal network and considered the role of experience-dependent plasticity in gradually modifying the internal and cue-driven dynamics of the grid cell response. Our model builds on the theoretical Vector-HaSH framework(*54*), extending the model by generalizing it to continuous space and modeling slow plasticity in the synapses from the hippocampus to grid cells.

Our model consists of three subcircuits (Fig. 5a): (a) multiple grid cell modules, each modeled as a continuous attractor integrator network (*13*), (b) sensory entorhinal cells, whose states are representations of the observed landmarks, and (c) hippocampal cells, which receive inputs from sensory entorhinal cells and grid cells to provide a combined estimate of the animals’ position in the environment. The circuit integrates two types of information about animal position: Grid cells compute position from noisy self-movement velocity estimates, and landmarks send positional information to sensory entorhinal cells (which is represented in the form of high-dimensional feature vectors). Hippocampal cells receive the sensory cell inputs and drive grid cells to influence their population firing phases; then grid cells update their states by integrating their incoming velocity inputs. We assume that the sensory-hippocampus and hippocampus-grid cell synapses undergo slow Hebbian-like synaptic plasticity, and that hippocampal activity is slightly delayed before it influences plasticity in the hippocampus-grid synapses.

**Figure 5.**
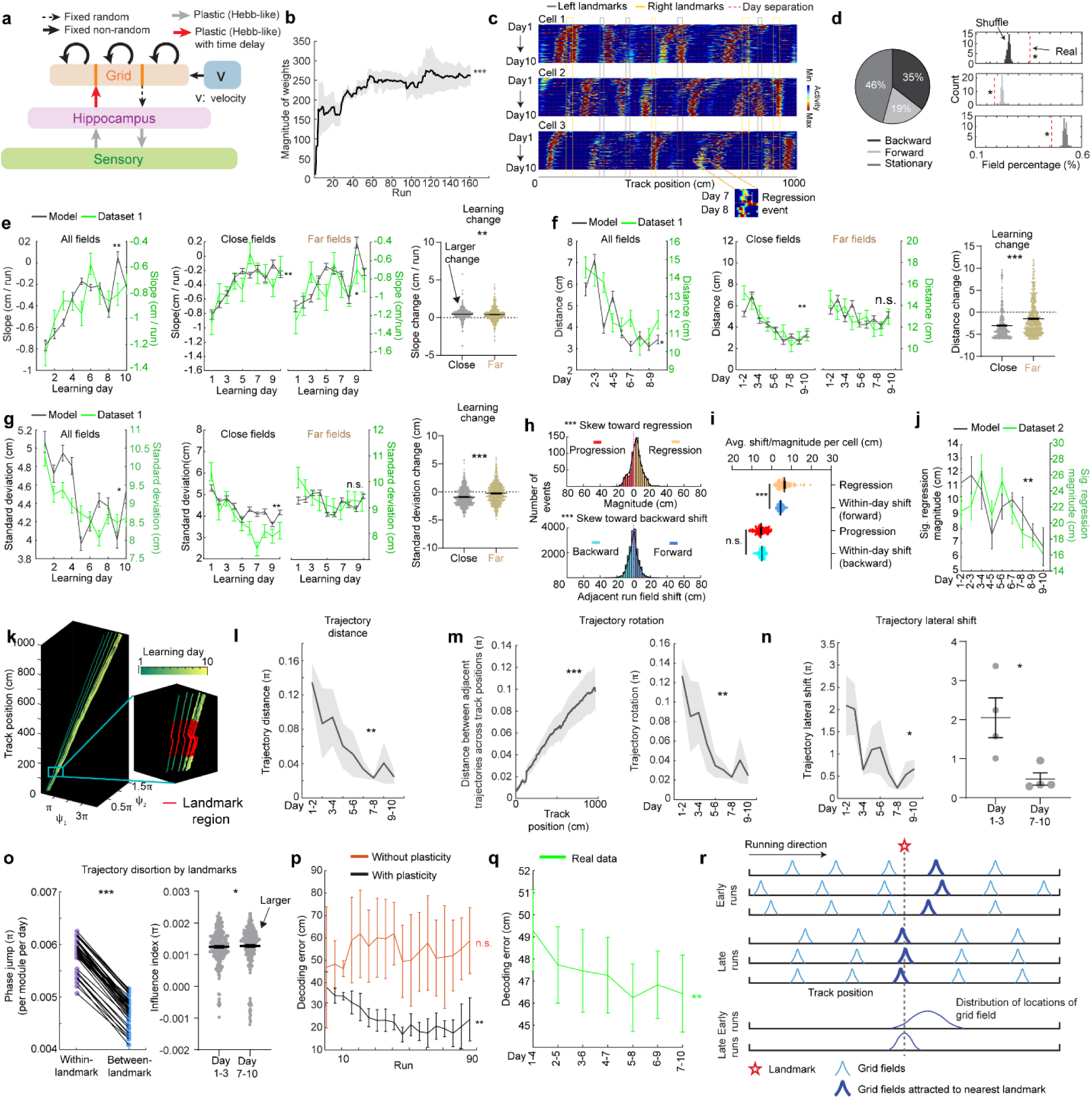
Delayed Hebbian plasticity from hippocampus to grid cells in Vector-HaSH reproduces changes and increased spatial knowledge over map learning **a.** Vector-HaSH model (*54*) with the addition of Hebbian plasticity from hippocampus to grid cells. **b.** Magnitude of the weights of the hippocampus-grid synapses across runs. **c.** Cross-day activity patterns of three simulated grid cells, showing shift fields. A field regression event is shown for cell 3 between days 7 and 8. **d.** Fractions of cross-day fields in three shift types. Left: original fraction of detected cross-day fields. Right: comparison of the percentage of three cross-day field types toward the percentage distribution of shuffled fields. **e.** Slope of day fields during learning: for all fields (left), close versus far fields (middle), and learning change from days 1-3 to days 7-10 (right). The calculations were identical to those for real data (Fig. 2h, 2q, and **2t**). The curves for real (green) and simulated data (black) are plotted together to demonstrate the consistency, similar in the following plots up to panel **j**. **f.** Similar to **e** but for distances of adjacent daily field centers. **g.** Similar to **e** but for COM standard deviation of run fields. **h.** Distribution of regression and progression events, and within-day adjacent run field shifts in the corresponding directions, across all simulated backward fields. **i.** Comparing the magnitude for regression and progression events toward the within-day adjacent run field shift of the same direction **j** Magnitude of significant regression events (>70^th^ percentile of run field shifts) across learning. The curves for real (green) and simulated (black) data are plotted together to demonstrate the consistency. **k.** Population activity trajectories of the simulated grid cells across 10 days. Right bottom zoom: all trajectories within the area of a landmark. **l.** Trajectory distance between trajectories in adjacent days in **k**. **m.** Left: trajectory rotations suggested by gradually increasing distances from the beginning to the end of the trajectories across track locations. Right: averaged trajectory rotation across modules between adjacent learning days. **n.** Left: averaged lateral shift between start points of trajectories in adjacent days. Right: comparison between the shifts on day 1-3 and day 7-10 of the learning. Each dot represents one simulation condition. **o.** Left: comparison of phase jumps between within- and between-landmark regions. Right: influence index in early (days 1-3) and late learning (days 7-10). Each dot represents one run in day 7-10. **p.** Location decoding error of simulated grid cells from the model with and without plasticity at sensory-to-hippocampus and hippocampus-to-grid synapses across runs. **q.** Location decoding error of real grid cells from dataset 1 across days. **r.** Schematic of noisy grid fields gradually stabilize at landmarks across spatial learning under Hebbian-plasticity based mechanism. *p ≤ 0.05 or the original fraction significantly higher than shuffled fraction in **d** right, **p ≤ 0.01, ***p ≤ 0.001, n.s. p > 0.05. Statistical test results are listed in Table S1. Error bars represent mean ± SEM.

We simulated this model on a 1000-cm linear track with eight landmarks, to match experimental conditions in dataset 1. Right or left landmarks activated different subsets of sensory cells (e.g., cue cells in the MEC (*42*)), and each landmark on a given side induced a different (random and fixed) pattern of sensory activity. The start and end track boundaries were modeled as strong inputs driving both left and right sensory subsets. We used the recorded trajectories of a mouse from dataset 1 which ran 161 runs over 10 days to compute velocity inputs, with the addition of noise.

This model reproduced diverse experimental observations. The hippocampus-to-grid synapses weights gradually increased across runs (Fig. 5b) and drove field shifts over days (Fig. 5c). We observed stationary fields, backward and forward fields (Fig. 5d), with a strong bias towards backward shifts (as in experiments, the percentage of backward fields exceeded random chance, while stationary and forward field occurrences were below random, Fig. 5d right). The dominance of backward shifts can be understood in the model: in-between landmarks, circuit dynamics are entirely determined by grid and hippocampal interactions. Plasticity in the return grid-to-hippocampal weights associates the hippocampal state with a slight time delay to the grid state, while the hippocampal state is determined by a fixed random projection from grid cells. Effectively, this means that the weights learn to map the current grid state to a slightly delayed hippocampal state, which is in turn determined by the grid state. Thus, the weights associate the current grid state to an earlier one, leading to the backward shift in grid fields. In turn, this mechanism would also predict that hippocampal fields should shift backwards, observed previously (*24, 32, 34, 37*). Consistent with this, the slope of the backward shift was controlled by the temporal plasticity delay duration (Fig. 5a).

When landmarks are present, they provide another source of input to the hippocampus (*42, 55–59*). Backward field shifts stabilized over learning as in good performer mice (Fig. 5e**-g**, left). Stabilization was stronger for fields closer to back landmarks (Fig. 5e**-g**, middle and right), matching the experimental data. The mechanism of stabilization in the model is a gradual equilibration of the push-pull between delayed plasticity-induced backward field movement and field placement by forward path integration-based phase advance.

Furthermore, adding a slower consolidation timescale to the plasticity process in the same pathway reproduced the partial resetting of learning between days shown in Fig. 3. We assumed that weights continuously update, but if plasticity-inducing experience is interrupted for an extended period, such as the end of a navigation session, they decay to values one consolidation timescale before the interruption. As a result, at the start of each navigation session (“day”), the backward shifting fields exhibited partial position regression (Fig. 5h**-j**), with decreasing amounts of regression (because there is decreasing drift) over learning.

The population states of grid cells in the model stay on a 2D torus (by construction, as grid modules were low-dimensional continuous attractor networks), so we directly analyzed the evolution of their population phase trajectories over learning from multiple simulation runs with different initial conditions. As seen in the mice, the population trajectory exhibits overall shifts (Fig. 5k and l), rotations about the starting point (Fig. 5m), and lateral shifts of the starting points (Fig. 5n), and these dynamics gradually stabilized over learning. Phase trajectories exhibited within-landmark jumps because of the sensory-driven hippocampal inputs (Fig. 5o **left**), and the learned trajectories were increasingly fragmented, with greater landmark influence (larger within-landmark jumps) later in learning (Fig. 5o **right**). This increasing fragmentation can be interpreted as a consequence of the backward drift of grid fields, in conjunction with the pinning of grid fields due to the influence of local landmarks. The shift in grid fields in the between-landmark region corresponds to a shift in the trajectories in phase space. However, since the grid fields are stabilized at back landmarks (Fig. 2q-v), the region of phase trajectory corresponding to landmarks does not shift in phase space as much. Thus, the backward shift in grid fields leads to an increasing fragmentation of the phase trajectory due to differential shift in the between-landmark region versus at the landmarks (quantified as the influence index, Fig. 4s and Fig. 5o **right**).

Finally, with the help of the model, we connected synaptic plasticity-driven field dynamics with spatial learning and function: We examined spatial information by quantifying the accuracy of position decoding from simulated grid cells across learning in our model, finding that the decoding error significantly decreases across learning (Fig. 5p). The decoding error remained higher in an alternate model with no plasticity in the sensory-to-hippocampus and hippocampus-to-grid synapses, and consequently with stationary grid fields (**Extended Data** Figure 10a and b). When we tested grid cells in mice, we found higher-fidelity spatial representations on later days (i.e., lower decoding error, Fig. 5q). Together with the model, this finding suggests that slow synaptic plasticity in hippocampal-to-grid synapses may be responsible for this increase in spatial fidelity.

Though plasticity in the hippocampus-to-grid cell pathway increases spatial fidelity, the delayed aspect of plasticity (required for backward-shifting fields) is not necessary for accuracy: implementing the plasticity without a delay in the hippocampal activity resulted in improved decoding without backward field shifts (**Extended Data** Figure 10a and b). Backward shifts are, however, responsible for increasing map fragmentation over learning due to the shifts of grid fields away from landmarks and relative stabilization of grid fields at landmarks.

In sum, our model recapitulated many features of experimental data and predicts that Hebbian plasticity from sensory to hippocampal cells and delayed Hebbian plasticity-based from the hippocampus to grid cells is responsible for map learning (Fig. 5r).

### Synaptic change in the indirect hippocampal output to the MEC is associated with spatial learning

We next test our model’s prediction about the plasticity in the hippocampus-to-MEC pathway for spatial learning. We hypothesize that larger hippocampus-grid weights during learning (Fig. 5b) correspond to synaptic potentiation in the superficial layers of the MEC and should be greater in good performers than in poor performers. We also expect non-saturation of such potentiation in one environment, for good performers to continue learning in new environments.

We performed *ex vivo* slice electrophysiology to examine synaptic plasticity in the hippocampal projections to the MEC in mice that had learned virtual environments. Mice were trained to navigate VR tracks for two weeks, then learned the same N1 as in dataset 1 for another two weeks (Fig. 6a). Based on reward-predictive behaviors during the last six sessions, mice were classified as good or poor performers using criteria from dataset 1 (*38*) (**Extended Data Figure 11a**). All mice were sacrificed within 30-40 minutes after the final behavioral session. *Ex vivo* brain slices containing the MEC and the hippocampus were then prepared to assess synaptic plasticity in MEC layer 2/3 while stimulating the hippocampus (*60–62*).

**Figure 6.**
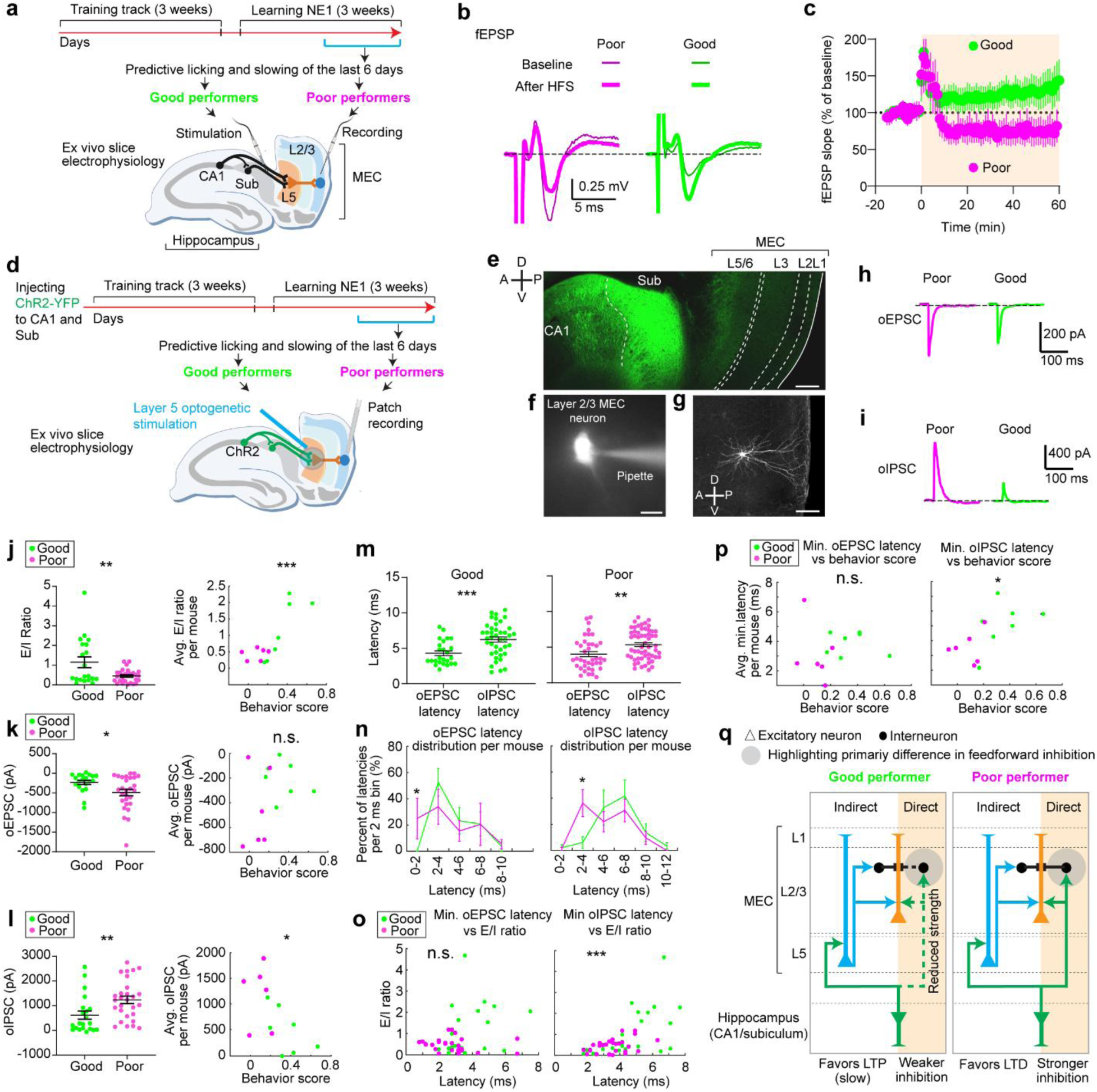
The microcircuits of layer 2 MEC in good and poor performers display different electrophysiological properties **a.** Schematic of the *ex vivo* electrophysiology experiments to recode fEPSPs in MEC layer 2/3 while stimulating hippocampal inputs. **b.** Representative traces of electrically evoked fEPSPs from poor (magenta) and good (green) performers before (thin) and after (thick) high-frequency stimulation (HFS). **c.** Time courses of the normalized fEPSP slope (mean ± SEM) from poor (magenta) and good (green) performers. HFS was given at 0 min. **d.** Schematic of optogenetic stimulation of axonal fibers from CA1/subiculum in layer 5 while performing patch recording from layer 2/3 neurons. **e.** Immunohistology of recoded slice showing Channelrhodopsin-2 expression in CA1 and subiculum. Axonal projections from the hippocampus to MEC layer 5 and layer 2/3 are reflected as dimmer fluorescence. Scale bar: 200 µm. **f.** Example a patched neuron. Scale bar: 15µm **g.** Confocal image of a recorded layer 2/3 neuron. Scale bar: 100µm **h.** Representative traces of optogenetically evoked oEPSCs in MEC layer 2/3 neurons, from a poor (magenta) performer and a good (green) performer. **i.** Similar to **h** but for oIPSC. **j.** Left: E/I ratio of individual neurons from good and poor performers. Right: Relationship between behavior score and averaged E/I ratio across neurons for each good and poor performer **k.** Similar to **j** but for oEPSC. **l.** Similar to **j** but for oIPSC. **m.** Comparison of oEPSC and oIPSC latency distribution for good (left) and poor (right) performers. **n.** Distribution of oEPSC (left) and oIPSC latencies (right) in good and poor performers, binned in every 2 ms. For each mouse, the number of latencies in each 2ms time bin were collected across all cells of the mouse and were normalized by the total number of latencies of the mouse. This produces “percent of latencies in each bin” per mouse. This percent in each 2ms time bin was compared between good and poor performers. **o.** Relationship between averaged minimum oEPSC (left) and oIPSC latency (right) and E/I ratio, for individual neurons in good and poor performers. **p.** Relationship between averaged minimum oEPSC (left) and oIPSC latency (right) across neurons of the same mouse versus behavior score of each mouse. **q.** Proposed circuit mechanism underling differential synaptic plasticity in MEC layer 2/3 of good and poor performers. For each performer group, “direct” and “indirect” indicate “direct pathway” and “indirect pathway” from the hippocampus, respectively. The reduced strength of the “direct pathway” in good performers primarily leads to reduced feedforward inhibition, favoring slow LTP induction through in the “indirect pathway”. The stronger inhibition in poor performers favors LTD. *p ≤ 0.05, **p ≤ 0.01, ***p ≤ 0.001, n.s. p > 0.05. Statistical test results are listed in Table S1. Error bars represent mean ± SEM.

We recorded extracellular field excitatory postsynaptic potentials (fEPSPs) from the MEC layer 2/3 while stimulating axonal fibers from the CA1 and subiculum (Fig. 6a). These fibers primarily convey hippocampal inputs to MEC layer 2/3 via MEC layer 5 (“indirect pathway”) (*63–66*), with a recent study also identifying a direct CA1 to MEC layer 2/3 pathway (“direct pathway”) (*67*). A 15-minute baseline fEPSP was recorded, followed by high-frequency stimulation (HFS) to induce synaptic plasticity (*68*). fEPSPs were monitored for 60 minutes post-HFS. The fEPSP slope changed after HFS (Fig. 6b). Good performers showed an increase in fEPSP slope (125 ± 18% from baseline), indicating long-term potentiation (LTP). Poor performers exhibited a decrease (67 ± 15% from baseline), indicating long-term depression (LTD) (Fig. 6c). These results suggest that hippocampal inputs are more likely to induce synaptic potentiation in MEC layer 2/3 of good performers compared to poor performers.

To explore the mechanisms behind synaptic plasticity differences in good and poor performers, we conducted whole-cell recordings from MEC layer 2/3 neurons while selectively stimulating hippocampal axonal projections using optogenetics. We virally expressed channelrhodopsin-2 variant H134R (*69*) in CA1 and subiculum neurons and prepared brain slices from VR-trained mice 4-6 weeks after virus injection (Fig. 6d and e). Mice underwent the same spatial learning task in N1 and were classified as good or poor performers (**Extended Data Figure 11b**). Brief blue light pulses were applied to the MEC layer 5 to stimulate inputs from CA1 and subiculum. Voltage-clamp recordings from layer 2/3 neurons (Fig. 6f and g) showed excitatory postsynaptic currents (oEPSCs), which were blocked by glutamate receptor blockers (NBQX and CPP) when holding the membrane potential at –60 mV (Fig. 6h). The same optogenetic stimulation also evoked inhibitory postsynaptic currents (oIPSCs), which were blocked by GABA receptor blockers (gabazine) when holding the membrane potential of the neuron at the reversal potential for glutamate receptor currents (0 mV) (Fig. 6i).

We calculated the ratio of oEPSC to oIPSC amplitude (E/I ratio) for each neuron, finding that E/I ratio was higher in neurons of good performers and correlated with learning performance (Fig. 6j**, and Extended Data Figure 11c** for behavior score calculation). While oEPSC amplitudes showed a small decrease in good performers compared to poor performers, this amplitude did not correlate well with their learning performance (Fig. 6k). In contrast, oIPSC amplitudes were significantly lower in good performers and negatively correlated with learning performance (Fig. 6l). We propose that in MEC layer 2/3 of good performers, there is reduced recruitment of both excitation and inhibition via the hippocampus-to-MEC projection, with a greater reduction in inhibition leading to a higher E/I ratio. This network state likely lowers the threshold for neural activation, increasing the likelihood of synaptic potentiation in good performers.

To identify specific synaptic pathways undergoing changes, we analyzed the latency of oEPSC and oIPSC response events relative to the optogenetic stimulation. Each event could have multiple latencies, reflecting multiple presynaptic inputs converging on the same postsynaptic cell (**Extended Data** Figure 12). After pooling individual latencies, we found that oIPSC latencies were significantly delayed compared to oEPSC latencies in both good and poor performers (Fig. 6m), indicating that inhibition requires the recruitment of more synapses than excitation. This finding is consistent with previous studies showing that while excitatory hippocampal projections reach MEC layer 2/3 through both “direct” and “indirect pathways”, these projections further synapse with interneurons to provide feedforward inhibition (*65, 67, 70*).

We observed that oEPSC latencies in both groups were mostly within 4 ms. However, good performers had a lower percentage of shorter latencies (0-2 ms) (Fig. 6n, left), suggesting that they received less excitation from the “direct pathway”. Meanwhile, oIPSC latencies of poor performers peaked around 2-4 ms and 6-8 ms, but good performers showed reduced latencies around 2-4 ms, and similar latency abundance at 6-8 ms (Fig. 6n, right). This reduced short-latency oIPSC in good performers indicates that they received less feedforward inhibition from the “direct pathway”. Thus, the weakening of the “direct pathway” from the hippocampus to MEC layer 2/3 of good performers could lead to the reduced strengths of oIPSC and oEPSC observed above (Fig. 6k and l).

Additionally, the minimal latency of individual oIPSCs per neuron, which reflected the timing of its fastest inhibitory inputs, positively correlated with the E/I ratio (Fig. 6o). The averaged minimal latencies of oIPSCs in individual mice also positively correlated with their behavioral performances (Fig. 6p). These correlations were not observed for oEPSC latencies. These findings suggest that delayed inhibition in good performers primarily contributes to their higher E/I ratio and better learning performance.

Together, *ex vivo* slice electrophysiology of MEC layer 2 neurons reveals a link between spatial learning and hippocampal-induced synaptic plasticity in the MEC. LTP in MEC layer 2/3 is specifically induced in good performers, potentially supporting the gradual stabilization of their grid cell activity. Our whole cell recording results suggest that this potentiation is facilitated mainly by the weakening of the “direct pathway” from the hippocampus to MEC layer 2/3 (Fig. 6q). While this pathway predominantly drives feedforward inhibition (*67*), our results suggest that the reduced inhibition in this pathway increases the E/I ratio, lowering the threshold for neural activation. This circuit feature favors LTP induction when good performers encounter a novel environment, leading to a gradually stabilized grid map. Thus, the LTP is polysynaptic as it is driven primarily by the “indirect pathway”, consistent with the delayed Hebbian plasticity mechanisms proposed by the Vector-HaSH model (Fig. 5). In contrast, stronger recruitment of the direct pathway in poor performers leads to stronger feedforward inhibition, favoring LTD and destabilizing grid cell activity.

## Discussion

Combining longitudinal two-photon calcium imaging and modeling, we uncovered the dynamics of grid fields at the timescale of cognitive map formation during spatial learning. The combined results provide a signature of a delayed Hebbian plasticity mechanism in shaping the map. In a novel environment, grid fields appeared immediately then shifted backward on a run-by-run basis within and across days. During successful but not unsuccessful learning, these fields gradually stabilized at the landmarks preceding them. The shifts partially reset daily, revealing a slower consolidation process requiring online experience. Population dynamics of co-modular grid cells remained on the same 2D toroidal manifold, with their trajectories on the torus slowly rotating, shifting and gradually stabilizing. The model showed that these dynamics can arise from a Hebbian plasticity mechanism in hippocampal-to-grid synapses with slightly delayed hippocampal activity. The time-delayed plasticity from hippocampus to grid cells potentially drives backward shifting grid fields. Overall, in imaging experiments and model, we showed that the plasticity process resulted in a map that became stabilized but more fragmented over time, with improved positional estimation. Finally, our *ex vivo* electrophysiology results demonstrated the association between hippocampus-driven synaptic plasticity in an indirect pathway to MEC with spatial learning performance. The link we find between grid cell stabilization, synaptic plasticity, and successful spatial learning further reinforces the role of grid cells and cognitive maps in memory.

Our findings augment the characterization of short time-scale grid cell dynamics, such as the observation of “one-shot” landmark-driven grid cell responses within a single day (*11*), and the very long-term dynamics of hippocampal representational drift over a month (*21*). The timescale of a few days is required for animals to fully learn a novel environment as reflected by their behaviors (*71–73*), and is also expected from the problem of self-localization and mapping (SLAM), which requires the bootstrapping of positional estimation from landmarks and path integration and the buildup of their associations (*25–28*). At this timescale, we find that maps require slow learning and plasticity over extended bouts of experience to properly stabilize and faithfully encode space. This is consistent with the observation of several-day timescale adjustments in the formation of a global grid cell map in landmark-free open fields (*29, 31, 74*). Intriguingly, in a compartmentalized 2D environment where doorways and interior walls serve as local cues, this several-day map acquisition timescale results in the gradual fusion of grid map fragments over time (*74*), while in our 1D environment with local landmarks, the circuit appears to retrench on fragmentation, accentuating the discontinuity at each landmark over learning. We hypothesize that this is because of the perceptual and exploratory differences in the two environments – the paths in our environment are unidirectional and the landmarks are strongly localized salient markers with no long-distance cues about the continuity of the space – and that bi-directional exploration should result in map fusion rather than increasing fragmentation.

Indeed, consistent with this hypothesis, backward field shifts are responsible for increasing map fragmentation in our experiments, and in a bidirectionally traversed environment, there is no consistent notion of “backward”. Additionally, hippocampal place fields shift backward under unidirectional 1D track traversal on a similar timescale (*24, 32–37*). As in our observations on grid cells, the degree of shifting decreases with experience (*24*) and there is a between-day partial regression in the shifts (*24*). This continuous shifting of place fields over days differs from the transient backward shift of CA1 place fields following burst firing during initial field formation, a hallmark of BTSP (*12, 75, 76*), though a recent study suggests that BTSP could also explain these continuous shifts (*77*). Our works suggests an alternate possibility: hippocampal cells in our model are driven by feedforward grid inputs rather than landmark cues over the majority of the track; thus, the properties of grid field shifts should be mirrored in result in corresponding shifts in hippocampal fields (data not shown). Thus our findings raise the possibility that hippocampal shifts on the timescale of days could be grid-cell-driven and due to Hebbian-like learning with a time-delay rather than due to STDP or BTSP (*77*).

The time-delay in our proposed plasticity dynamics could stem from synaptic eligibility traces(*78*), local recurrent connections in the hippocampus (*79*), or information processing delays due to the largely indirect inputs from the hippocampus to layer 2/3 of the MEC (*63–66*). While future studies are needed to fully investigate these mechanisms, our *ex vivo* slice physiology supports the latter. We found that CA1/subiculum stimulation induced synaptic potentiation in MEC layer 2/3 of good performers and synaptic depression in poor performers. This difference may be due to the weakening of the “direct pathway” from the hippocampus to MEC layer 2/3, which mainly drives feedforward inhibition (*67*). Reduced feedforward inhibition in good performers increases the E/I ratio and lowers the threshold for neural activation, favoring LTP induction via the “indirect pathway” and creating the time-delayed potentiation proposed by the Vector-HaSH model. Conversely, stronger inhibition through the “direct pathway” favors LTD. Additionally, a recent study showed that show that hippocampal stimulation can enhance synaptic responses in MEC layers 2/3 due to heterosynaptic potentiation of cortical inputs (*67*). Cortical inputs, which carry multimodal sensory information (*80*), might be enhanced in mice with extensive VR experience, further facilitating synaptic potentiation in good performers. The ability to induce LTP in mice that learned an environment indicates that the MEC circuit’s capacity to potentiate is not saturated, allowing continuous learning of new environments.

In summary, our *in vivo* calcium imaging, computational modeling, *ex vivo* electrophysiology, and behavioral analyses reveal that delayed synaptic plasticity from the hippocampus to the MEC underlies the stabilization and fragmentation of the grid map, highlighting a key mechanism in spatial learning. This mechanism may also extend to other memory processes involving the hippocampus-entorhinal circuit, which maps spatial and nonspatial domains defined by salient features during learning.

## Acknowledgement

We thank all colleagues in the Gu lab, Ila Fiete lab at MIT, and Veronica Alvarez lab for supporting the work. We thank Dr. Ben Sorscher from Surya Ganguli and Lisa Giocomo’s labs for his help on the population bump trajectory analysis and Dr. Kijung Yoon from Hanyang University for his help on the phase analysis of grid cell activities. This research was supported by the Intramural Research Program of the National Institutes of Health (NIH) to YG (ZIA NS009415), to VAA (ZIA MH002987 and ZIA AA000421), as well as support to IF from ONR award N00014-19-1-2584, NSF-CISE award IIS-2151077 under the Robust Intelligence program, ARO-MURI award W911NF-23-1-0277, the Simons Foundation SCGB program 1181110, and the K. Lisa Yang ICoN Center. The contributions of the NIH authors were made as part of their official duties as NIH federal employees, are in compliance with agency policy requirements, and are considered Works of the United States Government. However, the findings and conclusions presented in this paper are those of the authors and do not necessarily reflect the views of the NIH or the U.S. Department of Health and Human Services.

## Author contribution statement

Conceptualization: YG, IF

Methodology: YG, IF, VAA, LC, HS, TM, SC, LLD, FS, SOV

Investigation: LC, LLD, HS, FS, TM, YM, SOV, NWT, KC, KJZ, LA, SC, IF, VAA, YG

Visualization: LC, FS, SC, LLD Funding acquisition: YG, IF, VAA Project administration: YG, IF, VAA Supervision: YG, IF, VAA

Writing – original draft: LC, YG, SC, HS Writing – review & editing: YG, IF, VAA, SC

## Data availability

All data used for figure and analysis in the study is available at Zenodo, DOI: 10.5281/zenodo.16739877

## Code availability

All code used for figure and analysis in the study is available at Zenodo, DOI: 10.5281/zenodo.16739877. Code for Vector-Hash model are available at https://github.com/FieteLab

## Methods

### Animals

All animal procedures were performed in accordance with animal protocol 1524 approved by the Institutional Animal Care and Use Committee (IACUC) at NIH/NINDS. Dataset 1 (*38*) contains 8 male and 7 female GP5.3 mice (C57BL/6J-Tg(Thy1-GCaMP6f)GP5.3Dkim/J, JAX stock #028280 (*39*)). All mice were aged 4–8 months old at the time of the experiment. Dataset 2 contains 6 female and 4 male non-carriers of the strain B6; C3-Tg (Prnp-MAPT*P301S) PS19Vle/J (JAX stock #008169 (*81*)) crossed with GP5.3 mice. Mice were 7-10 months old at the time of the experiment. One male mouse and one female mouse were excluded from downstream analysis due to discontinued days in their imaging schedule. Hence in total 8 mice were analyzed here. All mice were maintained on a reverse 12-hr on/off light schedule. Slice recording experiment utilizes 10 male and 10 female wildtype mice aged 3-4 months. Five male and female mice had Channelrhodopsin-2 virally expressed in CA1 and subiculum.

### Microprism construction

Microprism construction follows the previously described procedures (*38*, *41*, *82*). A canula (MicroGroup, 304H11XX) was attached to one side of a circular cover glass (3 mm, Warner Instruments, 64-0720), and a right angle microprism coated with aluminum on the hypotenuse (1.5 mm, OptoSigma, RBP3-1.5-8-550) was attached to the other side of the cover glass. All attachments were performed using UV-curing optical adhesive (ThorLabs, NOA81).

### Microprism implantation surgery

All mice received surgeries as described in previous studies (*38*, *41*, *82*). Mice were anesthetized using a tabletop anesthesia system (VetEquip, 901806). Anesthesia induction was performed with 3% isoflurane and 1L/min oxygen, and anesthesia state was maintained using 0.5-1.5% isoflurane and 0.7L/min oxygen. Surgery was performed on a stereotaxic alignment system (Kopf Instruments, 1900). Mice body temperature was maintained around 37 °C using a homeothermic pad and monitoring system (Harvard Apparatus, 50-7220 F). After anesthesia induction, dexamethasone (2 mg/kg, VetOne, 13985-533-03) were administered by intraperitoneal (IP) injection for anti-inflammation purpose and slow-release buprenorphine (1 mg/kg, ZooPharm, Buprenorphine SR-LAB) was administered subcutaneously for pain relief. Exposed skull will receive antimicrobial wash by Enroflox 100 (10 mg/mL, VetOne, 13985-948-10).

Microprism implantation procedures followed the previous description (*38*). A 3 mm craniotomy was centered at 3.4 mm lateral to the midline and 0.75 mm posterior to the center of the transverse sinus (approximately 5.4 mm posterior to the bregma) at the left hemisphere (*38*). A durotomy was then performed over the cerebellum. Mannitol (3 g/kg, Millipore Sigma, 63559) was administered by IP prior to the durotomy. The microprism was inserted into the transverse sinus and sealed to the skull with Vetbond (3 M, 1469SB). The headplate was then mounted on the skull opposite the craniotomy, and the prism and headplate were then adhered to the skull with Metabond (Parkell, S396).

### Virtual Reality

A customized virtual reality (VR) setup was used for the experiment, as described previously (*38*). Mice were head-fixed onto an air-supported polystyrene ball (8-inch diameter, Smoothfoam) using a mounted headplate. The ball rotated around an axle which restricted its motion to forward and backward rotation, driving the VR environment along linear tracks without rotation. The VR environment was projected onto a dome screen filling the visual field of mice (270° projection). An optical flow sensor (Paialu, paiModule_10210) with infrared LEDs (DigiKey, 365-1056-ND) was used to record motion of the ball, and output the motion signal to a computer via an Arduino board (Newark, A000062). An approximately 4 mL water reward was provided via a lick tube at the reward location via a solenoid. A lick sensor connected to both the lick tube and headplate holder was used to detect mouse licking. A mouse licking the lick tube creates a closed circuit between the lick sensor, the lick tube, the mouse (from the tongue to the skull), the headplate (which directly contacts the skull), and the headplate holder. The solenoid and lick sensor were controlled using a Multifunction I/O DAQ (National Instruments, PCI-6229). Virtual environments were generated and projected using ViRMEn software (Princeton, version 2016-02-12) (*83*). Imaging and behavior data were synchronized by recording a voltage signal of behavioral parameters from the VR system using the DAQ. ViRMEn environments were updated at 60 Hz. The DAQ input/output rate was 1 kHz. The synchronization voltage signal was updated at 20 kHz. Final behavioral outputs were matched to the imaging frame rate (30 Hz, see below) for synchronization.

Environments were colored blue and projected through a blue Wratten filter (Kodak, 53-700) to reduce contamination of the imaging path with projected light. Virtual environments were one-dimensional (1D) linear tracks with patterned walls and patterned visual cues at fixed locations. At the end of the track, mice were immediately teleported to the start of the track.

### Two photon imaging

Imaging was performed using an Ultima 2P plus microscope (Bruker). A tunable laser (Coherent, Chameleon Discovery NX) with 920 nm excitation wavelength was used. Laser scanning was performed using a resonant-galvo scanner (Cambridge Technology, CRS8K). GCaMP fluorescence was isolated using a bandpass emission filter (525/25 nm) and detected using GaAsP photomultiplier tubes (Hamamatsu, H10770PB). A 16x water-immersion objective (Nikon, MRP07220) was used with ultrasound transmission gel (Sonigel, refractive index: 1.335985; Mettler Electronics, 1844) as the immersion media.

The anterior-posterior (AP) and the medial-lateral (ML) angle of the prism (i.e., the angle of the surface of the prism along to the AP or ML direction of the mouse) relative to the head-fixed position of the mouse were measured prior to the imaging. The headplate holder and rotatable objective angles were set daily to align the objective with the prism in the AP and ML direction, respectively, such that the objective was parallel to the prism surface. Black rubber tubing was wrapped around the objective and imaging window to prevent light leakage into the objective.

Microscope control and image acquisition were performed using Prairie View software (Bruker). Raw data was converted to images using the Bruker Image-Block Ripping Utility. In dataset 1, Imaging was performed at 30 Hz with 512 × 512 pixel resolution (corresponding to 500 µm × 500 µm). In dataset 2, dual plane imaging was used to capture two fields of view (FOVs) simultaneously (15Hz each with 512 × 512 pixel resolution, corresponding to 750 µm × 750 µm). Average beam power at the front of the objective was typically 70–115 mW. Imaging and behavior data were synchronized as described above.

### Behavior experiment for dataset 1

Behavior experiment paradigm for mice in dataset 1 was described in a published study (*38*). After surgery, mice were allowed to recover for 5 days. Mice were then trained in a 1D VR environment (familiar track 1, F1) and their anticipation of reward was quantified by predictive licking (PL) and predictive slowing (PS) metric. PL was measured as the percentage of licks that occurred prior to the reward among those in the whole track, except licks for reward consumption. PS was calculated as the percentile of deceleration prior the reward compared to those in the reset of the track(*38*). Mice were regarded as being familiar to F1 if they display stable reward anticipation (less than 5% change in PL and PS) for 3 days.

The mice then performed a 10-day spatial learning task in another 1D VR environment (Novel track, N1), which was a 1000-cm linear track with patterned walls and visual cues as shown in **Fig. 1c**. A reward was placed at the 890-cm position. During experiment, the 15 participating mice were separated into good performers and poor performers based on their PS and PL. PS and PL were calculated with different distance thresholds toward reward (PS: 50, 100, 150 and 200 cm distance threshold; PL: 20, 35, 50 and 75 cm distance threshold) for all 10 days in N1 and averaged across days, generating 8 values (4 from PS and 4 from PL) per mouse. All 15 mice were ranked based on each set of values from high to low. The mice that were consistently ranked in the lower half (i.e., with low performance values) for all 8 sets of values regardless of the chosen parameters were categorized as poor performers. The remaining mice were categorized as good performers.

### Behavior experiment for dataset 2

Mice were first trained in a 1 m environment to encourage running. Mice were then trained in a 6m environment (F2) with a single reward in a passive task (i.e., the mouse received a water reward as long as they reached the reward location) until they displayed significant reward prediction behaviors. They were further trained to perform an active learning task in the same track. For this task, mice were required to actively stop (defined as one second with speed less than 5 cm/s) within a 50 cm reward zone to receive a reward. Mice could only receive one reward per run. All mice underwent manual training, consisting of manually stopping the mouse in the reward zone each run, for the initial 3-4 days. If a mouse started to stop on its own, it was further trained without manual stopping with a shortened stop period (0.5s). Once a mouse stopped on its own with the shortened stop period, the full 1s stop period was applied. Additional manual training was performed periodically as necessary to reinforce correct behavior.

Once mice performed well on their own, as measured by a stable percentage of runs receiving reward (success rate), the mice were imaged in the F2 for several baseline days and then imaged in a N2 (6m with new landmarks and a 50 cm reward zone in a different location) for spatial learning and imaging across 10 days. PL was measured for reward prediction within 20 cm before reward delivery location, and PS was measured for reward prediction within 90 cm before reward delivery location. For the runs without reward delivery, PL was measured within 20 cm before the end of the reward zone, while PS was measured within 90 cm before the end of the reward zone. Averaged PS and PL across the last six learning days were calculated for each participated mice, and they were compared to the predefined PL threshold (22.2 percent) and PS threshold (65.7^th^ percentile) (*38*) to determine whether the mice is a good performer (*38*). The thresholds were obtained from dataset 1 and were the minima of averaged PL and PS of good performers across days 5 to 10 under the same distance threshold. A mouse was defined as a good performer if either of its averaged PL or PS passed the threshold.

### Behavior experiment for electrophysiology

Mice were first trained in a 1m environment to encourage running, and then were trained in a 10m environment (FE) with a single reward in a passive task described above. After 3 weeks of training, Mice performed 3-week spatial learning in another environment with different landmark settings and single reward (NE1). Averaged PS and PL across the last six days were calculated for each participated mice, and they were compared to the predefined PL threshold (22.2 percent) and PS threshold (65.7^th^ percentile) (*38*) to determine whether the mice is a good performer (*38*).

### Image processing

Imaging data of dataset 1 was down-sampled by taking the average of each consecutive block of 3 frames and processed as previously described using published MATLAB® scripts (*41*). Dataset 2 was similarly processed but by rolling average of consecutive 2 frames. Cross-correlation based rigid motion correction was performed to the imaging data. Identification of region of interests (ROIs) that could represent neuron footprints in dataset 1 was performed using principal component analysis combined with independent component analysis (*84*). ROI identification for dataset 2 was performed using Suite2p (*85*). To remove artifactual ROIs occasionally caused by light leakage, ROIs with an extreme non-circularity (the aspect ratio of the ellipse best fitting the shape of the ROI ≥ 3) were removed from further analysis. The fluorescence time course of remaining ROIs was then extracted. The fractional change in fluorescence with respect to baseline (ΔF/F) was calculated as (F(t) – F_0_(t)) / F_0_(t) (*82*). For each cell, significant calcium transients (ΔF/F) were identified using amplitude and duration thresholds, such that the false-positive rate of significant transient identification was 1% (*86*). A final ΔF/F including only the significant calcium transients was used for all further analyses. A speed threshold was further applied to ΔF/F and was calculated by generating a 100-point histogram of all instantaneous velocities greater than 0 and taking the value twice the center of the first bin (approximately 1% of max positive speed). The ΔF/F points when the mouse was moving below the speed threshold were excluded from following analyses.

### Cross-day common cell identification

With the anatomical footprints of detected cells in each day, we used an established probabilistic modeling methods described in previous studies (*38*, *87*) to identify the common cells across days. We first perform automatic pairwise rigid registration of the imaging sessions to identify common neuron pairs between each of the two imaging days. Manual registrations were applied to the imaging session pairs with failed automatic alignment. Pairwise registration results were combined to determine the common neuron across all 10 days of the spatial learning task (cross-day neuron). Due to the pairwise nature of the registration, a neuron in a day may be aligned to different neurons in the following days. Manual examinations were applied to these cases to select the optimal alignment results that display the highest level of anatomical footprint consistency across all the last day in familiar track and the 10 learning days in novel track.

### Classification of grid cells within a single day

The classification of grid cells within a single day was conducted based on their spatial fields of averaged activities across runs, as reported previously (*38*, *41*). Speed-thresholded ΔF/F for a cell was spatially binned along the track in 5 cm bins and averaged across runs. A gaussian window of 3 spatial bins was applied to the averaged ΔF/F trace to smooth the data. Spatial fields were identified by comparing the amplitude of original smoothed ΔF/F with a random ΔF/F distribution created by 1000 bootstrapped shuffled responses. Each bootstrapped shuffled response was generated by rotating the ΔF/F trace of the whole session so that for every time point of the recording, its track position was preserved but its calcium response was changed. The ΔF/F trace was rotated by starting the trace from random sample numbers chosen from the interval 0.05 × N_samples_ to 0.95 × N_samples_, where N_samples_ was the number of samples in the ΔF/F trace. A shuffled mean ΔF/F was calculated for each rotation. For each 5 cm bin, the p_value_ equaled the percentage of shuffled mean ΔF/F that was greater than or equal to the real mean ΔF/F. Therefore, 1-p_value_ equaled the percentage of shuffled mean ΔF/F lower than the real mean ΔF/F. A spatial field was defined as a region of at least 3 consecutive 5 cm bins (except that the fields at the beginning and end of the track could have 2 bins) that had a mean ΔF/F higher than at least 80% of 1000 shuffles at the corresponding bins (1-p_value_ ≥ 0.2). Additionally, at least 10% of runs were required to have a significant calcium transient in the spatial field.

Grid cells were further identified within each recording day based on the following criteria. (a) Grid cells need to have at least two spatial fields on the track. (b) Each grid cell must have more than L/(5w) transitions between in-field and out-of-field periods, where L represents track length and w represents mean field width of the grid cell’s response. (c) The widest field of the response must be smaller than 5w. (d) At least 30% of the bins must be assigned to either in-field or out-of-field periods. (e) The ratio between in-field and out-of-field mean ΔF/F to be > 2 (*41, 53, 88*).

### Classification of cue cells within a single day

To identify cue cells, cue scores of each cell on a specific day session was first were calculated based on a previously published method (*42*). The activity of the cell was first shifted to best match a cue template, which contained ones and zeros representing track areas with and without landmarks, respectively. A landmark zone was identified for each landmark by including the landmark itself and the surrounding region expanded by half of the landmark width on both sides. The correlations between shifted activity and cue template within individual landmark zones were calculated and were further averaged as cue score of the cell. 200 shuffled cue scores were generated for each cell using the same calculation but by randomizing the landmark location in the cue template. Shuffled cue scores from all cells across all FOVs and imaging days were pooled to generate an overall cue score distribution. A cell was identified as a single-day cue cell if its cue score on the day was above the 80^th^ percentile of the shuffle cue score distribution. While previous studies used a 95^th^ percentile threshold (*38*, *41*, *42*), an 80^th^ percentile threshold was used here to ensure the identification of all cells with landmark-matched activity. Cue scores of cells were calculated toward left landmarks, right landmarks and both side of landmarks. A cue cell could be identified based on any landmark type.

### Detecting cross-day fields on a run-by-run basis

We developed a four-stage algorithm to more faithfully detect activity fields of grid cells as they could shift on a run-by-run basis and also fail to appear in specific runs (“missed-field runs”) (*41, 88*). In general, run fields were first identified on a run-by-run (RBR) basis (Stage 1). Then we connect individual run fields in neighboring runs into cross-run field segments (Stage 2). Next, we connected the neighboring cross-run field segments into cross-day fields (Stage 3). Finally, we merge the cross-day fields with similar trajectories (Stage 4).

### Stage 1. Identifying spatial fields on a run-by-run basis (“run fields”)

The speed-thresholded ΔF/F for a cell was spatially binned along the track in 2.5 cm bins to generate a 2D spatial activity matrix representing the cell’s activation along track in individual runs. Run fields were identified on a run-by-run basis using the p_value_ based method described above (“**Classification of grid cells within a single day**”). In each run, 1-p_value_ of each bin were calculated by comparing the ΔF/F amplitude within the bin to a random ΔF/F distribution created by 1000 bootstrapped shuffled responses, generated by rotating the ΔF/F trace of the run from random sample numbers chosen from the interval 0.05 × N_samples_ to 0.95 × N_samples_, where N_samples_ was the number of samples in the ΔF/F trace of the run. In-field periods were defined as three or more adjacent bins (except at the beginning and end of the track where two adjacent bins were sufficient) whose 1-p_value_ ≥ 0.85. Bins with intermediate mean ΔF/F remained unassigned. For dataset 1, only run fields with ≤ 95 cm width were included in the following analysis. For dataset 2, only run fields with ≤ 150cm width were included as the run fields were generally longer in this dataset (**Extended Data Figure 3a, the second panel from the top**).

### Stage 2: Connecting run fields in consecutive runs into cross-run field segments

We connected the centers-of-mass (COMs) of run fields to generate the cross-run field segments. COM of a run field was calculated based on an established method(*24*)

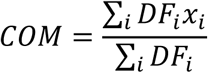

where *DF*_*i*_ is the ΔF/F of in-run field bin *i* and *x*_*i*_ is the index of bin *i* from the start of the track.

For each run field, we searched all the run fields in the next run to identify a closest subsequent run field, a COM of which was within a certain distance range (28 cm for dataset 1, 32 cm for dataset 2, and 10 cm for simulated data).

In the case when there are multiple qualified subsequent fields, we selected the subsequent field with the greatest spatial bin overlap with the current run field. If all subsequent fields had the same level of overlap, we selected the field with the smallest field width. If all remaining subsequent fields had the same width, then we do not connect the current run field with any of the candidates.

In the case when a run field was assigned to multiple run fields in the previous run (preceding fields), we selected the preceding field with the smallest COM distance and largest overlap with the current run field. If all preceding fields had the same level of COM distance and overlap, we selected the field with the smallest field width. If all remaining preceding fields have the same width, then we will not connect the current run field with any of the preceding fields.

After detecting all cross-run field segments by performing the steps above, all other run fields that were not assigned to any segments were kept for the next stage (Extended Data Figure 3a, the third panel from the top).

### Stage 3: Connecting multiple cross-run field segments to form a cross-day field

Next, we connected multiple cross-run field segments into cross-day field while permitting runs with missing fields in the middle. For two cross-run field segments, one started earlier (S1) and the other started later (S2) without overlapping runs, we calculated the distances between the averaged COM positions in the last three runs of S1 and the averaged COM positions in the first three runs of S2. S1 and S2 were connected if they showed the closest distance, which was within a distance threshold (28 cm for dataset 1, 38 cm for dataset 2, and 10 cm for simulated data), and the number of runs between the last run of S1 and the first run of S2 that did not have activity (gap) was no more than 7 runs.

In the case where an S1 had multiple qualified S2s, we selected the S2 with the largest number of runs, and the smallest gap toward S1. If multiple qualified S2s still existed after previous selection, we selected the S2 with the smallest track coverage (distance between the most backward COM to the most forward COM). If all S2s displayed the same number of runs, gap size, and track coverage, we did not connect the S1 to any of the S2s.

In the case where an S2 was assigned to multiple S1s, we selected the S1 with the largest number of runs, and the smallest gap toward S2. If multiple qualified S1s still existed after previous selection, we selected the S1 with the smallest track coverage. If all candidate S1s displayed the same number of runs, gap size, and track coverage, we did not connect the S2 to any of the S1s.

All putative cross-day fields were detected by performing the steps above. Cross-run field segments with more than one run and which were not assigned to any cross-day fields were kept for the following step (Extended Data Figure 3a, the fourth panel from the top).

### Stage 4: Connecting cross-day fields sharing similar trajectories

As stage 3 only examined the distance between the ending runs of S1s and onset runs of S2s, a potential problem was that the large field resets between the individual onset and ending runs (e.g., field regression or progression) could make an S2 disconnected with an S1, while their overall trajectories of the two segments were the similar, indicating that they could belong to the same field (**Extended Data Figure 3a, the fifth panel from the top**). To handle this case, for each of the two cross-run fields from stage 3, one appeared earlier (F1) and the other appeared later (F2) without run overlap, we linearly extended the trend of F1 to the runs occupied by F2, as well as extended F2’s trend to the runs of F1. F1 and F2 were connected if (a) the averaged COM location across runs between F1’s extended trend line and F2, as well as that between F2’s extended trend line and F1, did not exceed a distance threshold (28 cm for dataset 1, 34 cm for dataset 2, and 10 cm for simulated data; (b) the gap between F1’s ending run and F2’s onset run was no more than 7 runs.

In the case where an F1 had multiple qualified F2s for connection, we selected the F2 with the largest number of runs, and the smallest gap toward F1. If multiple qualified F2s still existed after the previous selection, we selected the F2 with the smallest track coverage. If all candidate F2s displayed the same number of runs, gap size, and track coverage, we did not connect any of the F2s to the F1.

In the case where an F2 was assigned to multiple F1s, we selected the F1 with the largest number of runs, and the smallest gap toward F2. If multiple qualified F1s still existed after previous selection, we selected the F1 with the smallest track coverage. If all candidate F1s displayed the same number of runs, gap size, and track coverage, we did not connect any of the F1s to the F2.

By following the steps above, we constructed the final set of cross-day fields. Only cross-day fields with more than 8 runs were included for the following analysis.

### Normalized run field count per run

Since two-photon imaging started after initiating a VR session and ended before terminating the session, there were track areas in the beginning and the end of the session without imaging data. Therefore, run field counts for the first and the last runs were normalized by the length of actual track area with imaging data. For the first run, we excluded the track area at the beginning with at least three consecutive bins without imaging data. For the last run, we excluded the track area at the end with at least three consecutive bins without imaging data. The run field counts were calculated as the number of detected run fields divided by the length of the remaining track for the first and last run. For all remaining runs, the run field counts were normalized by the true track length.

### Slope of cross-day fields

For each cross-day fields detected, we performed linear regression analysis (MATLAB® function polyfit) with the COM locations across run-fields, using run indices of run fields as the independent variable and COM locations as the dependent variable. The slope thus had the unit cm/run. Strictly non-shifting cross-day fields will have slope = 0. This method is similar to a previous analysis of shifting place fields in CA1 and CA3 (*24*).

### Determining shift direction of cross-day fields

We tested the slope of the cross-day field toward the baseline slope distribution generated by bootstrapping the field. We first sorted run fields of each cross-day field based on their COM locations to obtain their possible maximal shifts in forward and backward directions. Then we circularly rotate the sorted run fields and calculated the slope of each rotated fields to construct the bootstrap slope distribution. Two types of cross-day shifting fields were identified:

1. Cross-day backward fields: cross-day fields with slope smaller than or equal to the bottom 10% of the bootstrap slope distribution
2.Cross-day forward fields: cross-day fields with slope larger than or equal to the top 10% of the bootstrap slope distribution.

All remaining fields were regarded as cross-day stationary fields.

### Baseline percentage distribution of the cross-day field shift types

To determine whether a shift field type was significantly more abundant than baseline level, for each grid cell we randomized its activity by shuffling run fields within each detected cross-day field 100 times. With the shuffled activity map, we reapplied the cross-day field detection and calculated the fraction of each shift field type. The baseline fraction distribution was constructed from 100 shuffles (Extended Data Figure 3e). The original fraction is considered significantly higher or lower than the baseline if its value is >=95% or <=5% of the entire baseline distribution.

### Separating cross-day backward shifting fields to day fields

For each detected cross-day backward shifting field, we isolated its run fields within each day to constitute its day field of that day. Day fields needed to have at least 3 runs to be included in the following analysis. The center of the day fields (along track) was calculated by averaging the COMs of individual run fields of the day fields.

### Slope of day fields

Slope of a day field was similarly calculated as “**Slope of cross-day fields**” with the COM locations across the segment’s run fields, with run indices of run fields as the independent variable and COM locations as the dependent variable. The slope thus had the unit cm/run.

### Field center distances of adjacent day fields

Within a cross-day field, day fields on adjacent days were collected and their centers were calculated as the averaged COM of their run-fields. The absolute distance between the centers of day fields in adjacent days was calculated.

### Standard deviation of day fields

Standard deviation of a day field was calculated as the standard deviation of the COM locations of its run-fields.

### Normal distribution of day fields in learning days 7-10

This distribution was constructed by fields that ended within days 7 to 10. A cross-day field appeared between an adjacent landmark pair, containing a back landmark and a front landmark, which was determined based on its onset run field’s COM. Its onset distance to the back landmark was calculated as the percentage of the distance between the COM and the back landmark relative to the full distance (100%) between the landmark pair. Its end location within days 7 to 10 was defined as the averaged COM locations of all its run fields within these days. The distance of this end location to the back landmark (identified above for the onset run) was similarly converted to a distance percentage relative to the full distance of the same landmark pair above. This distance, termed “end percentage distance”, was negative or positive if the field passed or did not pass the back landmark, respectively.

Similar calculations were applied to all qualified fields, and their combined “end percentage distances” were grouped into 20% bins to obtain the probability density function (PDF). Then the envelope of the PDF was obtained by connecting the counts of each PDF bin. The peak location of the PDF was determined as the peak location of the envelop curve.

The hypothetical normal distribution *N*(μ, σ) was built such that μ equaled the peak location of the PDF, and σ equaled the absolute range of “end percentage distances” of all fields divided by 6, so that μ ± 3σ of the normal distribution covered >99% of the range (Empirical rule). With the two parameters, 20000 sets of random values from the distribution were simulated, with each set containing the same number of points as the original “end percentage distance”. The PDF of each random set was calculated with the same binning as in the true PDF.

The percentage of day fields passing the back landmark in each track bin were compared to the percentages of the 20000 simulated PDFs at the same location. The percentage of passing fields was defined as significantly smaller than the normal distribution if it was below the bottom 5% of the 20000 simulated percentages, and as significantly larger if it was above the top 95% of the 20000 simulated percentages.

### Grouping backward fields to assess their stabilization based on their distances to the back landmarks

For dataset 1 and simulated data, backward fields between two adjacent landmarks were grouped into 2 distance bins based on the distance between their median day field COMs and their back landmarks. For dataset 2, fields were grouped into 3 distance bins. Within each group, the slopes of day fields, the field center distance of adjacent day fields, and the standard deviation of day fields, were calculated using methods described above.

### Change in stabilization metrics for close and far fields between early and late learning

To assess the change between the stabilization parameters (the slopes of day fields, the field center distance of adjacent day fields, and the standard deviation of day fields) of early and late learning stages, for each field group (e.g., close or far fields) the averaged parameters of early learning stage (Days 1-3 for dataset 1 and simulated data, Days 1-2 for dataset 2) were subtracted from the parameters from late learning stage (Days 7-10 for all datasets).

### Identification of regression and progression events

All regression and progression events needed to happen between two day fields on adjacent learning days and with any of the two conditions: (a) the earlier day field’s last run reached the last run of the day, or (b) the later day field’s first run was present in the first run of the day. Both day fields needed to have more than 6 runs (for dataset 1), 3 runs (for dataset 2), and 9 runs (simulated data) to ensure the events were reliably detected. The larger numbers of runs for day fields were used for dataset 1 and simulated data to construct a more reliable “baseline distribution” to reveal their regression events (as described below, “Significant regression”), which had smaller magnitudes due to a multi-FOV imaging artifact (dataset1).

### Significant regression

A significant regression was determined by comparing the magnitude of the regression with the shift between all adjunct run fields of the pair of day fields that participated in the regression (“baseline distribution”). For dataset 2, the regression magnitude needed to surpass the top 85% of the baseline distribution to be considered significant. The percentile threshold for dataset 1 was 70% due to their significantly smaller magnitudes than those of dataset 2. The percentile threshold for modeling was set to 70% as the simulated environment matches dataset 1.

### Co-modular grid cells

Grid modules were identified based on the spacing of the cells of the same mouse(*51*). Since grid fields dynamically changed across days, we inspected the activity of each cell on all days (day 0 in F1 and days 1-10 in N1). The spacing of a cell was determined to be its smallest spacing of within-day spatial fields, which were identified based on the p_value_ of spatially averaged activity across all runs within a day in comparison to shuffles (*38, 41*), as described in “ Classification of grid cells within a single day”.

To further identify grid modules, we pooled the spacings of all cells from all FOVs in each mouse and identified peaks of the spacing distribution (evaluated by MATLAB® function “ksdensity”). The cells between two valleys were identified as co-modular cells.

### Run correlation of co-modular grid cells

For simultaneously imaged co-modular grid cells (with at least 6 cells per module), we first calculated pairwise correlations of the temporal activities of each cell pair in each run. We further calculated the correlation of the pairwise correlations of the exact same pairs on different runs (“run correlation” or “run corr.”).

### Correlation between trajectories of backward fields at different track distance

The correlation between backward fields was calculated between the segments of two backward fields from different co-modular grid cells that display activations across the same runs. The distance between the two backward field segments were estimated as the distance between the median run field COMs of the two segments. Correlation values were averaged within each 20-cm track bin. Pearson correlation was used to estimate the decreasing trend of the correlation across the track.

### Phase trajectory grid cell population

With a groups of simultaneously imaged co-modular grid cells, we first calculated the power spectral density (PSD) of individual grid cells on each learning day using Welch’s method (“pwelch” function in MATLAB®). Grid cell spatial activity was averaged across runs on each day for the calculation. We used WINDOW = hamming (512), NFFT = 2992, NOVERLAP = 461 (∼90% overlap between adjacent windows) and Fs=1000 (corresponding to the 1000 1-cm bins on the track) as input parameters to the “pwelch” function. The detrend operation was applied over the resultant PSD from 0.001 cm^-1^ to 0.150 cm^-1^ to reduce low frequency bias (using MATLAB®’s “detrend” function). The top three peaks (f1, f2, and f3) of the PSD were obtained from the “findpeaks” function in MATLAB®. Spatial frequencies of the three peaks were required to be larger than 0.006 cm^-1^ to avoid capturing low-frequency noise. To identify qualified co-modular grid cell populations for the analysis, we first averaged the spatial frequencies of the largest three peaks f1-3 for all cells within a day, and then averaged the results across days, producing a population-level f1-3 (avgf1, avgf2, avgf3) for the three frequency peaks. We included co-modular grid cell populations for further analysis if the difference between avgf3 and the sum of avgf1 and avgf2 was within 8% of avgf3, and the module contained more than 10 grid cells.

The phase of a grid cell on each day was determined based on the spatial frequencies of the three largest peaks (f1, f2, and f3) of its PSD, calculated from its averaged spatial activity across runs on the day. We further performed a 2992-point Fast Fourier Transform (FFT) to its averaged spatial activity. Phases of the FFT were obtained by applying the “angle” function of MATLAB® to the FFT result. The phases of f1, f2, and f3 (φ1, φ2, and φ3, respectively) were determined using the FFT phase at the same spatial frequency.

To visualize the population activity bump within the rhombus, we used an approach similar to a previous study(*11*): we first rounded the phase coordinates (φ1, φ2) of each co-modular grid cell to their closest integers, and then binned the coordinates into M*N tiles, where M is the largest value of all φ1s and N is the largest value of all φ2s of co-modular cells, respectively. Each binned coordinate (m, n) thus corresponded to a unique rounded (φ1, φ2). Multiple cells could have the same coordinates due to the rounding. We then sheared the binned coordinates (m, n) into a rhombus shape by:

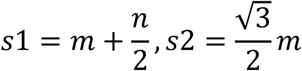

in which s1 and s2 are the coordinates in the rhombus. We represented each bin by the averaged calcium activity across all grid cells along 5-cm bins along the track, and Gaussian smoothed the activities for final visualization.

To calculate the coordinates of the population-level activity bump, we obtained firing-rate-weighted averages of the phases φ1, φ2, and φ3 across grid cells as functions of track position x:

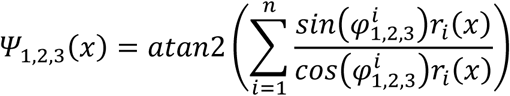

Here, *r*_*i*_(*x*) represents the firing rate for cell i at track location *x*, and Ψ_1,2,3_(*x*) represents the phase coordinates of the population phase trajectory at track location *x*. We use only Ψ_1,2_ to represent the phase trajectories based on the practice of a previous report (*11*). For visualization purposes, we unwrapped Ψ_1,2_ using the “unwrap” function in MATLAB® to avoid boundary conditions. Therefore, the population phase trajectory was represented across multiple phase rhombuses.

### Distance between population phase trajectories

The distance between the population phase trajectories on adjacent days were calculated as follows: between two trajectories in day n and day n+1, we first corrected phase coordinates of the earlier day’s starting point by subtracting or adding 2π if the offset between Ψ_1,2_ of the starting points of the two trajectories were larger than π or smaller than -π. The earlier day’s phase coordinates were corrected based on its starting point’s correction. We also sheared the trajectory coordinates Ψ_1,2_ to fit a rhombus shape:

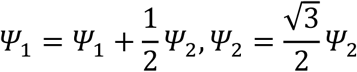

After correction, for each track point with phase coordinates Ψ_1,2_ on the trajectory of day n, we calculated its distance to the point at the same track location (“same-location point”) on the trajectory of day n+1. The distances across all track locations were averaged as the distance between the two trajectories.

### Rotation of population phase trajectory

To estimate the rotation between two bump trajectories, we subtract the starting point’s coordinates from the remaining coordinates of each trajectory to eliminate the effect of lateral shift between two trajectories. Rotation was then estimated from the distance between two trajectories across all track locations.

### Lateral shift of population phase trajectory

Lateral shift between start points of trajectories on day n and day n+1 was estimated using the distance between the starting points. Within each grid cell module, shifts between adjacent days were normalized by the averaged shift across all adjacent day pairs.

### Phase jumps on the population phase trajectory

To calculate phase jumps, we first calculated the phase trajectories using spatially binned (1 cm) activity within individual runs on each day. We obtained population phase trajectories of individual runs along the track. Each point had coordinates [Ψ_1_, Ψ_2_]. For two points along the track with 1 cm distance, their “phase jump” was calculated as the distance between their phase coordinates [Ψ_1_, Ψ_2_]. To define the within landmark phase jumps, we symmetrically expanded the landmark in both forward and backward directions along the track. For dataset 1 (Fig. 4), we expanded the landmark by 35 cm. For simulated data (Fig. 5), we expanded the landmark by 5 cm. Phase jumps within the expanded region were included in the “within landmark” jumps. All other phase jumps were considered as “between landmark” jumps. To focus the analysis on the landmark’s influence, we removed the jumps between the last two landmarks (863cm to 909cm), which included a water reward location, and 10 cm in the beginning and end of the track to exclude VR teleportation effects.

Influence index was calculated by subtracting the averaged between-landmark phase jump across runs from the within-landmark jump of each run, representing the general difference of “between landmark jumps” with individual “within landmark jumps”.

### Modeling

The core architecture of the model follows the recently published Vector-HaSH model (*54*) — we use a three-layer architecture, with a grid cell layer, a hippocampal layer, and a sensory layer. However, as compared to (*54*), we modify Vector-HaSH in a few key aspects: the construction of the grid cell network, the learning of hippocampus-to-grid synapses in the memory scaffold, and the addition of a second timescale of plasticity. Below, we describe the construction of the model, including these changes in detail.

### Vector-HaSH is implemented through three inter-connected layers

#### (1) Grid cells

Grid cells are simulated using a continuous attractor model of grid cells (*13*). Neurons are arranged on a lattice of N=128×128 neurons with periodic boundary conditions, with recurrent connectivity *W*_*GG*_ in the form of the mexican-hat interaction profile used in Burak & Fiete 2009(*13*). The network is initialized by driving the network for 250 timesteps with aperiodic boundary conditions, followed by 250 timesteps with periodic boundary conditions with unidirectional velocity inputs. This initialization protocol ensures that the initial state of the grid network has a stable triangular lattice of activity(*13*). Each timestep corresponds to 2 ms. At the start of each numerical experiment, the network is randomly reinitialized, resulting in a random grid phase at the beginning of the track.

#### (2) Hippocampal cells

The hippocampal layer receives inputs from sensory cells and is bidirectionally connected to the grid cell layer. As in Vector-HaSH, grid cells randomly project onto a hippocampal layer of size *N*_*H*_=2000 neurons using a projection matrix *W*_*HG*_. The elements of *W*_*HG*_are each independently drawn from a normal distribution with zero mean and unit variance. In Ref. (*54*), the return projections from the hippocampal layer to the grid layer, *W*_*GH*_, are pretrained through Hebbian learning to map hippocampal states to their corresponding grid states, and they are then held fixed. Here, we introduce a temporal offset in the learning between the hippocampal and grid layer, i.e., *W*_*GH*_ associates the grid state *g*(*t*) with an earlier hippocampal state, ℎ(*t* − τ). We describe this learning in greater detail below.

#### (3) Sensory cells

Sensory inputs, corresponding to cues observed along the linear track, are encoded as *N*_*s*_ =200 dimensional random vectors, with each element drawn independently from a normal distribution with zero mean and unit variance. Each landmark activates a subset of sensory cells corresponding to left or right cue cells. Boundaries at the start and end of the track are modeled as strong sensory inputs that drive both left and right sensory cell populations. We set the track length (1000 cm) and landmark locations (8 total) to match experimental conditions. These sensory inputs are fed into the hippocampal layer though sensory-to-hippocampus projections, *W*_*HS*_, that are heteroassociatively learned to map sensory states to place states.

The simulated animal navigates a linear track of 1000 cm featuring 8 landmarks placed to match experimental conditions. Motion is modeled over 5000 timesteps per run, with each timestep corresponding to 30 ms. The position along the track is updated corresponding to a uniform velocity. This velocity is also fed into the grid cell network, using path integration to update the grid phase. Specifically, the uniform velocity vector along the track maps to an input velocity along the grid cell network at an angle that is randomly drawn from [0,2π] and is held fixed for each numerical experiment. In addition to this rotation, time-independent Gaussian noise was injected into both (linear and angular) velocity and sensory inputs at each timestep.

The dynamics of Vector-HaSH proceed as follows:

In the absence of any sensory cues, grid cells path integrated the noisy velocity inputs to update the grid phases at each time step. The updated grid states project onto the hippocampal layer as

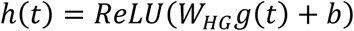

where *ReLU* is the rectifying function (*ReLU*(*x*) = *x* for *x* > 0 and *ReLU*(*x*) = 0 for *x* ≤ 0), and b is a parameter that controls the sparsity of the hippocampal representations in Vector-HaSH.

At every 20^th^ timestep (corresponding to 40 ms), grid cells receive additional inputs from hippocampal cells

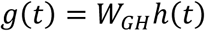

The grid cell network runs additional steps of the recurrent network dynamics without any velocity input to arrive at a stable steady state of the continuous attractor network dynamics.

During these hippocampus-to-grid updates, the hippocampus-to-grid synaptic weights are also updated using the temporal delayed learning rule previously described.

Since the grid activations are now continuous valued, we learn the hippocampus-to-grid associations using an online version of pseudoinverse learning(*89*) instead of Hebbian learning. The weights *W*_*GH*_ are thus updated through online pseudoinverse learning to associate *g*(*t*) and ℎ(*t* − τ). The time delay τ was set to 16 timesteps (corresponding to 32 ms) for all simulations.

To quantify the change in the magnitude of the hippocampus-to-grid synaptic weights, in Fig. 5b, we compute the Frobenius norm of the matrix of synaptic weights.

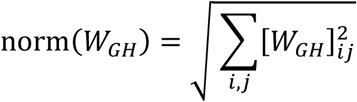

where [*W*_*GH*_]_*ij*_ refers to the (*i, j*)^*t*ℎ^ element of the matrix *W*_*GH*_, i.e., [*W*_*GH*_]_*ij*_ equals the synaptic weight from neuron *j* in the model’s hippocampal layer to neuron *i* in the model’s grid layer.

Note that this measure of the magnitude of synaptic weights quantifies how strong the hippocampal-to-grid synapses are, irrespective of their excitatory or inhibitory nature.

At a sensory cue location, the sensory inputs associated with the cue project into the hippocampal layer and update the hippocampal state. This new hippocampal state is in turn used to update the grid state, as well as the hippocampus-to-grid weights:

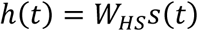

and

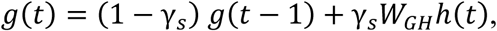

where γ_*s*_=0.1 is a parameter that controls the strength of the sensory cue information in determining the grid phase in Vector-HaSH. Note that this hippocampal update is implemented at all sensory cue locations, regardless of whether they occur at the periodic hippocampal updates of every 20 timesteps.

Before beginning the numerical simulation in a novel environment, the model is trained on a different environment for 20 runs, with identical track length but randomly drawn sensory cues, to simulate previous learning in familiar environments as in experimental conditions. This initial learning provides the initial conditions for the synaptic weight matrices in the novel environment. To capture the partial reset of grid fields at the start of each session, we additionally implement a second timescale of the learning dynamics, corresponding to synaptic consolidation. We implement this using two weight matrices: a fast-updating weight buffer that implements rapid online learning using pseudoinverse learning as described above, and a separate slow-updating weight matrix whose elements are given by the fast-updating weight matrix from τ_*C*_=4 runs prior. At the beginning of each new session (i.e., the start of a new day), the fast-updating synaptic weights are re-initialized to the permanent set of slow-updating weights, leading to a partial reset of the learned mappings.

To examine the role of plasticity across trials, we assessed the decoding of spatial position from grid cell population vectors. We constructed a Key Value Memory (KVM) network to learn the association with the grid cell population activity and spatial location. Specifically, we used a 120-cell subsample of the grid cell population activity vector (chosen to evenly sample cells by grid phase) as the keys, and position along the 1000 cm track (binned at 1 cm resolution) as the values. In testing, recall is performed by evaluating the similarity between the query population vector with the stored keys in memory. A softmax is then applied on these similarities, with a large inverse temperature set to 1000, effectively predicting position as the state that has the highest similarity. Additionally, we constrained the predictions of KV network to lie within a 300cm-sized neighborhood of the ground truth, to eliminate any highly discontinuous predictions of spatial location.

The decoding performance was evaluated every five trials using a sliding window: For each evaluation, a block of ten trials was considered for training the Key Value Memory network, which was then evaluated on the subsequent 10 trials. This resulted in 10^4^ training samples used to train the network for each evaluation (10 trials, each with 1000 positions). Decoding accuracy was assessed using the mean absolute error (MAE), calculated as the average absolute difference between predicted and true positions. In **Fig. 5p**, we then compared the decoding accuracy for the model as described above, with an equivalent model without any plasticity in the sensory-to-hippocampus and hippocampus-to-grid synaptic weights.

To further examine the role of plasticity, we repeated this analysis in Extended Data Figure 10 with a ten-fold increase in the magnitude of velocity and sensory input noise to accentuate the effect of noise reduction through plasticity. Specifically, we compare the model in three different conditions: (a) With plasticity in the synaptic weights, i.e., as used in the main text; (b) without any plasticity in the synaptic weights; and (c) with plasticity in the synaptic weights, but without any delay in the hippocampus-to-grid associative learning, i.e., setting τ = 0.

### Decoding performance of real grid cell data from dataset 1

Decoding analysis of real grid cell data from dataset 1was done using the same strategy as the decoding of simulated data mentioned above. Only FOVs with more than 10 cells were analyzed. Spatial activity within the first and last 150 cm of the track were removed from analysis due to unstable behavior(38). The decoding performance was evaluated every day using a sliding window: for each evaluation, a block of two days was used for training the KVM network, while the subsequent 2 days were used for the evaluation.

### Slice preparation

After animals were anesthetized with isoflurane, brain was quickly removed following decapitation in ice-cold solution containing (in mM) 90 sucrose, 80 NaCl, 3.5 KCl, 0.5 CaCl_2_, 4.5 MgCl_2_, 24 NaHCO_3_, 1.25 NaH_2_PO_4_, and 10 glucose saturated with 95% O_2_ and 5% CO_2_. For field recordings, horizontal hippocampal-entorhinal slices (300 µm) were prepared with the plate angled around 12 degree so that the caudal portion is raised compared to the rostral portion (ref) using a Leica vibratome 1200S, as described in (*60–62*). For whole cell recordings, sagittal slices (300 µm) were prepared with the plate also angled around 12 degrees (*60–62*) so that the caudal portion is raised compared to the rostral portion. Slices were then warm incubated (∼32℃) in artificial cerebrospinal fluid (ACSF) containing (in mM) 124 NaCl, 2.5 KCl, 2.5 CaCl_2_, 1.3 MgCl_2_, 26.2 NaHCO_3_, 1 NaH_2_PO_4_, 20 glucose, and 0.4 ascorbic acid saturated with 95% O_2_ and 5% CO_2_ for 20 minutes and then returned to room temperature until recording. For recording, slices were transferred to a submerged recording chamber and perfused at 2 ml/min with ACSF saturated with 95% O_2_ and 5% CO_2_, which was heated at 32℃ using an inline heater (Harvard Apparatus).

### Slice electrophysiology

Field recordings were done using patch pipettes filled with ACSF. Electrical stimulation of subiculum is achieved using bipolar electrodes while resultant fEPSPs signals were recorded from layer II and Layer V entorhinal cortex with two separate electrodes. The slope and amplitude of the fEPSPs were measured. A strong electrical stimulation of 4mA is used to induce a post synaptic response in layer V neurons. After a stable baseline of 15 minutes a high frequency theta burst protocol was applied to induce a long-term potentiation as described in Larson et al., 1986(*90*). Herein, the Theta burst protocol is composed of 10 bursts of 4 pulses at 100 Hz repeated 3 times every 20 second. Field potentials were acquired using Multiclamp 700B (Molecular Devices) were filtered at 1 kHz and acquired using Digidata 1550B and pClamp software (Molecular Devices). All data were analyzed using pClamp (Molecular Devices).

For optogenetic stimulation, whole-cell voltage clamp recordings were measured from layer 2 MEC neurons. Recordings were made using glass electrodes (2.5–3.5 MΩ) filled with an internal solution containing (in mM): 120 Cs-methanesulfate, 10 CsCl, 10 HEPES, 0.2 Cs-EGTA, 4 Na-ATP, 0.4 Na-GTP, 10 phosphocreatine, and 4.4 QX-314 (pH ∼ 7.3, ∼290 mOsm). 10 µM Alexa Fluo 594 was added to the internal solution to visualize cells after recording. Cells were held at −60 mV for EPSC and 0 mV for IPSC recordings. 10 ms light pulse of 470 nm light from pE-800 (CoolLED) was delivered on layer V every 20 s through 40x lens. Currents from Multiclamp 700B (Molecular Devices) were filtered at 1 kHz and acquired using Digidata 1550B and pClamp software (Molecular Devices). All data were analyzed using Igor Pro 7 (Sutter Instruments). E/I ratio is measured as the ratio between a cell’s oEPSC and oIPSC.

### Behavior score

Behavior score for an individual mouse is calculated as follows:

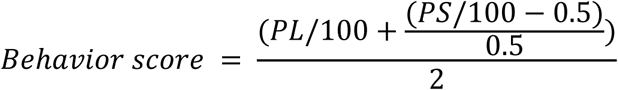

In which PS and PL are the predictive slowing and predictive licking of the mouse.

### oEPSC and oIPSC latencies

Onset latencies of excitatory or inhibitory responses were scored manually using the traces averaged from five traces at each condition. The latencies were calculated from the onset of optogenetic stimulations. Latencies from responses appear after the largest oEPSC and oIPSC level were not selected.

To compare the difference of latency distribution between good and poor performers, for each good and poor performer, the latencies from all individual cells were binned into 2ms bins and the number of latencies were counted in each bin. The latency counts were further normalized by the total counts of each mouse. The difference between the normalized latency counts within each 2ms bin were compared across good and poor performers.

### Statistics

Values with error bars are reported as mean ± standard error (SEM). Statistical analyses were described in figure legends. Two-tailed Mann-Whitney U test was used to test the difference between two unpaired value distributions. Two-tailed Wilcoxon matched pairs signed rank test was used to test the difference between two value distributions that constituted paired relationship. Two-tailed Student’s t test was used to test the difference between E/I ratio, EPSC and IPSC values of good and poor performers. One-sample t-test (two-tailed) was used to test the distribution of regression, progression, and within-day shifts toward standard normal distribution. Repeat measure two-way ANOVA (two-tailed) was used to test the overall difference between two metrics of across different days or runs. Pearson correlation was used to assess the level the increasing (or decreasing) trend of variables across learning. Significance was defined using a p value threshold of 0.05. In figures, we use different numbers of “*” to represent different ranges of p values: * for p ≤ 0.05, ** for p ≤ 0.01, and *** for p ≤ 0.001. “n.s.” indicates statistically non-significant (p > 0.05). For the analysis in Fig.3d bottom, Fig.1j and Fig.5d right if the percentile ≤ 5% or ≥ 95%, we used “*” to indicate the original value is significantly different from the shuffle distribution. All statistical test results are presented in **Table S1**.

## Extended Data Figures for

**Extended Data Figure 1.**
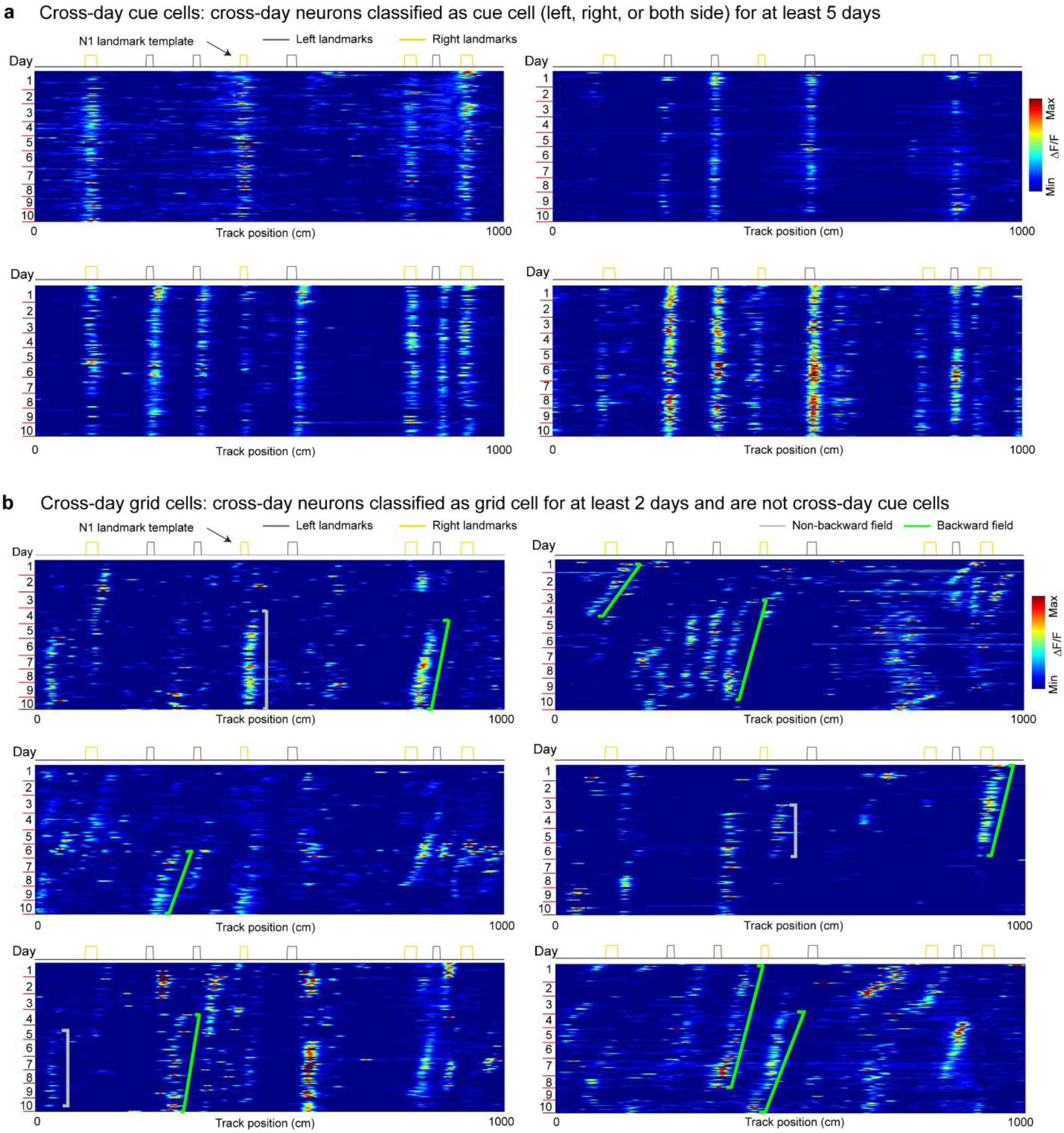
Example cross-day cue cells and cross-day grid cells, related to Figure 1 a. Example cross-day cue cells consistently responsive to right landmarks, left landmarks, and landmarks on both sides of the track across all learning days. b. Example cross-day grid cells displaying cross-day shifts.

**Extended Data Figure 2.**
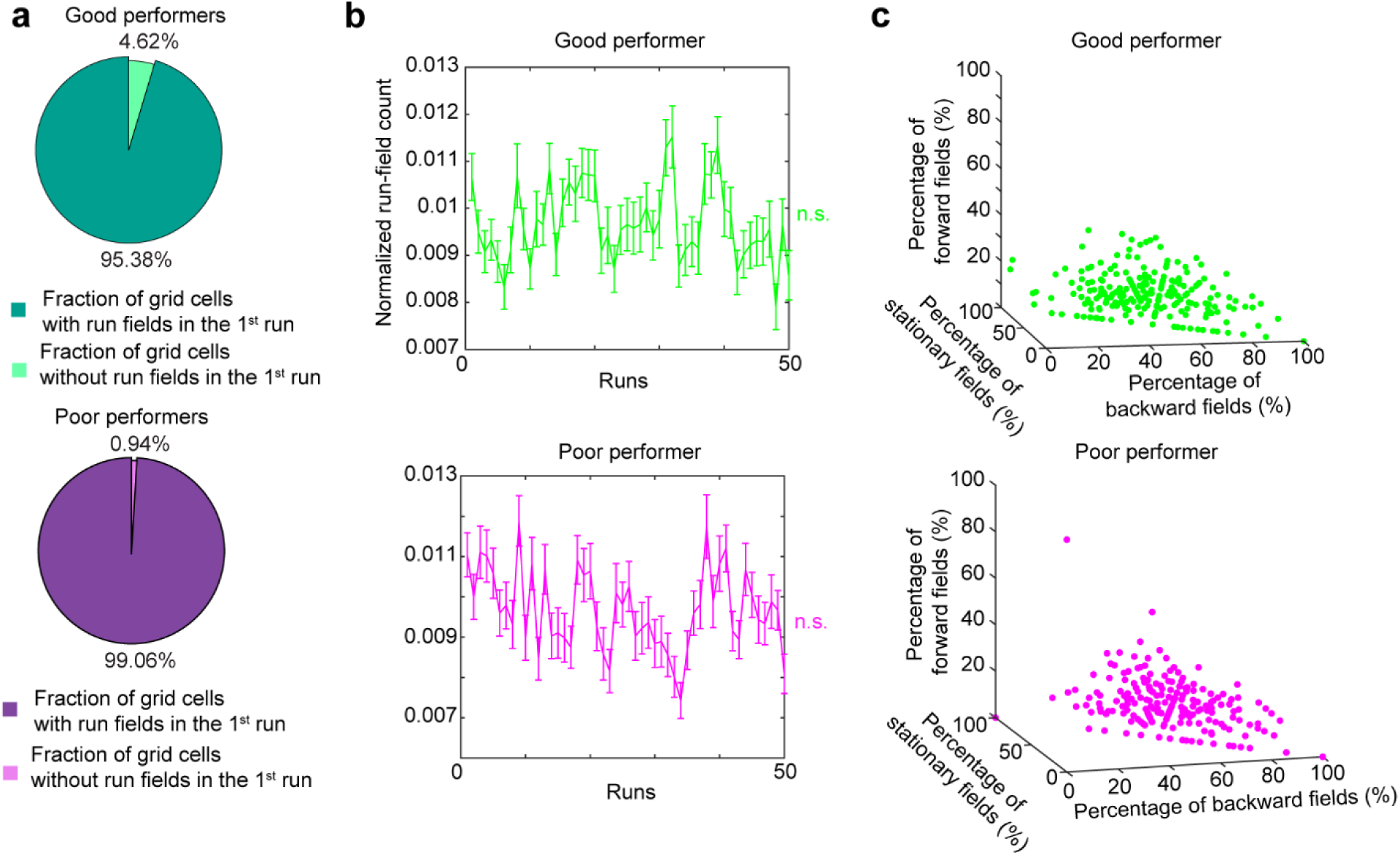
Grid cells display activity immediately upon exploring the novel environment and cross-day fields of different types, related to Figure 1 a. Percentage of grid cells recorded in the first field of view (FOV1) that display run field in the first run of the first novel environment learning day. b. Normalized run field count of first 50 laps for grid cells in FOV1. For first and last run, the number of fields were normalized by the track area with imaging data, for all remaining runs the number of fields were normalized by track length. c. Percentages of cross-day fields with different shift types for individual grid cells recorded in good and poor performers. *p ≤ 0.05, **p ≤ 0.01, ***p ≤ 0.001, n.s. p > 0.05. Statistical test results are listed in Table S1. Error bars represent mean ± SEM.

**Extended Data Figure 3.**
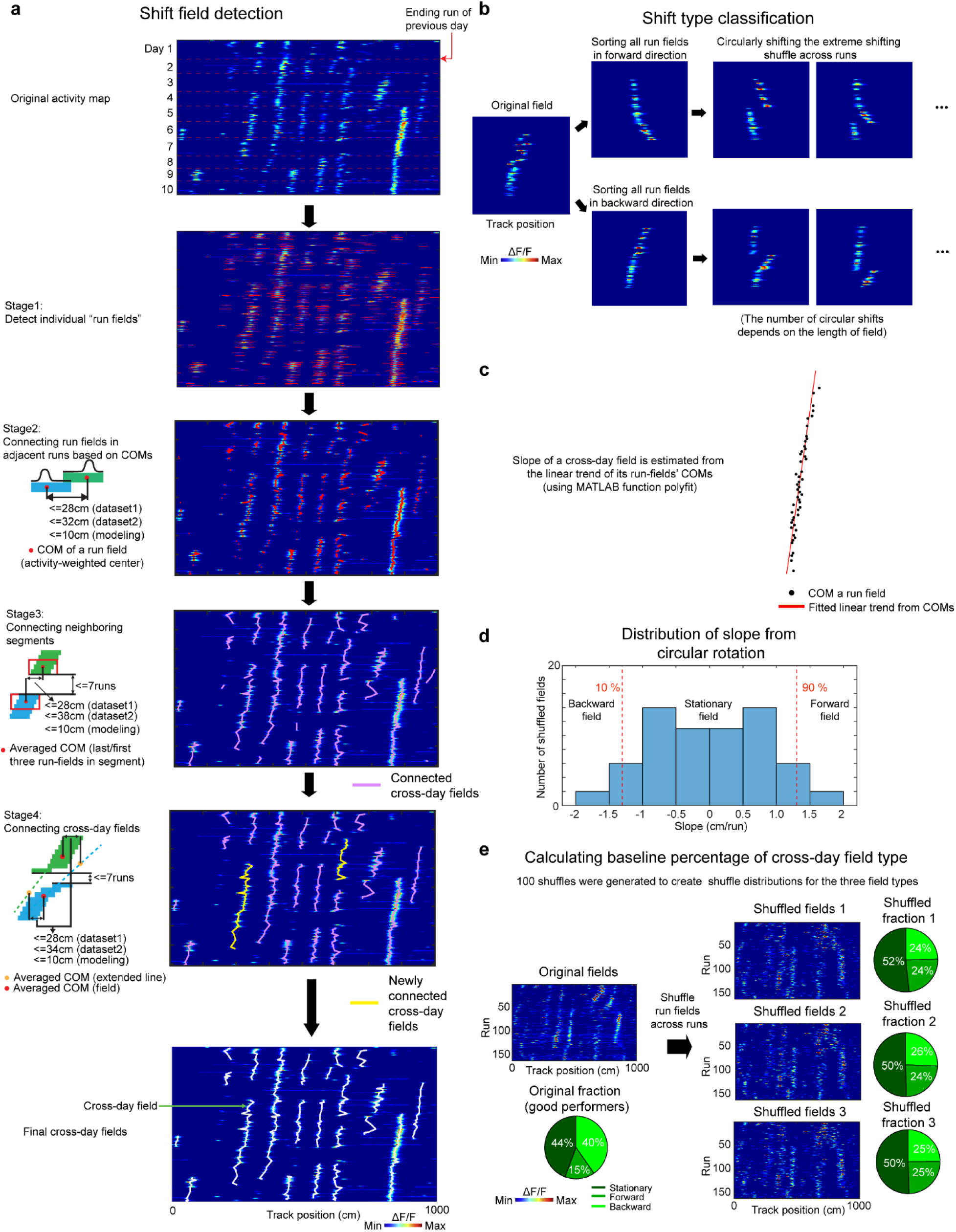
Schematic of cross-day field detection and shifting type classification, related to Figure 1 a. Schematic of cross-day field detection. After run field detection, cross-day fields were identified by first connecting closest run fields in adjacent runs, then connecting neighboring run field segments, and lastly connecting close-by cross-day fields sharing a similar trend and with averaged run field center of masses (COMs) at nearby track positions. b. Schematic of the cross-day field bootstrap shuffling for generating the baseline slope distribution. c. Schematic to determine the slope of the cross-day field using the COMs of individual run fields. d. An example bootstrap slope distribution used to determine the shifting type of a single cross-day field. All shuffled slopes of one field were pooled to construct a slope distribution, which was used to determine the significance and the direction of the field’s shift. e. Schematic of calculating the shuffle percentage for cross-day field types.

**Extended Data Figure 4.**
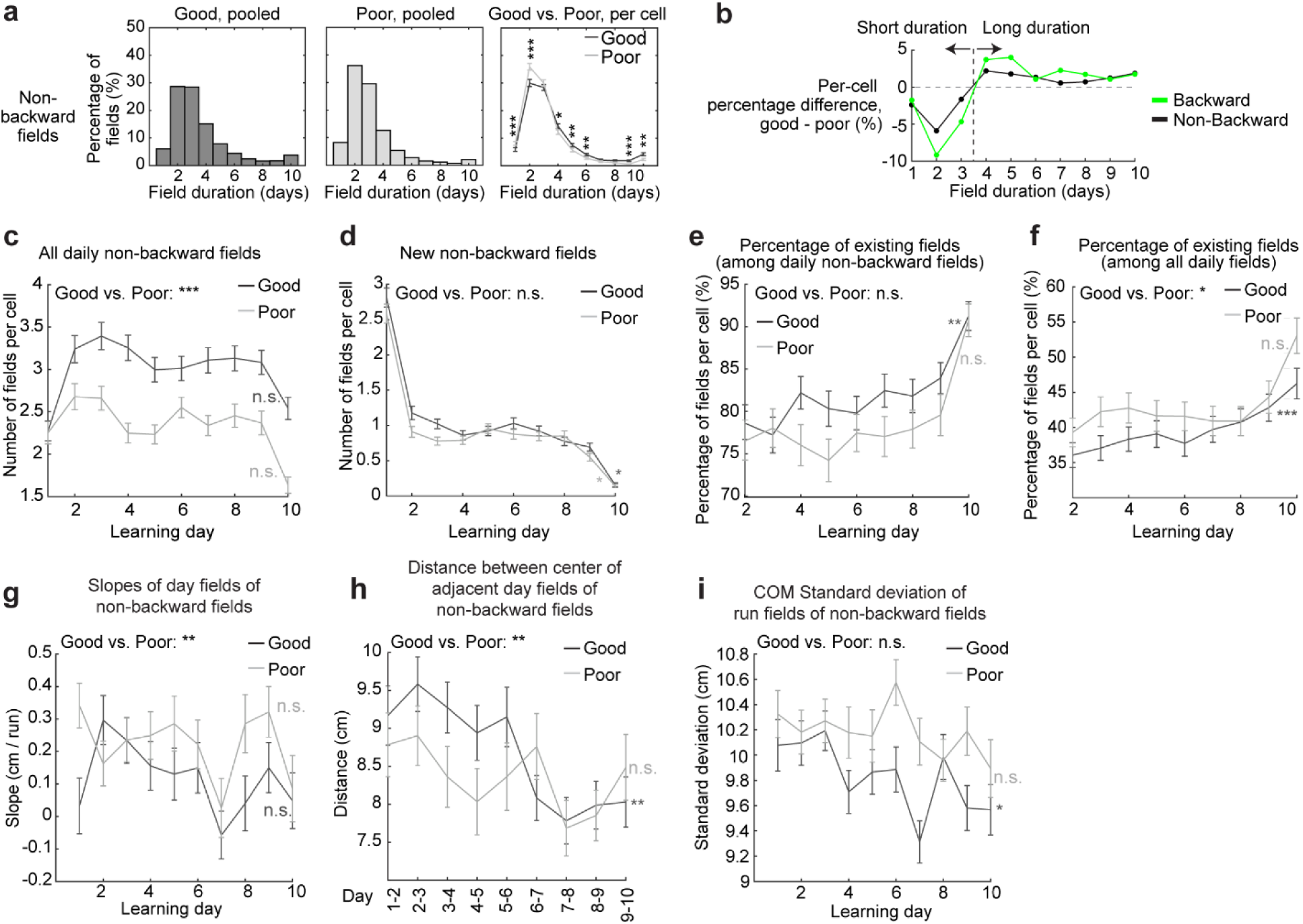
Stabilization analysis for the non-backward fields, related to Figures 1 and 2 **a**. Comparison of cross-day non-backward field duration (days) between good and poor performers. Left and middle: duration distribution of non-backward fields of good and poor performers. Right: Comparison between the percentage of non-backward fields with different durations per cell. **b**. Difference between good and poor performers in the percentages of their backward (green) and non-backward (black) fields with different durations. c. Numbers of daily non-backward fields per cell on each learning day. d. Number of new non-backward fields per cell appear on each learning day. e. Percentage of existing non-backward fields among daily non-backward fields per grid cell across learning days. f. Percentage of existing non-backward fields among all daily fields of different shift types per cell across learning days. g. Slopes of day fields of non-backward fields across learning days. h. Distances between the centers of adjacent day fields of non-backward fields. i. Standard deviation of COMs of run fields within day fields, for non-backward fields. *p ≤ 0.05, **p ≤ 0.01, ***p ≤ 0.001, n.s. p > 0.05. Statistical test results are listed in Table S1. Error bars represent mean ± SEM.

**Extended Data Figure 5:**
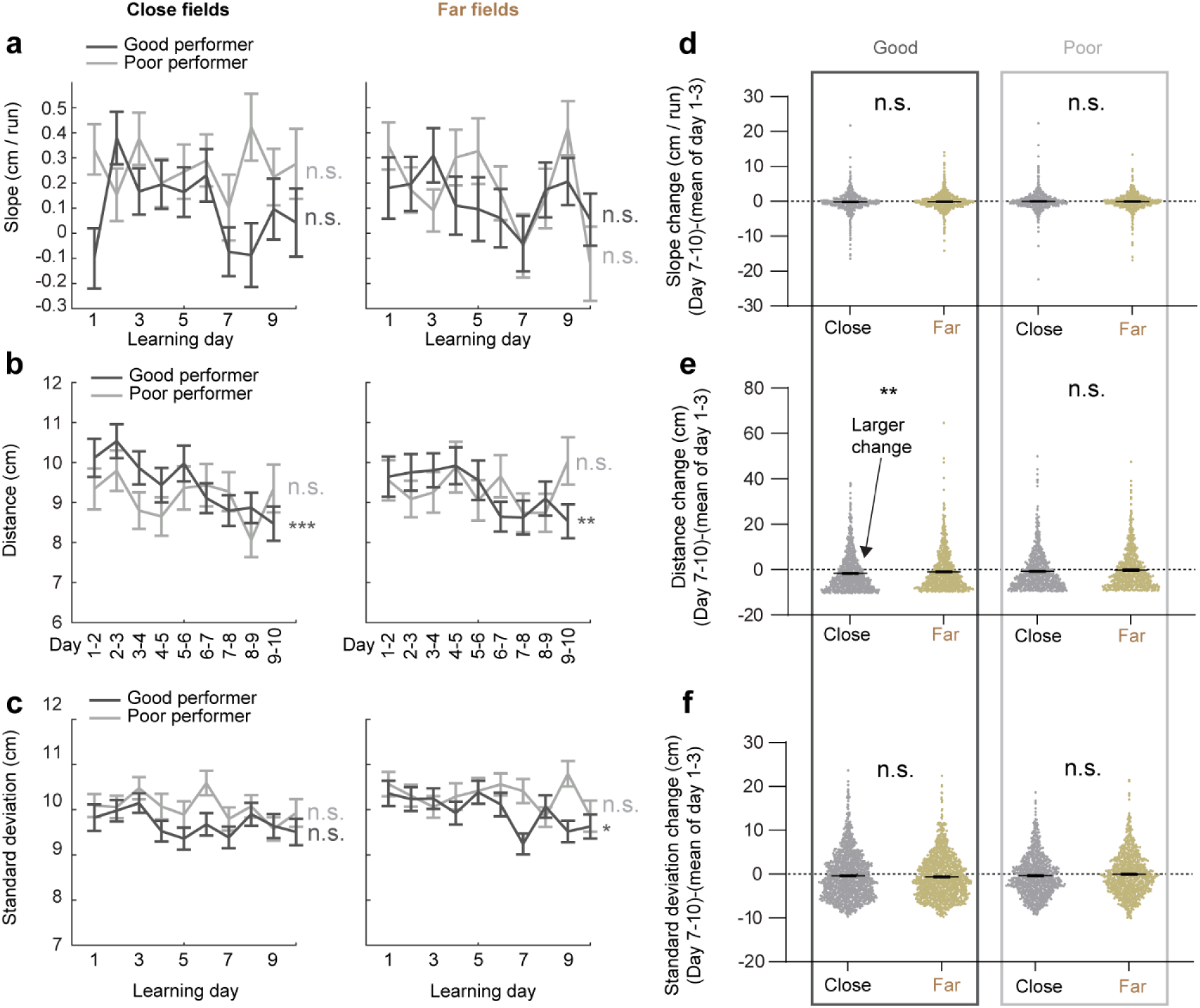
landmarks do not robustly affect the trajectories of cross-day non-backward fields, related to Figure 2. **a.** Slope of day fields across learning days for close and far non-backward fields. b. Center distance of adjacent day fields across days for close and far non-backward fields. c. COM standard deviation of run fields within day fields across learning days for close and far non-backward fields respectively. **d**. Change in day field slopes between days 1-3 and days 7-10, for close and far fields. Each dot is one day field on day 7-10. **e**. Change in field center distance between days 1-3 and days 7-10, for close and far fields. Each dot is one day field on day 7-10. **f**. Change in COM standard deviation of run fields between days 1-3 and days 7-10, for close and far fields. Each dot is one day field on day 7-10. *p ≤ 0.05, **p ≤ 0.01, ***p ≤ 0.001, n.s. p > 0.05. Statistical test results are listed in Table S1. Error bars represent mean ± SEM.

**Extended Data Figure 6:**
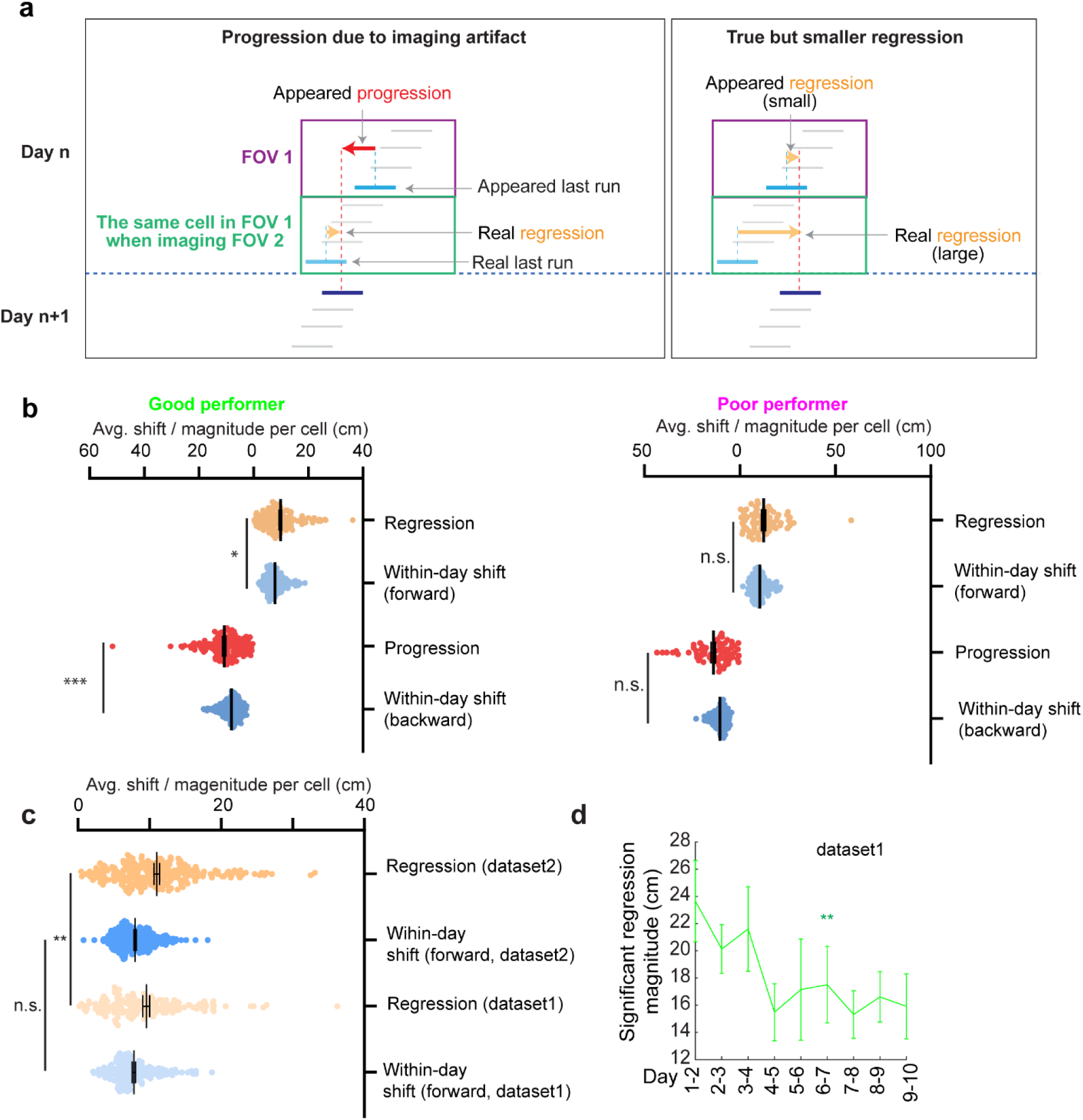
How dataset 1’s imaging scheme affect the investigation of forward reset, related to Figure 3 **a**. Schematic of how the imaging protocol of dataset 1 could affect the magnitudes of progression and regression events, respectively. Left: progression could be artificially created due to the missing data of FOV1 when FOV2 was imaged in the second half of the imaging session. Right: the magnitude of regression could be reduced due to the same reason. b. Comparison of the magnitudes of progression and regressions with within-day adjacent run field shifts in the same direction. **c.** The comparison between regression magnitudes in datasets 1 and 2, and the within-day adjacent run field shift in the forward direction, indicating that the regression magnitude was larger in dataset 2 while the within-day field shifts were comparable in the two datasets. d. Magnitudes of significant regression during learning in dataset 1.The significant regression is determined as those above 70^th^ percentile of within-day adjacent run field shift. the 70^th^ percentile was used based on the regression-magnitude-scaled percentile of dataset 2 (significant regression percentile for dataset 2: 85^th^, average forward regression shift of dataset 2: 10.984cm; average forward regression magnitude of dataset 1: 9.503 cm. Scaled significant regression percentile for dataset 1 = 85 × (9.503/10.984) = 73.539 ≈ 70). *p ≤ 0.05, **p ≤ 0.01, ***p ≤ 0.001, n.s. p > 0.05. Significant p values are listed in Table S1. Error bars represent mean ± SEM.

**Extended Data Figure 7:**
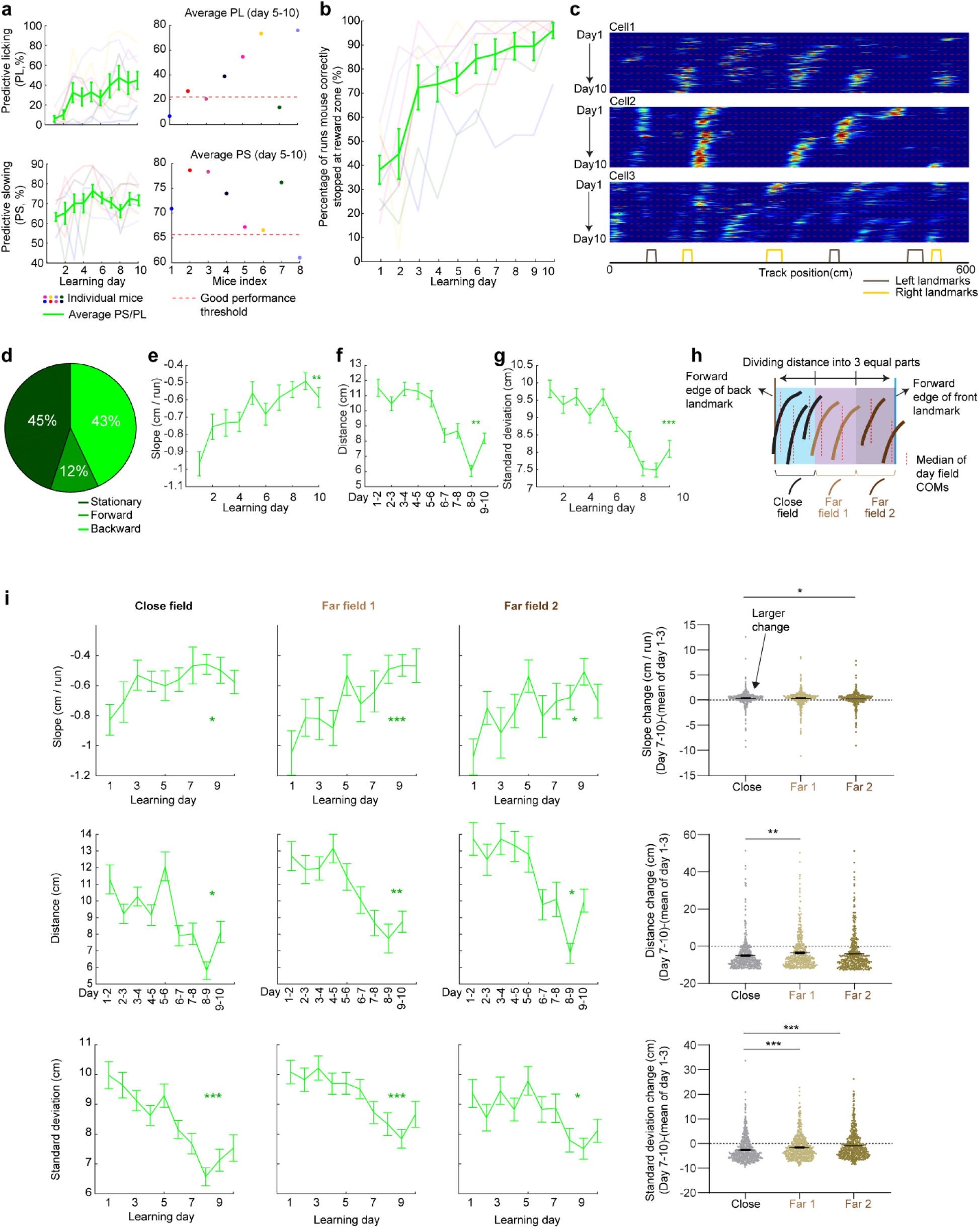
Grid cell dynamics in dataset 2 produce cross-day backward fields similar to dataset 1, related to Figure 3 a. Determining good performers in dataset 2.Predictive licking (PL) and predictive slowing (PS) of mice were collected for all the mice during learning. Averaged PS and PL of the last 6 learning days were calculated and compared to the previously reported PS and PL threshold for good performers (PL threshold: 22.2percent, PS threshold: 65.7^th^ percentile), as conducted previously (38). A mouse needed to have both averaged PL and PS below threshold to be classified as poor performers. Otherwise, the mouse was identified as a good performer. Therefore, all mice were classified as good performers. b. Percentage of runs when mice stopped within the correct reward zone across learning. c. Example cross-day fields of grid cells. d. Fraction of shifting types for all cross-day fields. e. Slopes of backward day fields across learning days. f. Distances between the field centers of backward day fields in adjacent days. g. COM standard deviations of run fields within backward day fields across learning days. h. Schematic of defining the close and far fields toward the back landmark for dataset 2. Areas between a landmark pair were divided into 3 equal parts, and “close fields”, “far fields 1”, and “far fields 2” were defined if their median field centers fell into specific parts. i. Stabilization of close fields, far fields 1, and far fields 2 by landmarks, reflected by their slopes, distances of adjacent day fields, and COM standard deviations of run fields across learning, similar to Fig. 2q-v. *p ≤ 0.05, **p ≤ 0.01, ***p ≤ 0.001, n.s. p > 0.05. Statistical test results are listed in Table S1. Error bars represent mean ± SEM.

**Extended Data Figure 8.**
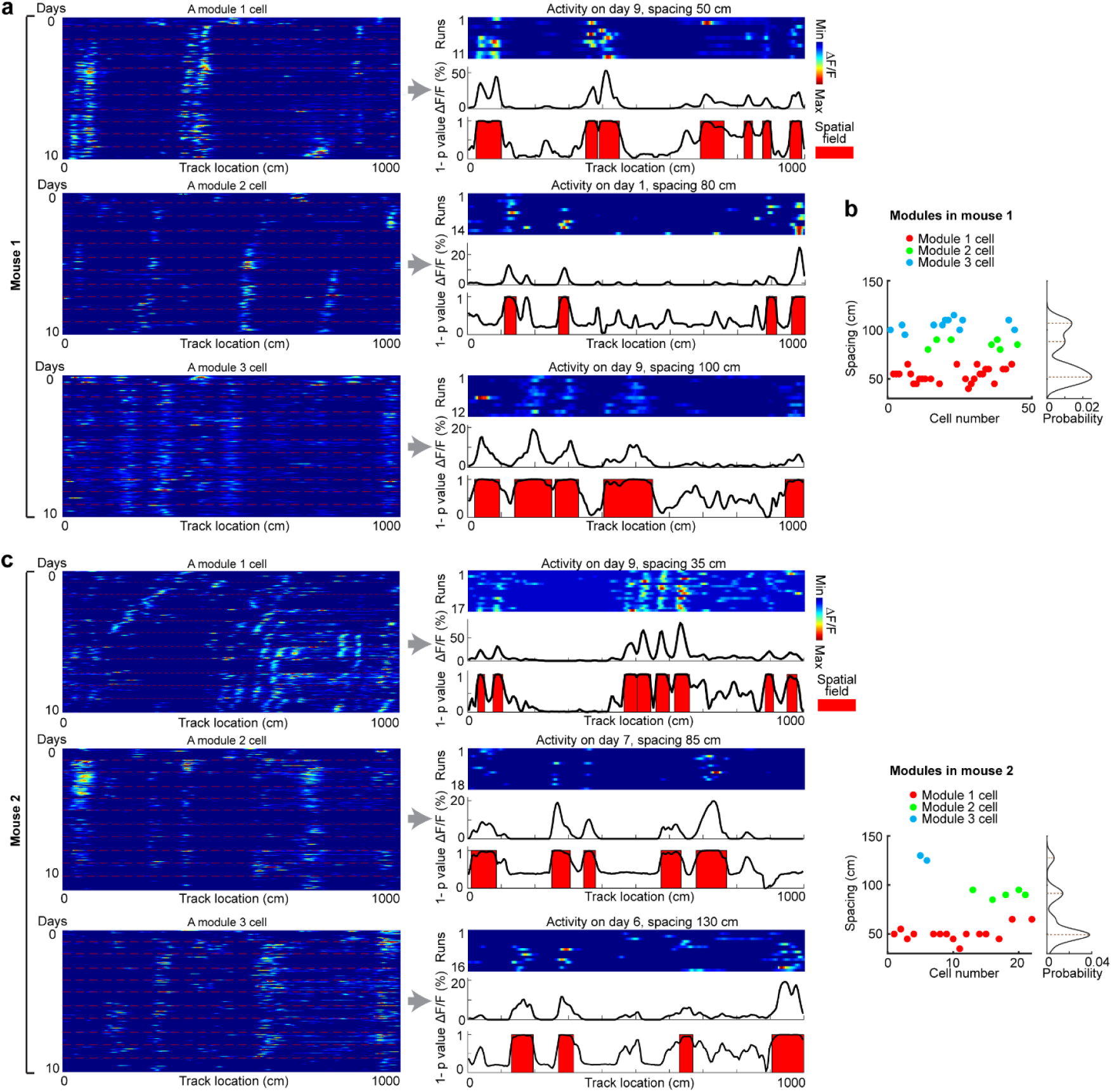
Identification of grid modules, related to Figure 4 Grid modules were identified based on the spacing of the cells within individual days. Since grid fields dynamically changed across days, we inspected the activity of each cell across all days (day 0 in F1 and days 1-10 in N1). The spacing of a cell was determined as its smallest spacing of day fields, which were identified based on the p values of spatially averaged activity (across runs) of the cell within a day in comparison to shuffles (*41*). **a**. Example cells for three modules in mouse 1. For each cell: Left: cross-day activity of the cell. Right: activity of the cell on a particular day, during which the cell showed the smallest spacing. Top: run-by-run activity heat plot. Middle: spatially averaged ΔF/F. Bottom: 1-p_value_ of the activity at each spatial bin. Higher 1-p_value_ corresponds to higher activity of the cell compared to shuffle. Spatial fields are highlighted in red. b. Grid modules of all cells in mouse 1. This mouse has three modules based on the spacings. c and d. Similar to a and b but for mouse 2.

**Extended Data Figure 9:**
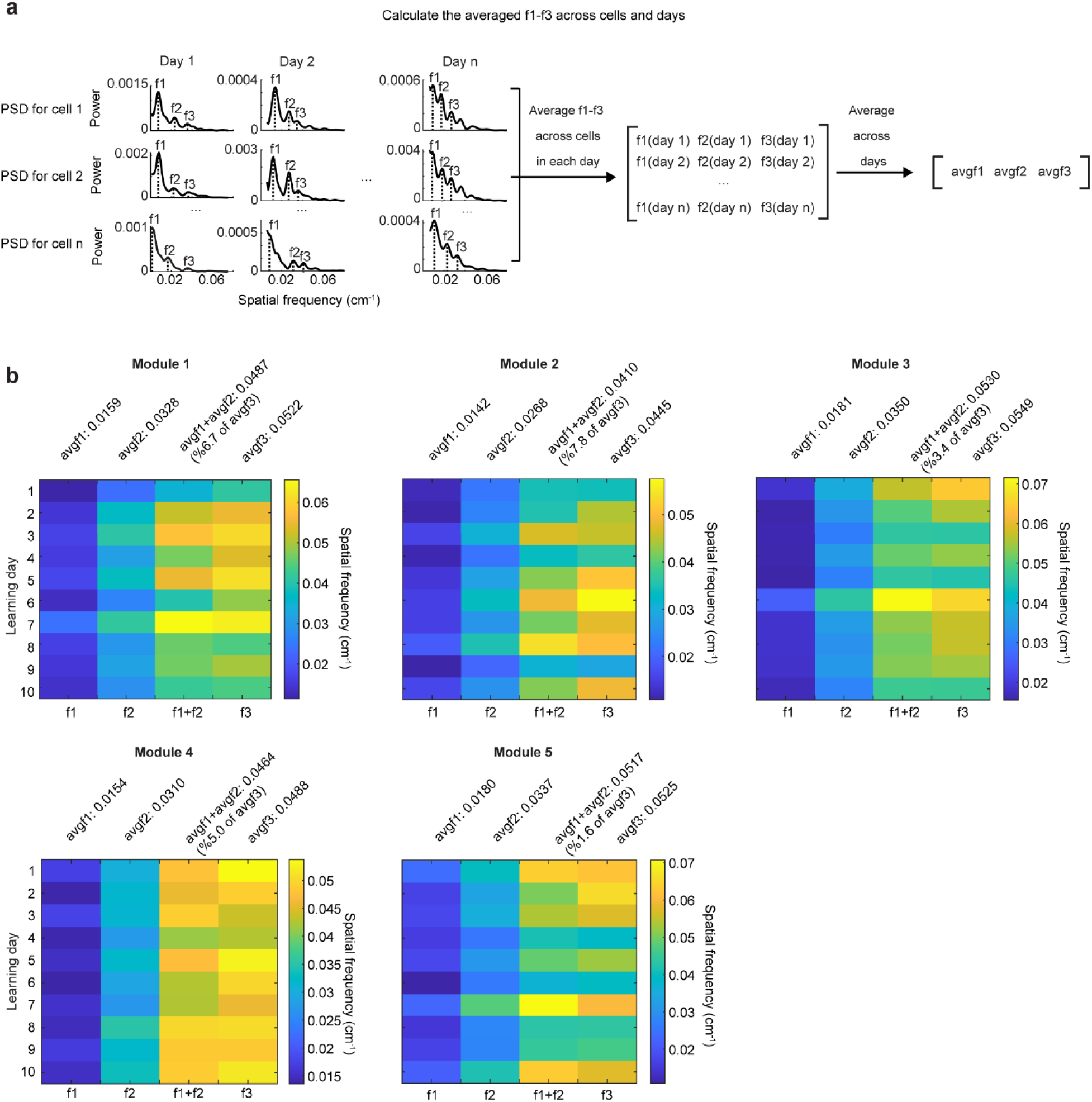
Spatial frequency feature of the selected grid cell modules, related to Figure 4 a. Schematic for calculating the avgf1-3 (averaged f1, f2, f3 across days). Within each day, the spatial frequency of the three largest peaks of each co-modular grid cell’s PSD (f1-3) were averaged separately. Following this, the population per day f1-3 were further averaged across days to obtain the final avgf1-3. b. Per day f1-3 for the selected five grid cell modules. For all qualified modules, the absolute difference between their avgf1+avgf2 and avgf3 must be smaller than 8% of avgf3.

**Extended Data Figure 10:**
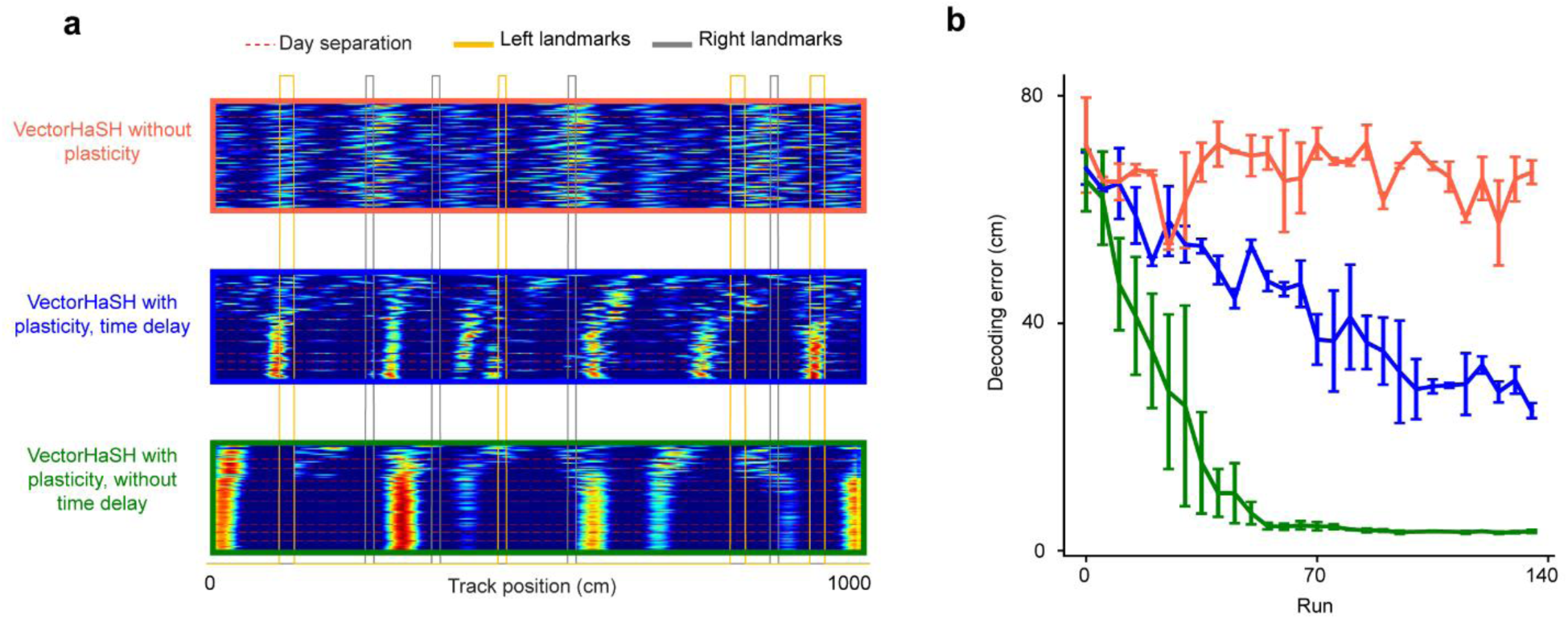
Decoding error of the model in the absence of plasticity or time-delays, related to Figure 5 a. Run-by-run cross-day activity of randomly sampled simulated grid cells in each of the three conditions: without any plasticity (orange), with plasticity at sensory-to-hippocampus and time-delayed plasticity hippocampus-to-grid synapses (blue), and with plasticity at sensory-to-hippocampus and hippocampus-to-grid synapses without any time delay (green). b. Location decoding error of simulated grid cells from the model in three conditions in a.

**Extended Data Figure 11:**
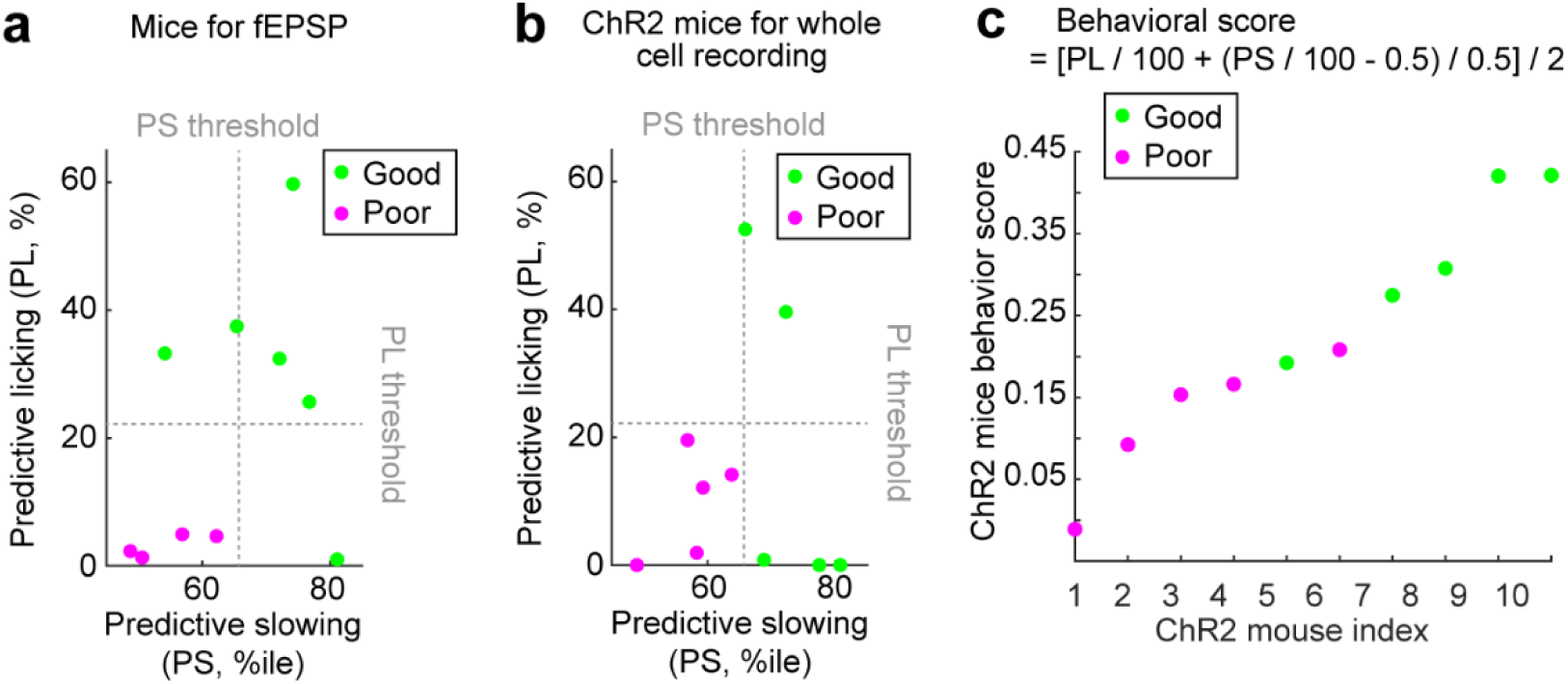
Behavior of mice in the electrophysiology experiment, related to Figure 6 a. Predictive licking (PL) and predictive slowing (PS) metric for mice participated in the fEPSP measurement experiment. Averaged PS and PL of last 6learning days in novel environment were compared to reported PS and PL threshold for good performers (PL threshold: 22.2percent, PS threshold: 65.7^th^ percentile) (38). Mice with both averaged PL and PS below thresholds were classified as poor performers b. Same as a but for mice participated in the optogenetic stimulation experiment c. Behavior score calculated from PL and PS of mice participated in optogenetic stimulation experiment. The score is represented by values from 0 to 1 based on the formula on top.

**Extended Data Figure 12:**
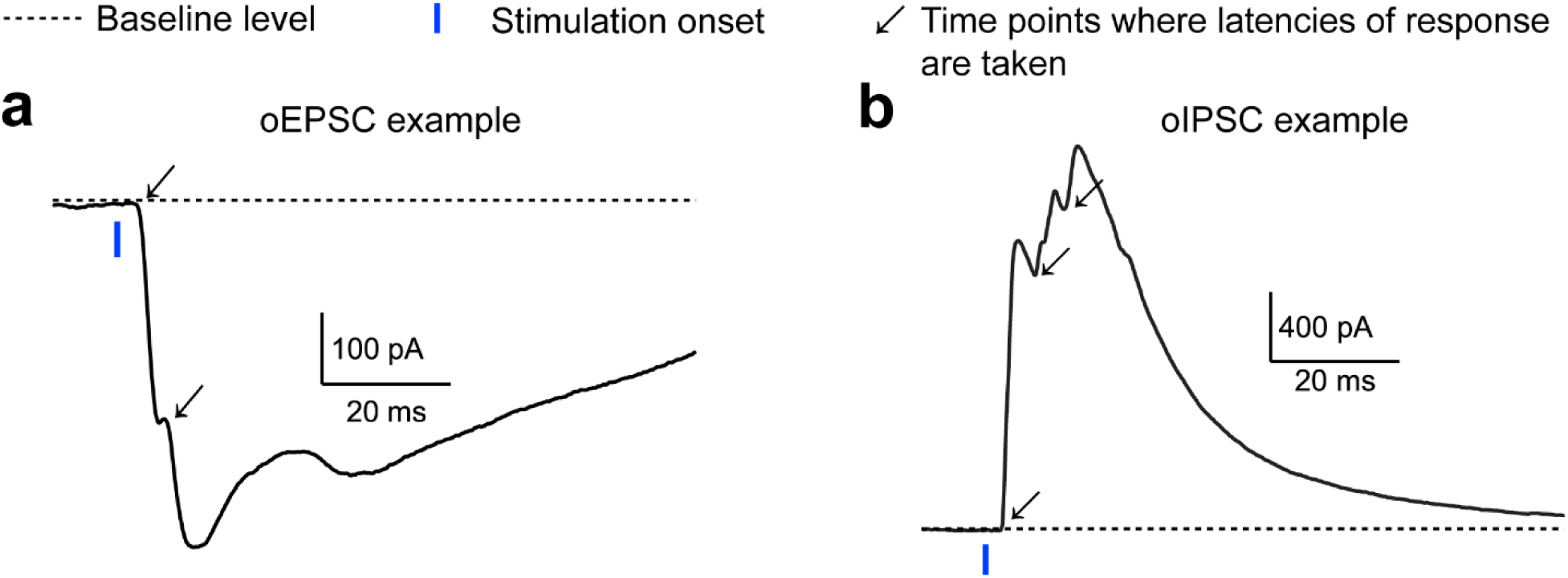
Multiple latencies within oEPSC and oIPSC events, related to Figure 6 a. Example oEPSC trace display multiple latency responses. Responses were labeled with arrow. b. Similar to a but for example oIPSC trace.

## Supplementary Materials for

**Table S1:**
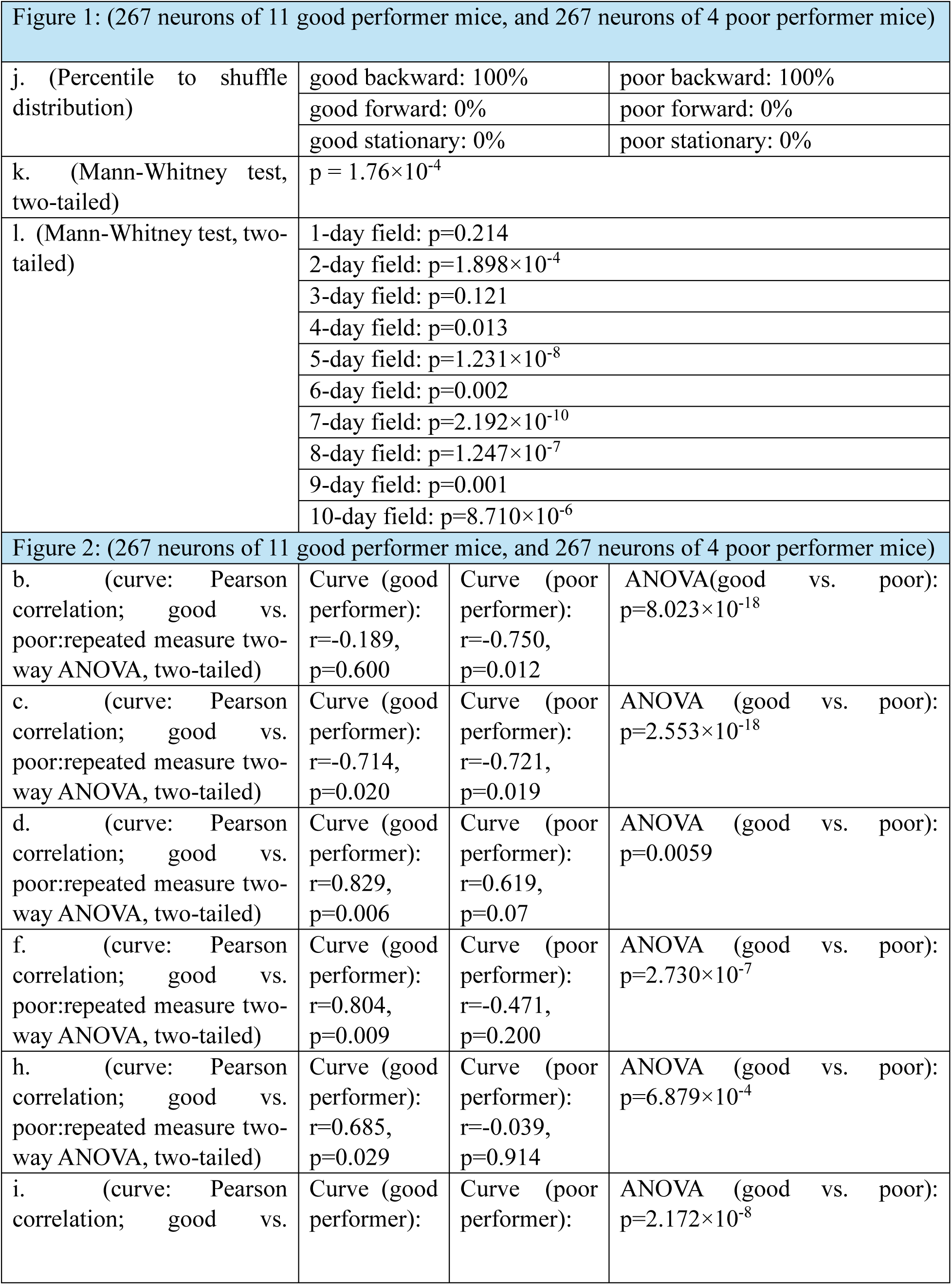
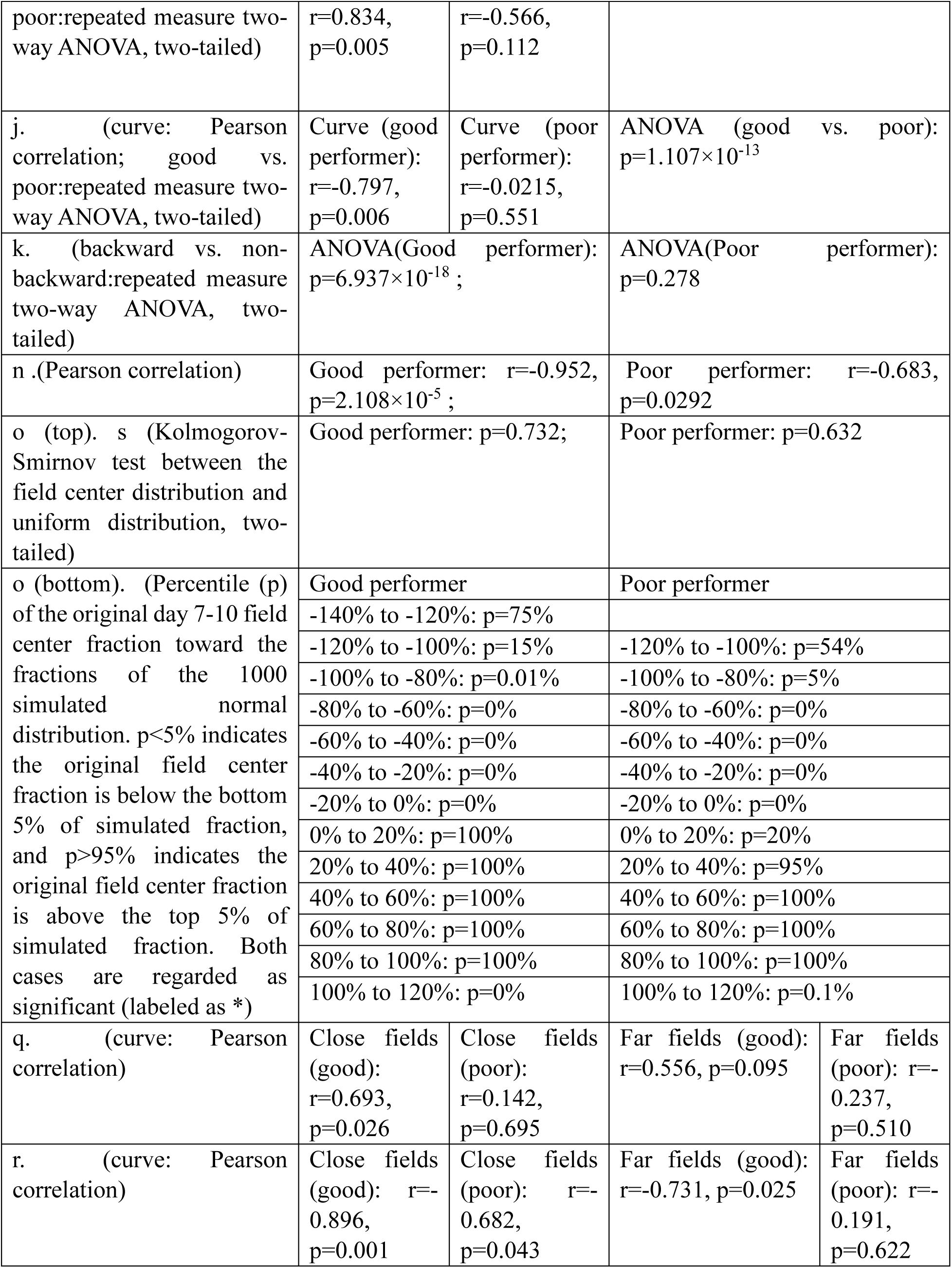
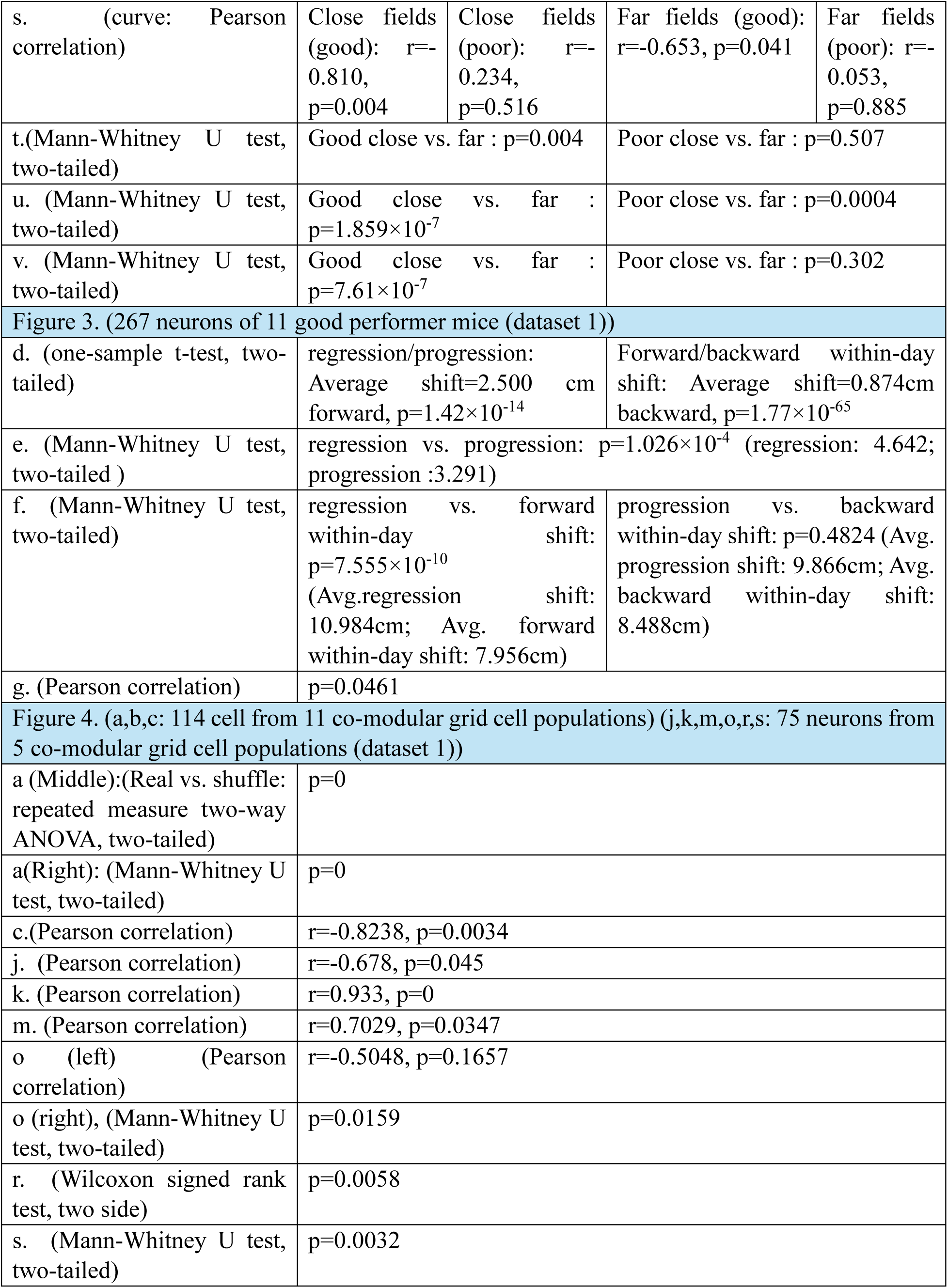
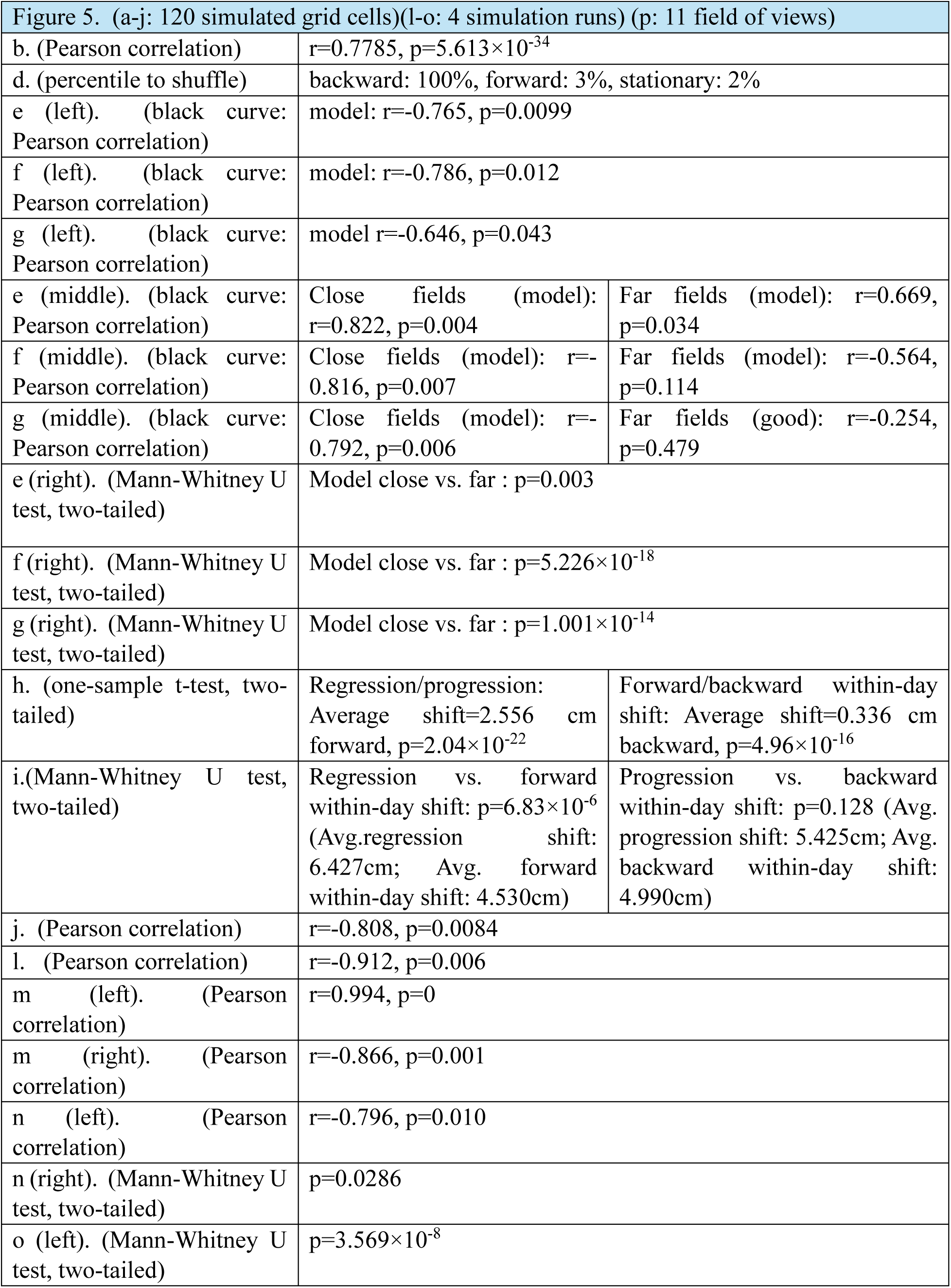
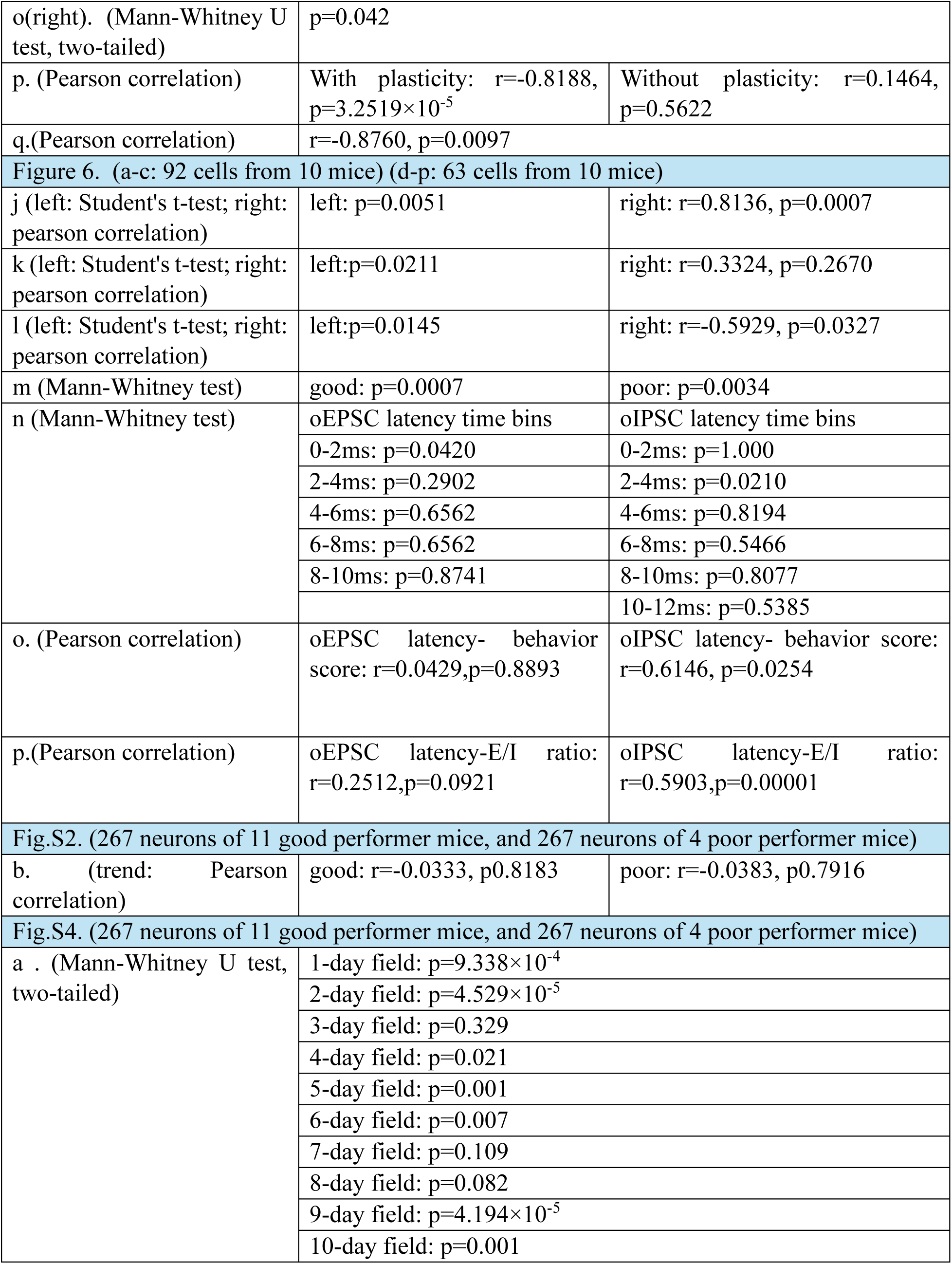
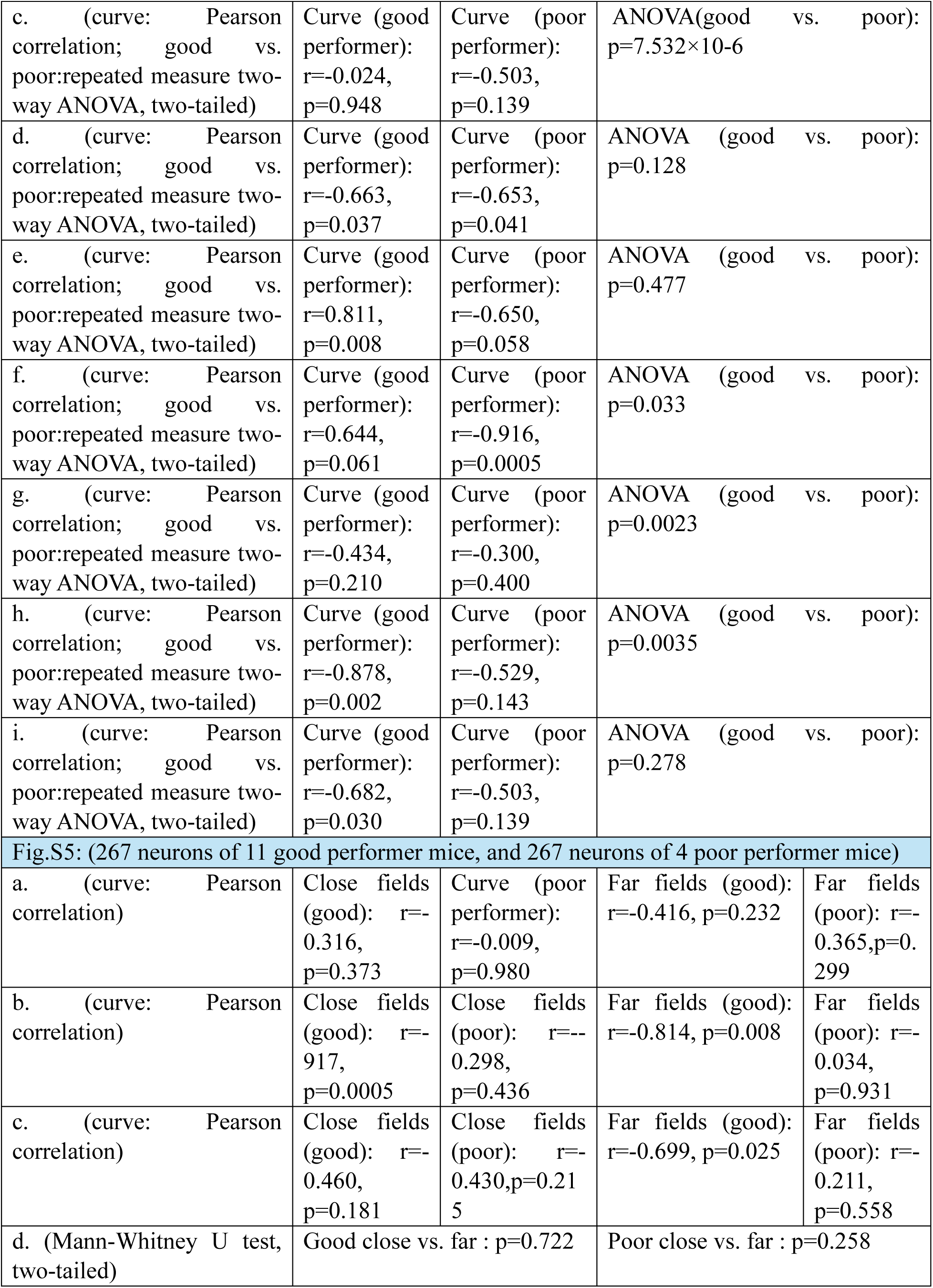
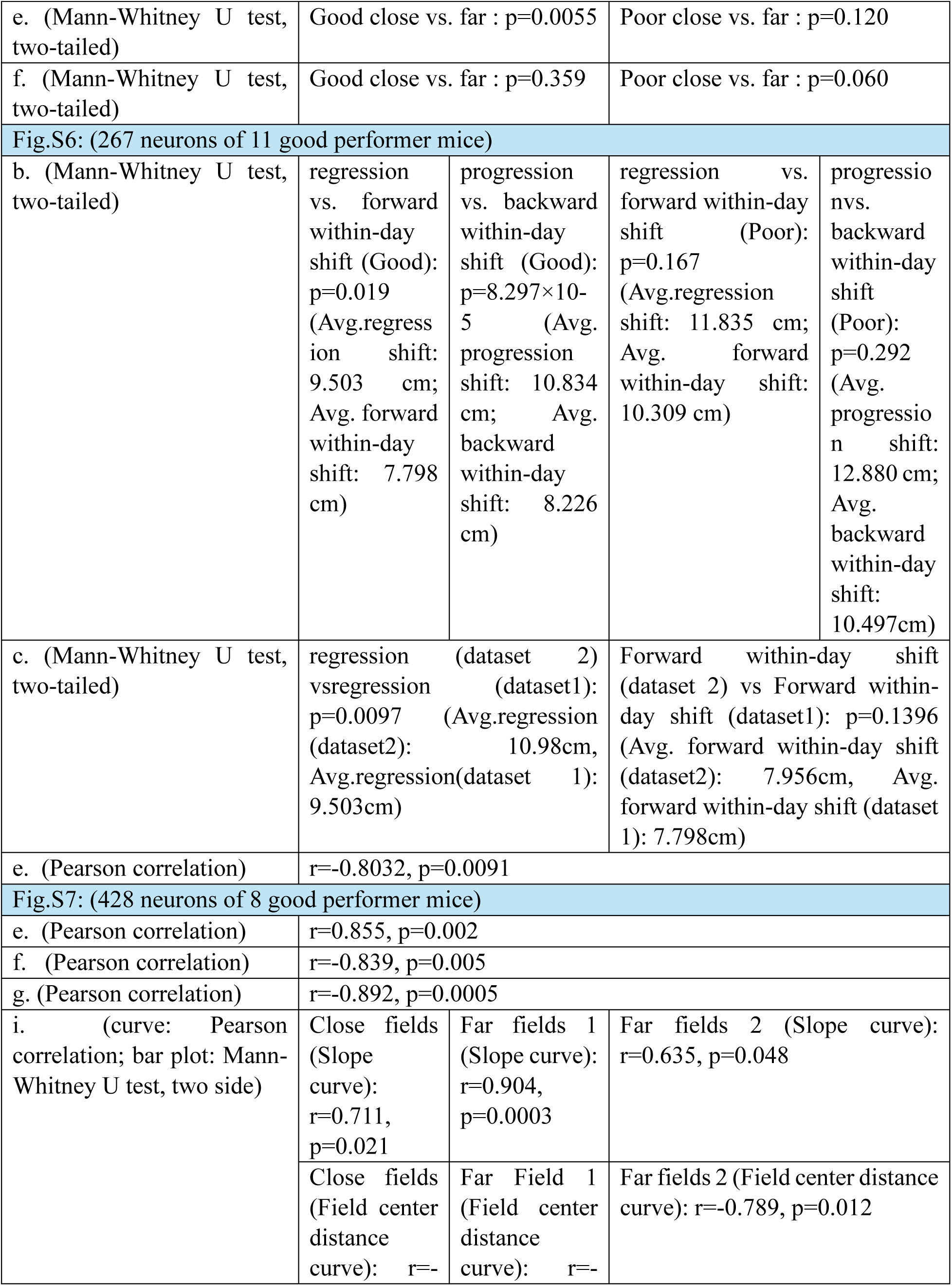
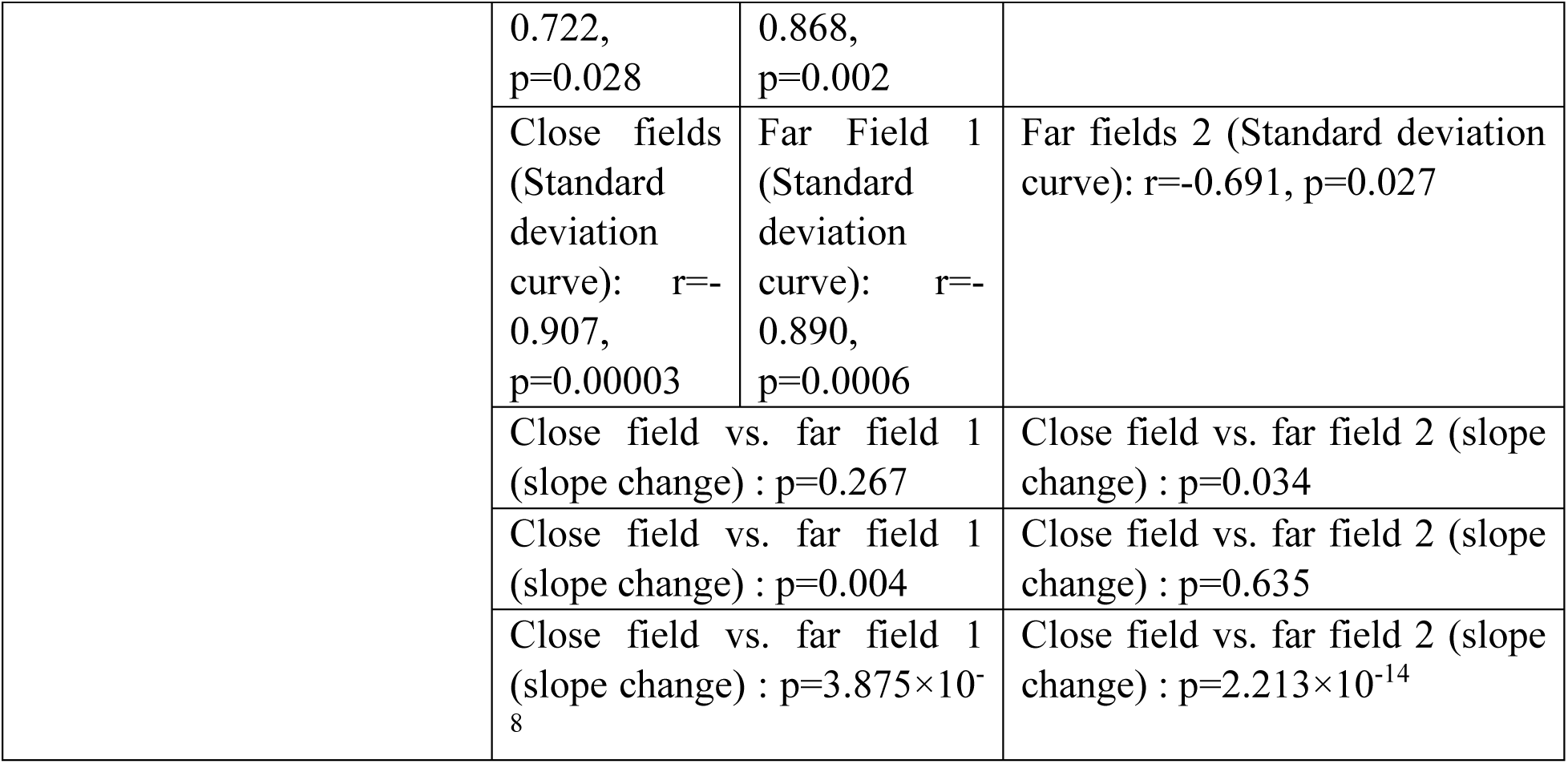
Statistics of analysis in each figure.

| Figure 1: (267 neurons of 11 good performer mice, and 267 neurons of 4 poor performer mice) |  |  |  |
| --- | --- | --- | --- |
| j. (Percentile to shuffle distribution) | good backward: 100% |  | poor backward: 100% |
|  | good forward: 0% |  | poor forward: 0% |
|  | good stationary: 0% |  | poor stationary: 0% |
| k. (Mann-Whitney test, two-tailed) | p = 1.76×10<sup>-4</sup> |  |  |
| l. (Mann-Whitney test, two-tailed) | 1-day field: p=0.214 |  |  |
|  | 2-day field: p=1.898×10<sup>-4</sup> |  |  |
|  | 3-day field: p=0.121 |  |  |
|  | 4-day field: p=0.013 |  |  |
|  | 5-day field: p=1.231×10<sup>-8</sup> |  |  |
|  | 6-day field: p=0.002 |  |  |
|  | 7-day field: p=2.192×10<sup>-10</sup> |  |  |
|  | 8-day field: p=1.247×10<sup>-7</sup> |  |  |
|  | 9-day field: p=0.001 |  |  |
|  | 10-day field: p=8.710×10<sup>-6</sup> |  |  |
| Figure 2: (267 neurons of 11 good performer mice, and 267 neurons of 4 poor performer mice) |  |  |  |
| b. (curve: Pearson correlation; good vs. poor:repeated measure two-way ANOVA, two-tailed) | Curve (good performer):<br>r=-0.189,<br>p=0.600 | Curve (poor performer):<br>r=-0.750,<br>p=0.012 | ANOVA(good vs. poor):<br>p=8.023×10<sup>-18</sup> |
| c. (curve: Pearson correlation; good vs. poor:repeated measure two-way ANOVA, two-tailed) | Curve (good performer):<br>r=-0.714,<br>p=0.020 | Curve (poor performer):<br>r=-0.721,<br>p=0.019 | ANOVA (good vs. poor):<br>p=2.553×10<sup>-18</sup> |
| d. (curve: Pearson correlation; good vs. poor:repeated measure two-way ANOVA, two-tailed) | Curve (good performer):<br>r=0.829,<br>p=0.006 | Curve (poor performer):<br>r=0.619,<br>p=0.07 | ANOVA (good vs. poor):<br>p=0.0059 |
| f. (curve: Pearson correlation; good vs. poor:repeated measure two-way ANOVA, two-tailed) | Curve (good performer):<br>r=0.804,<br>p=0.009 | Curve (poor performer):<br>r=-0.471,<br>p=0.200 | ANOVA (good vs. poor):<br>p=2.730×10<sup>-7</sup> |
| h. (curve: Pearson correlation; good vs. poor:repeated measure two-way ANOVA, two-tailed) | Curve (good performer):<br>r=0.685,<br>p=0.029 | Curve (poor performer):<br>r=-0.039,<br>p=0.914 | ANOVA (good vs. poor):<br>p=6.879×10<sup>-4</sup> |
| i. (curve: Pearson correlation; good vs. poor:repeated measure two-way ANOVA, two-tailed) | Curve (good performer): | Curve (poor performer): | ANOVA (good vs. poor):<br>p=2.172×10<sup>-8</sup> |

|  |  |  |  |  |
| --- | --- | --- | --- | --- |
| poor:repeated measure two-way ANOVA, two-tailed) | r=0.834,<br>p=0.005 | r=-0.566,<br>p=0.112 |  |  |
| j. (curve: Pearson correlation; good vs. poor:repeated measure two-way ANOVA, two-tailed) | Curve (good performer):<br>r=-0.797,<br>p=0.006 | Curve (poor performer):<br>r=-0.0215,<br>p=0.551 | ANOVA (good vs. poor):<br>p=1.107×10<sup>-13</sup> |  |
| k. (backward vs. non-backward:repeated measure two-way ANOVA, two-tailed) | ANOVA(Good performer):<br>p=6.937×10<sup>-18</sup> ; |  | ANOVA(Poor performer):<br>p=0.278 |  |
| n.(Pearson correlation) | Good performer: r=-0.952,<br>p=2.108×10<sup>-5</sup> ; |  | Poor performer: r=-0.683,<br>p=0.0292 |  |
| o (top). s (Kolmogorov-Smirnov test between the field center distribution and uniform distribution, two-tailed) | Good performer: p=0.732; |  | Poor performer: p=0.632 |  |
| o (bottom). (Percentile (p) of the original day 7-10 field center fraction toward the fractions of the 1000 simulated normal distribution. p<5% indicates the original field center fraction is below the bottom 5% of simulated fraction, and p>95% indicates the original field center fraction is above the top 5% of simulated fraction. Both cases are regarded as significant (labeled as *)) | Good performer |  | Poor performer |  |
|  | -140% to -120%: p=75% |  |  |  |
|  | -120% to -100%: p=15% |  | -120% to -100%: p=54% |  |
|  | -100% to -80%: p=0.01% |  | -100% to -80%: p=5% |  |
|  | -80% to -60%: p=0% |  | -80% to -60%: p=0% |  |
|  | -60% to -40%: p=0% |  | -60% to -40%: p=0% |  |
|  | -40% to -20%: p=0% |  | -40% to -20%: p=0% |  |
|  | -20% to 0%: p=0% |  | -20% to 0%: p=0% |  |
|  | 0% to 20%: p=100% |  | 0% to 20%: p=20% |  |
|  | 20% to 40%: p=100% |  | 20% to 40%: p=95% |  |
|  | 40% to 60%: p=100% |  | 40% to 60%: p=100% |  |
|  | 60% to 80%: p=100% |  | 60% to 80%: p=100% |  |
|  | 80% to 100%: p=100% |  | 80% to 100%: p=100% |  |
|  | 100% to 120%: p=0% |  | 100% to 120%: p=0.1% |  |
| q. (curve: Pearson correlation) | Close fields (good):<br>r=0.693,<br>p=0.026 | Close fields (poor):<br>r=0.142,<br>p=0.695 | Far fields (good):<br>r=0.556, p=0.095 | Far fields (poor): r=-0.237,<br>p=0.510 |
| r. (curve: Pearson correlation) | Close fields (good): r=-0.896,<br>p=0.001 | Close fields (poor): r=-0.682,<br>p=0.043 | Far fields (good):<br>r=-0.731, p=0.025 | Far fields (poor): r=-0.191,<br>p=0.622 |
| s. (curve: Pearson correlation) | Close fields (good): $r=-0.810$, $p=0.004$ | Close fields (poor): $r=-0.234$, $p=0.516$ | Far fields (good): $r=-0.653$, $p=0.041$ | Far fields (poor): $r=-0.053$, $p=0.885$ |
| t.(Mann-Whitney U test, two-tailed) | Good close vs. far : $p=0.004$ | | Poor close vs. far : $p=0.507$ | |
| u. (Mann-Whitney U test, two-tailed) | Good close vs. far : $p=1.859\times10^{-7}$ | | Poor close vs. far : $p=0.0004$ | |
| v. (Mann-Whitney U test, two-tailed) | Good close vs. far : $p=7.61\times10^{-7}$ | | Poor close vs. far : $p=0.302$ | |
| Figure 3. (267 neurons of 11 good performer mice (dataset 1)) |  |  |  |  |
| d. (one-sample t-test, two-tailed) | regression/progression: Average shift=2.500 cm forward, $p=1.42\times10^{-14}$ | | Forward/backward within-day shift: Average shift=0.874cm backward, $p=1.77\times10^{-65}$ | |
| e. (Mann-Whitney U test, two-tailed ) | regression vs. progression: $p=1.026\times10^{-4}$ (regression: 4.642; progression :3.291) | | | |
| f. (Mann-Whitney U test, two-tailed) | regression vs. forward within-day shift: $p=7.555\times10^{-10}$ (Avg.regression shift: 10.984cm; Avg. forward within-day shift: 7.956cm) | | progression vs. backward within-day shift: $p=0.4824$ (Avg. progression shift: 9.866cm; Avg. backward within-day shift: 8.488cm) | |
| g. (Pearson correlation) | $p=0.0461$ | | | |
| Figure 4. (a,b,c: 114 cell from 11 co-modular grid cell populations) (j,k,m,o,r,s: 75 neurons from 5 co-modular grid cell populations (dataset 1)) |  |  |  |  |
| a (Middle):(Real vs. shuffle: repeated measure two-way ANOVA, two-tailed) | $p=0$ | | | |
| a(Right): (Mann-Whitney U test, two-tailed) | $p=0$ | | | |
| c.(Pearson correlation) | $r=-0.8238$, $p=0.0034$ | | | |
| j. (Pearson correlation) | $r=-0.678$, $p=0.045$ | | | |
| k. (Pearson correlation) | $r=0.933$, $p=0$ | | | |
| m. (Pearson correlation) | $r=0.7029$, $p=0.0347$ | | | |
| o (left) (Pearson correlation) | $r=-0.5048$, $p=0.1657$ | | | |
| o (right), (Mann-Whitney U test, two-tailed) | $p=0.0159$ | | | |
| r. (Wilcoxon signed rank test, two side) | $p=0.0058$ | | | |
| s. (Mann-Whitney U test, two-tailed) | $p=0.0032$ | | | |

| Figure 5. (a-j: 120 simulated grid cells)(l-o: 4 simulation runs) (p: 11 field of views) |  |  |
| --- | --- | --- |
| b. (Pearson correlation) | r=0.7785, p=5.613×10<sup>-34</sup> |  |
| d. (percentile to shuffle) | backward: 100%, forward: 3%, stationary: 2% |  |
| e (left). (black curve: Pearson correlation) | model: r=-0.765, p=0.0099 |  |
| f (left). (black curve: Pearson correlation) | model: r=-0.786, p=0.012 |  |
| g (left). (black curve: Pearson correlation) | model r=-0.646, p=0.043 |  |
| e (middle). (black curve: Pearson correlation) | Close fields (model): r=0.822, p=0.004 | Far fields (model): r=0.669, p=0.034 |
| f (middle). (black curve: Pearson correlation) | Close fields (model): r=-0.816, p=0.007 | Far fields (model): r=-0.564, p=0.114 |
| g (middle). (black curve: Pearson correlation) | Close fields (model): r=-0.792, p=0.006 | Far fields (good): r=-0.254, p=0.479 |
| e (right). (Mann-Whitney U test, two-tailed) | Model close vs. far : p=0.003 |  |
| f (right). (Mann-Whitney U test, two-tailed) | Model close vs. far : p=5.226×10<sup>-18</sup> |  |
| g (right). (Mann-Whitney U test, two-tailed) | Model close vs. far : p=1.001×10<sup>-14</sup> |  |
| h. (one-sample t-test, two-tailed) | Regression/progression: Average shift=2.556 cm forward, p=2.04×10<sup>-22</sup> | Forward/backward within-day shift: Average shift=0.336 cm backward, p=4.96×10<sup>-16</sup> |
| i.(Mann-Whitney U test, two-tailed) | Regression vs. forward within-day shift: p=6.83×10<sup>-6</sup> (Avg.regression shift: 6.427cm; Avg. forward within-day shift: 4.530cm) | Progression vs. backward within-day shift: p=0.128 (Avg. progression shift: 5.425cm; Avg. backward within-day shift: 4.990cm) |
| j. (Pearson correlation) | r=-0.808, p=0.0084 |  |
| l. (Pearson correlation) | r=-0.912, p=0.006 |  |
| m (left). (Pearson correlation) | r=0.994, p=0 |  |
| m (right). (Pearson correlation) | r=-0.866, p=0.001 |  |
| n (left). (Pearson correlation) | r=-0.796, p=0.010 |  |
| n (right). (Mann-Whitney U test, two-tailed) | p=0.0286 |  |
| o (left). (Mann-Whitney U test, two-tailed) | p=3.569×10<sup>-8</sup> |  |
| o(right). (Mann-Whitney U test, two-tailed) | p=0.042 |  |
| p. (Pearson correlation) | With plasticity: r=-0.8188, p=3.2519×10<sup>-5</sup> | Without plasticity: r=0.1464, p=0.5622 |
| q.(Pearson correlation) | r=-0.8760, p=0.0097 |  |
| Figure 6. (a-c: 92 cells from 10 mice) (d-p: 63 cells from 10 mice) |  |  |
| j (left: Student's t-test; right: pearson correlation) | left: p=0.0051 | right: r=0.8136, p=0.0007 |
| k (left: Student's t-test; right: pearson correlation) | left:p=0.0211 | right: r=0.3324, p=0.2670 |
| l (left: Student's t-test; right: pearson correlation) | left:p=0.0145 | right: r=-0.5929, p=0.0327 |
| m (Mann-Whitney test) | good: p=0.0007 | poor: p=0.0034 |
| n (Mann-Whitney test) | oEPSC latency time bins | oIPSC latency time bins |
|  | 0-2ms: p=0.0420 | 0-2ms: p=1.000 |
|  | 2-4ms: p=0.2902 | 2-4ms: p=0.0210 |
|  | 4-6ms: p=0.6562 | 4-6ms: p=0.8194 |
|  | 6-8ms: p=0.6562 | 6-8ms: p=0.5466 |
|  | 8-10ms: p=0.8741 | 8-10ms: p=0.8077 |
|  |  | 10-12ms: p=0.5385 |
| o. (Pearson correlation) | oEPSC latency- behavior score: r=0.0429,p=0.8893 | oIPSC latency- behavior score: r=0.6146, p=0.0254 |
| p.(Pearson correlation) | oEPSC latency-E/I ratio: r=0.2512,p=0.0921 | oIPSC latency-E/I ratio: r=0.5903,p=0.00001 |
| Fig.S2. (267 neurons of 11 good performer mice, and 267 neurons of 4 poor performer mice) |  |  |
| b. (trend: Pearson correlation) | good: r=-0.0333, p0.8183 | poor: r=-0.0383, p0.7916 |
| Fig.S4. (267 neurons of 11 good performer mice, and 267 neurons of 4 poor performer mice) |  |  |
| a . (Mann-Whitney U test, two-tailed) | 1-day field: p=9.338×10<sup>-4</sup> |  |
|  | 2-day field: p=4.529×10<sup>-5</sup> |  |
|  | 3-day field: p=0.329 |  |
|  | 4-day field: p=0.021 |  |
|  | 5-day field: p=0.001 |  |
|  | 6-day field: p=0.007 |  |
|  | 7-day field: p=0.109 |  |
|  | 8-day field: p=0.082 |  |
|  | 9-day field: p=4.194×10<sup>-5</sup> |  |
|  | 10-day field: p=0.001 |  |

|  |  |  |  |  |
| --- | --- | --- | --- | --- |
| c. (curve: Pearson correlation; good vs. poor:repeated measure two-way ANOVA, two-tailed) | Curve (good performer):<br>r=-0.024,<br>p=0.948 | Curve (poor performer):<br>r=-0.503,<br>p=0.139 | ANOVA(good vs. poor):<br>p=7.532×10-6 |  |
| d. (curve: Pearson correlation; good vs. poor:repeated measure two-way ANOVA, two-tailed) | Curve (good performer):<br>r=-0.663,<br>p=0.037 | Curve (poor performer):<br>r=-0.653,<br>p=0.041 | ANOVA (good vs. poor):<br>p=0.128 |  |
| e. (curve: Pearson correlation; good vs. poor:repeated measure two-way ANOVA, two-tailed) | Curve (good performer):<br>r=0.811,<br>p=0.008 | Curve (poor performer):<br>r=-0.650,<br>p=0.058 | ANOVA (good vs. poor):<br>p=0.477 |  |
| f. (curve: Pearson correlation; good vs. poor:repeated measure two-way ANOVA, two-tailed) | Curve (good performer):<br>r=0.644,<br>p=0.061 | Curve (poor performer):<br>r=-0.916,<br>p=0.0005 | ANOVA (good vs. poor):<br>p=0.033 |  |
| g. (curve: Pearson correlation; good vs. poor:repeated measure two-way ANOVA, two-tailed) | Curve (good performer):<br>r=-0.434,<br>p=0.210 | Curve (poor performer):<br>r=-0.300,<br>p=0.400 | ANOVA (good vs. poor):<br>p=0.0023 |  |
| h. (curve: Pearson correlation; good vs. poor:repeated measure two-way ANOVA, two-tailed) | Curve (good performer):<br>r=-0.878,<br>p=0.002 | Curve (poor performer):<br>r=-0.529,<br>p=0.143 | ANOVA (good vs. poor):<br>p=0.0035 |  |
| i. (curve: Pearson correlation; good vs. poor:repeated measure two-way ANOVA, two-tailed) | Curve (good performer):<br>r=-0.682,<br>p=0.030 | Curve (poor performer):<br>r=-0.503,<br>p=0.139 | ANOVA (good vs. poor):<br>p=0.278 |  |
| Fig.S5: (267 neurons of 11 good performer mice, and 267 neurons of 4 poor performer mice) |  |  |  |  |
| a. (curve: Pearson correlation) | Close fields (good): r=-0.316,<br>p=0.373 | Curve (poor performer):<br>r=-0.009,<br>p=0.980 | Far fields (good):<br>r=-0.416, p=0.232 | Far fields (poor): r=-0.365,p=0.299 |
| b. (curve: Pearson correlation) | Close fields (good): r=-0.917,<br>p=0.0005 | Close fields (poor): r=-0.298,<br>p=0.436 | Far fields (good):<br>r=-0.814, p=0.008 | Far fields (poor): r=-0.034,<br>p=0.931 |
| c. (curve: Pearson correlation) | Close fields (good): r=-0.460,<br>p=0.181 | Close fields (poor): r=-0.430,p=0.215 | Far fields (good):<br>r=-0.699, p=0.025 | Far fields (poor): r=-0.211,<br>p=0.558 |
| d. (Mann-Whitney U test, two-tailed) | Good close vs. far : p=0.722 |  | Poor close vs. far : p=0.258 |  |
| e. (Mann-Whitney U test, two-tailed) | Good close vs. far : p=0.0055 |  | Poor close vs. far : p=0.120 |  |
| f. (Mann-Whitney U test, two-tailed) | Good close vs. far : p=0.359 |  | Poor close vs. far : p=0.060 |  |
| Fig.S6: (267 neurons of 11 good performer mice) |  |  |  |  |
| b. (Mann-Whitney U test, two-tailed) | regression vs. forward within-day shift (Good): p=0.019 (Avg.regression shift: 9.503 cm; Avg. forward within-day shift: 7.798 cm) | progression vs. backward within-day shift (Good): p=8.297×10-5 (Avg. progression shift: 10.834 cm; Avg. backward within-day shift: 8.226 cm) | regression vs. forward within-day shift (Poor): p=0.167 (Avg.regression shift: 11.835 cm; Avg. forward within-day shift: 10.309 cm) | progression vs. backward within-day shift (Poor): p=0.292 (Avg. progression shift: 12.880 cm; Avg. backward within-day shift: 10.497cm) |
| c. (Mann-Whitney U test, two-tailed) | regression (dataset 2) vs regression (dataset1): p=0.0097 (Avg.regression (dataset2): 10.98cm, Avg.regression(dataset 1): 9.503cm) |  | Forward within-day shift (dataset 2) vs Forward within-day shift (dataset1): p=0.1396 (Avg. forward within-day shift (dataset2): 7.956cm, Avg. forward within-day shift (dataset 1): 7.798cm) |  |
| e. (Pearson correlation) | r=-0.8032, p=0.0091 |  |  |  |
| Fig.S7: (428 neurons of 8 good performer mice) |  |  |  |  |
| e. (Pearson correlation) | r=0.855, p=0.002 |  |  |  |
| f. (Pearson correlation) | r=-0.839, p=0.005 |  |  |  |
| g. (Pearson correlation) | r=-0.892, p=0.0005 |  |  |  |
| i. (curve: Pearson correlation; bar plot: Mann-Whitney U test, two side) | Close fields (Slope curve): r=0.711, p=0.021 | Far fields 1 (Slope curve): r=0.904, p=0.0003 | Far fields 2 (Slope curve): r=0.635, p=0.048 |  |
|  | Close fields (Field center distance curve): r=- | Far Field 1 (Field center distance curve): r=- | Far fields 2 (Field center distance curve): r=-0.789, p=0.012 |  |
|  | 0.722,<br>p=0.028 | 0.868,<br>p=0.002 |  |
|  | Close fields<br>(Standard<br>deviation<br>curve): r=-<br>0.907,<br>p=0.00003 | Far Field 1<br>(Standard<br>deviation<br>curve): r=-<br>0.890,<br>p=0.0006 | Far fields 2 (Standard deviation<br>curve): r=-0.691, p=0.027 |
|  | Close field vs. far field 1<br>(slope change) : p=0.267 |  | Close field vs. far field 2 (slope<br>change) : p=0.034 |
|  | Close field vs. far field 1<br>(slope change) : p=0.004 |  | Close field vs. far field 2 (slope<br>change) : p=0.635 |
|  | Close field vs. far field 1<br>(slope change) : p=3.875×10<sup>-8</sup> |  | Close field vs. far field 2 (slope<br>change) : p=2.213×10<sup>-14</sup> |

## References

1. E. C. Tolman, Cognitive maps in rats and men. Psychol Rev 55, 189–208 (1948).

2. M. Hernandez-Frausto, C. Vivar, Entorhinal cortex-hippocampal circuit connectivity in health and disease. Front Hum Neurosci 18, 1448791 (2024).

3. J. O’Keefe, J. Krupic, Do hippocampal pyramidal cells respond to nonspatial stimuli? Physiol Rev 101, 1427–1456 (2021).

4. L. L. Dong, I. R. Fiete, Grid Cells in Cognition: Mechanisms and Function. Annual Review of Neuroscience 47, 345–368 (2024).

5. J. O’Keefe, J. Dostrovsky, The hippocampus as a spatial map. Preliminary evidence from unit activity in the freely-moving rat. Brain Res 34, 171–175 (1971).

6. T. Hafting, M. Fyhn, S. Molden, M.-B. Moser, E. I. Moser, Microstructure of a spatial map in the entorhinal cortex. Nature 436, 801–806 (2005).

7. D. Derdikman et al., Fragmentation of grid cell maps in a multicompartment environment. Nat Neurosci 12, 1325–1332 (2009).

8. A. J. Hill, First occurrence of hippocampal spatial firing in a new environment. Exp Neurol 62, 282–297 (1978).

9. M. A. Wilson, B. L. McNaughton, Dynamics of the hippocampal ensemble code for space. Science 261, 1055–1058 (1993).

10. L. M. Frank, G. B. Stanley, E. N. Brown, Hippocampal plasticity across multiple days of exposure to novel environments. J Neurosci 24, 7681–7689 (2004).

11. J. H. Wen, B. Sorscher, E. A. Aery Jones, S. Ganguli, L. M. Giocomo, One-shot entorhinal maps enable flexible navigation in novel environments. Nature 635, 943–950 (2024).

12. K. C. Bittner, A. D. Milstein, C. Grienberger, S. Romani, J. C. Magee, Behavioral time scale synaptic plasticity underlies CA1 place fields. Science 357, 1033–1036 (2017).

13. Y. Burak, I. R. Fiete, Accurate Path Integration in Continuous Attractor Network Models of Grid Cells. PLOS Computational Biology 5, e1000291 (2009).

14. M. C. Fuhs, D. S. Touretzky, A spin glass model of path integration in rat medial entorhinal cortex. J Neurosci 26, 4266–4276 (2006).

15. A. Guanella, D. Kiper, P. Verschure, A model of grid cells based on a twisted torus topology. Int J Neural Syst 17, 231–240 (2007).

16. S. G. Trettel, J. B. Trimper, E. Hwaun, I. R. Fiete, L. L. Colgin, Grid cell co-activity patterns during sleep reflect spatial overlap of grid fields during active behaviors. Nat Neurosci 22, 609–617 (2019).

17. R. J. Gardner, L. Lu, T. Wernle, M. B. Moser, E. I. Moser, Correlation structure of grid cells is preserved during sleep. Nat Neurosci 22, 598–608 (2019).

18. R. J. Gardner et al., Toroidal topology of population activity in grid cells. Nature 602, 123–128 (2022).

19. C. N. Boccara, M. Nardin, F. Stella, J. O’Neill, J. Csicsvari, The entorhinal cognitive map is attracted to goals. Science 363, 1443–1447 (2019).

20. W. N. Butler, K. Hardcastle, L. M. Giocomo, Remembered reward locations restructure entorhinal spatial maps. Science 363, 1447–1452 (2019).

21. Y. Ziv et al., Long-term dynamics of CA1 hippocampal place codes. Nat Neurosci 16, 264–266 (2013).

22. J. D. Monaco, G. Rao, E. D. Roth, J. J. Knierim, Attentive scanning behavior drives one-trial potentiation of hippocampal place fields. Nat Neurosci 17, 725–731 (2014).

23. S. P. Vaidya, G. Li, R. A. Chitwood, Y. Li, J. C. Magee, Formation of an expanding memory representation in the hippocampus. Nat Neurosci 28, 1510–1518 (2025).

24. C. Dong, A. D. Madar, M. E. J. Sheffield, Distinct place cell dynamics in CA1 and CA3 encode experience in new environments. Nature Communications 12, 2977 (2021).

25. I. Kanitscheider, Fiete, I., Making our way through the world: Towards a functional understanding of the brain’s spatial circuits. Current Opinion in Systems Biology 3, 186–194 (2017).

26. J. Widloski, Fiete, I., in Space,Time and Memory in the Hippocampal Formation., D. Derdikman, Knierim, J., Ed. (Springer, Vienna, 2014).

27. J. A. Mayer, Filliat, D., Map-based navigation in mobile robots: II. A review of map-learning and path-planning strategies. Cogn Syst Res 4, 283–317 (2003).

28. H. Durrant-Whyte, Bailey, T., Simultaneous localization and mapping (SLAM): part 1 the essential algorithms. Robot Autom Mag 13, (2006).

29. C. Barry, L. L. Ginzberg, J. O’Keefe, N. Burgess, Grid cell firing patterns signal environmental novelty by expansion. Proc Natl Acad Sci U S A 109, 17687–17692 (2012).

30. J. Krupic, M. Bauza, S. Burton, C. Barry, J. O’Keefe, Grid cell symmetry is shaped by environmental geometry. Nature 518, 232–235 (2015).

31. B. E. Gutierrez-Guzman, J. J. Hernandez-Perez, H. Dannenberg, Tiling of large-scaled environments by grid cells requires experience. bioRxiv, (2025).

32. M. R. Mehta, C. A. Barnes, B. L. McNaughton, Experience-dependent, asymmetric expansion of hippocampal place fields. Proc Natl Acad Sci U S A 94, 8918–8921 (1997).

33. T. Geiller, M. Fattahi, J. S. Choi, S. Royer, Place cells are more strongly tied to landmarks in deep than in superficial CA1. Nat Commun 8, 14531 (2017).

34. M. R. Mehta, M. C. Quirk, M. A. Wilson, Experience-dependent asymmetric shape of hippocampal receptive fields. Neuron 25, 707–715 (2000).

35. I. Lee, G. Rao, J. J. Knierim, A double dissociation between hippocampal subfields: differential time course of CA3 and CA1 place cells for processing changed environments. Neuron 42, 803–815 (2004).

36. A. D. Ekstrom, J. Meltzer, B. L. McNaughton, C. A. Barnes, NMDA receptor antagonism blocks experience-dependent expansion of hippocampal “place fields”. Neuron 31, 631–638 (2001).

37. E. D. Roth, X. Yu, G. Rao, J. J. Knierim, Functional differences in the backward shifts of CA1 and CA3 place fields in novel and familiar environments. PLoS One 7, e36035 (2012).

38. T. J. Malone et al., A consistent map in the medial entorhinal cortex supports spatial memory. Nature Communications 15, 1457 (2024).

39. H. Dana et al., Thy1-GCaMP6 transgenic mice for neuronal population imaging in vivo. PLoS One 9, e108697 (2014).

40. T. W. Chen et al., Ultrasensitive fluorescent proteins for imaging neuronal activity. Nature 499, 295–300 (2013).

41. Y. Gu et al., A Map-like Micro-Organization of Grid Cells in the Medial Entorhinal Cortex. Cell 175, 736–750.e730 (2018).

42. A. A. Kinkhabwala, Y. Gu, D. Aronov, D. W. Tank, Visual cue-related activity of cells in the medial entorhinal cortex during navigation in virtual reality. eLife 9, e43140 (2020).

43. A. Goto, Y. Hayashi, Offline neuronal activity and synaptic plasticity during sleep and memory consolidation. Neurosci Res 189, 29–36 (2023).

44. M. S. Halvagal, F. Zenke, The combination of Hebbian and predictive plasticity learns invariant object representations in deep sensory networks. Nat Neurosci 26, 1906–1915 (2023).

45. J. L. McGaugh, Memory--a Century of Consolidation. Science 287, 248–251 (2000).

46. O. Tchernichovski, D. Margoliash, in Animal Models of Speech and Language Disorders, S. A. Helekar, Ed. (Springer New York, New York, NY, 2013), pp. 43–60.

47. B. P. Ölveczky, A. S. Andalman, M. S. Fee, Vocal Experimentation in the Juvenile Songbird Requires a Basal Ganglia Circuit. PLOS Biology 3, e153 (2005).

48. B. L. McNaughton, F. P. Battaglia, O. Jensen, E. I. Moser, M. B. Moser, Path integration and the neural basis of the ‘cognitive map’. Nat Rev Neurosci 7, 663–678 (2006).

49. M. Fyhn, T. Hafting, A. Treves, M. B. Moser, E. I. Moser, Hippocampal remapping and grid realignment in entorhinal cortex. Nature 446, 190–194 (2007).

50. T. Wernle et al., Integration of grid maps in merged environments. Nature Neuroscience 21, 92–101 (2018).

51. H. Stensola et al., The entorhinal grid map is discretized. Nature 492, 72–78 (2012).

52. K. Yoon et al., Specific evidence of low-dimensional continuous attractor dynamics in grid cells. Nature Neuroscience 16, 1077–1084 (2013).

53. K. Yoon, S. Lewallen, Amina A. Kinkhabwala, David W. Tank, Ila R. Fiete, Grid Cell Responses in 1D Environments Assessed as Slices through a 2D Lattice. Neuron 89, 1086–1099 (2016).

54. S. Chandra, S. Sharma, R. Chaudhuri, I. Fiete, Episodic and associative memory from spatial scaffolds in the hippocampus. Nature, (2025).

55. R. M. Yoder, B. J. Clark, J. S. Taube, Origins of landmark encoding in the brain. Trends in Neurosciences 34, 561–571 (2011).

56. Ø. A. Høydal, E. R. Skytøen, S. O. Andersson, M.-B. Moser, E. I. Moser, Object-vector coding in the medial entorhinal cortex. Nature 568, 400–404 (2019).

57. G. Casali, S. Shipley, C. Dowell, R. Hayman, C. Barry, Entorhinal Neurons Exhibit Cue Locking in Rodent VR. Frontiers in Cellular Neuroscience Volume 12 **-** 2018, (2019).

58. S. S. Deshmukh, J. J. Knierim, Influence of local objects on hippocampal representations: Landmark vectors and memory. Hippocampus 23, 253–267 (2013).

59. K. M. Scaplen, A. A. Gulati, V. L. Heimer-McGinn, R. D. Burwell, Objects and landmarks: Hippocampal place cells respond differently to manipulations of visual cues depending on size, perspective, and experience. Hippocampus 24, 1287–1299 (2014).

60. J. Bischofberger, D. Engel, L. Li, J. R. P. Geiger, P. Jonas, Patch-clamp recording from mossy fiber terminals in hippocampal slices. Nature Protocols 1, 2075–2081 (2006).

61. A. Rafiq, R. J. DeLorenzo, D. A. Coulter, Generation and propagation of epileptiform discharges in a combined entorhinal cortex/hippocampal slice. Journal of Neurophysiology 70, 1962–1974 (1993).

62. A. C. Au - Whitebirch, JoVE, e61704 (2020).

63. A. Rozov et al., Processing of Hippocampal Network Activity in the Receiver Network of the Medial Entorhinal Cortex Layer V. J Neurosci 40, 8413–8425 (2020).

64. G. Surmeli et al., Molecularly Defined Circuitry Reveals Input-Output Segregation in Deep Layers of the Medial Entorhinal Cortex. Neuron 88, 1040–1053 (2015).

65. S. Ohara et al., Intrinsic Projections of Layer Vb Neurons to Layers Va, III, and II in the Lateral and Medial Entorhinal Cortex of the Rat. Cell Rep 24, 107–116 (2018).

66. S. Ohara et al., Hippocampal-medial entorhinal circuit is differently organized along the dorsoventral axis in rodents. Cell Rep 42, 112001 (2023).

67. T. Butola et al., Hippocampus shapes entorhinal cortical output through a direct feedback circuit. Nat Neurosci 28, 811–822 (2025).

68. S. Craig, S. Commins, Plastic and metaplastic changes in the CA1 and subicular projections to the entorhinal cortex. Brain Res 1147, 124–139 (2007).

69. G. Nagel et al., Light activation of channelrhodopsin-2 in excitable cells of Caenorhabditis elegans triggers rapid behavioral responses. Curr Biol 15, 2279–2284 (2005).

70. T. van Haeften, L. Baks-te-Bulte, P. H. Goede, F. G. Wouterlood, M. P. Witter, Morphological and numerical analysis of synaptic interactions between neurons in deep and superficial layers of the entorhinal cortex of the rat. Hippocampus 13, 943–952 (2003).

71. C. V. Vorhees, M. T. Williams, Assessing spatial learning and memory in rodents. ILAR J 55, 310–332 (2014).

72. C. V. Vorhees, M. T. Williams, Morris water maze: procedures for assessing spatial and related forms of learning and memory. Nat Protoc 1, 848–858 (2006).

73. E. F. Willis, P. F. Bartlett, J. Vukovic, Protocol for Short- and Longer-term Spatial Learning and Memory in Mice. Front Behav Neurosci 11, 197 (2017).

74. F. Carpenter, D. Manson, K. Jeffery, N. Burgess, C. Barry, Grid cells form a global representation of connected environments. Curr Biol 25, 1176–1182 (2015).

75. C. Grienberger, J. C. Magee, Entorhinal cortex directs learning-related changes in CA1 representations. Nature 611, 554–562 (2022).

76. J. B. Priestley, J. C. Bowler, S. V. Rolotti, S. Fusi, A. Losonczy, Signatures of rapid plasticity in hippocampal CA1 representations during novel experiences. Neuron, (2022).

77. A. D. Madar, A. Jiang, C. Dong, M. E. J. Sheffield, Synaptic plasticity rules driving representational shifting in the hippocampus. Nature Neuroscience, (2025).

78. W. Gerstner, M. Lehmann, V. Liakoni, D. Corneil, J. Brea, Eligibility Traces and Plasticity on Behavioral Time Scales: Experimental Support of NeoHebbian Three-Factor Learning Rules. Front Neural Circuits 12, 53 (2018).

79. C. Le Duigou, J. Simonnet, M. T. Telenczuk, D. Fricker, R. Miles, Recurrent synapses and circuits in the CA3 region of the hippocampus: an associative network. Front Cell Neurosci 7, 262 (2014).

80. N. L. M. Cappaert, Van Strien, N. M. & Witter, M. P., The Rat Nervous System. Chapter 20 — Hippocampal formation (ed. Fourth Edition, 2015).

81. Y. Yoshiyama et al., Synapse loss and microglial activation precede tangles in a P301S tauopathy mouse model. Neuron 53, 337–351 (2007).

82. R. J. Low, Y. Gu, D. W. Tank, Cellular resolution optical access to brain regions in fissures: Imaging medial prefrontal cortex and grid cells in entorhinal cortex. Proceedings of the National Academy of Sciences 111, 18739–18744 (2014).

83. D. Aronov, David W. Tank, Engagement of Neural Circuits Underlying 2D Spatial Navigation in a Rodent Virtual Reality System. Neuron 84, 442–456 (2014).

84. E. A. Mukamel, A. Nimmerjahn, M. J. Schnitzer, Automated Analysis of Cellular Signals from Large-Scale Calcium Imaging Data. Neuron 63, 747–760 (2009).

85. M. Pachitariu et al., Suite2p: beyond 10,000 neurons with standard two-photon microscopy. bioRxiv, 061507 (2017).

86. James G. Heys, Krsna V. Rangarajan, Daniel A. Dombeck, The Functional Micro-organization of Grid Cells Revealed by Cellular-Resolution Imaging. Neuron 84, 1079–1090 (2014).

87. L. Sheintuch et al., Tracking the Same Neurons across Multiple Days in Ca2+ Imaging Data. Cell Reports 21, 1102–1115 (2017).

88. C. Domnisoru, A. A. Kinkhabwala, D. W. Tank, Membrane potential dynamics of grid cells. Nature 495, 199–204 (2013).

89. J. Tapson, A. van Schaik, Learning the pseudoinverse solution to network weights. Neural Networks 45, 94–100 (2013).

90. J. Larson, D. Wong, G. Lynch, Patterned stimulation at the theta frequency is optimal for the induction of hippocampal long-term potentiation. Brain Research 368, 347–350 (1986).

